# Stimulation strategy matters: A B-cell focused single-cell transcriptomic study on *in vitro* stimulated immune cells

**DOI:** 10.64898/2026.10.22.760934

**Authors:** D’hamers Tine, Neirinck Jana, Malicorne Buysse, Vriendt De Ciel, Kerre Tessa, Haerynck Filomeen, Hofmans Mattias, Bonroy Carolien, Everaert Celine

## Abstract

Distinct *in vitro* B-cell stimulation strategies, defined by stimulation condition and incubation duration, have been described in diverse studies. Often, these approaches are considered interchangeable due to comparable flow-cytometric outputs concerning fluorescence intensities and B-cell subset distributions. However, whether these phenotypic similarities also reflect comparable underlying transcriptional states have not been systematically evaluated yet. In this study, we provide an unbiased comparison of B-cell activation strategies on healthy individuals, resulting in a single-cell transcriptomic atlas (4 donors; ∼2,029 cells per sample; 81,160 cells in total with 56.3% annotated as B cells). Two T-cell independent stimulators targeting toll-like receptors (ODN2006 and IFN-γ/R-848), were assessed next to two T-cell dependent stimulators (anti-CD40 and anti-CD3/CD28 antibodies), all incubated for eight and 30 hours. Observed overlap for differential abundance, differential gene expression and cell-cell interactions could directly be linked to T-cell (in)dependent working mechanism. Strategy-specific results uncovered time-point specific transcriptional changes, revealed the largest set of differentially expressed genes for IFN-γ/R-848 and ODN2006, and showed that IFN-γ/R-848 and anti-CD3/CD28 induced the majority of the cell-cell interactions. Ultimately, our reference atlas emphasizes the importance and facilitates the rational selection of optimal B-cell stimulation strategies, particularly for studies aiming to interrogate B-cell dysregulation in immune-mediated diseases.

**GRAPHICAL ABSTRACT:** 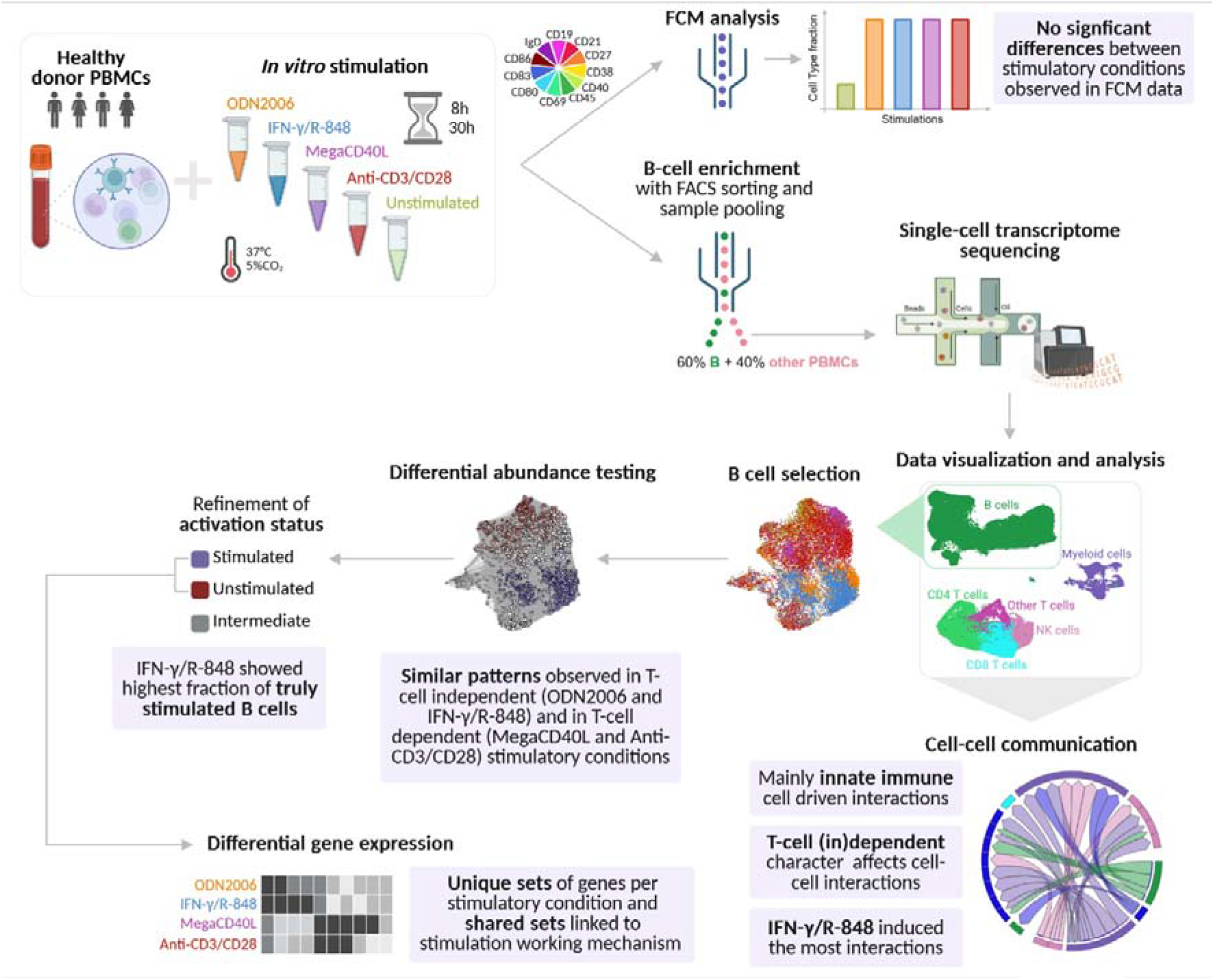

## INTRODUCTION

The humoral immune response protects the body against extracellular threats through antibody production, which depends on the proper differentiation of naive B cells into antibody-secreting cells and memory B cells^1,2^. B-cell activation involves antigen binding to the B-cell receptor (BCR) and secondary signalling from either T-cell dependent or independent pathways^3,4^. T-cell dependent responses, induced by protein antigens, require follicular helper T (Tfh) cells, involving cytokines like interleukin-4 (IL-4) and IL-21, and CD40 ligand (CD40L) interactions^5^. T-cell independent responses, induced by non-protein antigens, such as bacterial lipopolysaccharides, activate B cells often through toll-like receptors (TLRs)^5^. Downstream B-cell processes are regulated by intracellular pathways, such as PI3K/AKT and NF-κB, affecting expression of specific genes as a response to the activation stimulus^2,4,6–10^.

Various conditions have been used to *in vitro* stimulate B cells, focusing on distinct stimulation mechanisms and incubation periods ^11–13^. Commonly used direct B-cell stimulators comprise antibodies targeting CD40L and anti-CD40, mimicking T-cell dependent B-cell response pathways, or targeting immunoglobulins (e.g. IgG, IgA, IgM) to promote BCR aggregation^11,12,14^. TLR agonists, such as CpG-oligodeoxynucleotides (ODN) and resiquimod (R-848), targeting TLR9 and TLR7/8 respectively, are used for *in vitro* B-cell stimulation by mimicking T-cell independent B-cell responses^11,12,15^. Indirect stimulation via T-cell activation with anti-CD3/CD28 is also common^13^. To enhance B-cell proliferation, immunoglobulin production, and cytokine secretion, stimulators are often combined with cytokines like IL-4, IL-21, or interferon-γ (IFN-γ)^11,12^. Next to distinct stimulators, the incubation time forms another important aspect of the stimulation strategy, generally ranging from 4h to multiple days^11–13,16–20^. Despite their extensive characterization and application, the diversity of B-cell stimulation strategies makes it challenging to compare results across studies.

Different studies have been conducted to understand the impact of *in vitro* B-cell stimulation on the immunophenotype, B-cell subset decomposition and the molecular level, which traditionally focus on profiling of the stimulated B-cell compartment using flow cytometry (FCM), targeted RNA expression, and immunoglobulin secretion^11,12,21,22^. Contrastingly, unbiased single-cell analysis techniques are increasingly being used to explore *in vitro* immune responses^22–24^. Single-cell RNA sequencing (scRNA-seq) provides high-throughput, genome wide transcriptome profiling at single-cell level, enabling in-depth characterization of individual cells while capturing the heterogeneity of complex samples^25^. For example, Lawlor *et al.* (2021) has used an expanded scRNA-seq approach to profile the effect of lipopolysaccharides (LPS) and anti-CD3/CD28 stimulation on the different immune cell subsets with a focus on monocytes, B, NK and T cells^13^. Recently, another study conducted by Cui *et al.* (2024) revealed that *in vitro* stimulation with different cytokines induced highly cell-type and stimulation specific single-cell transcriptomic profiles in mouse lymph nodes^26^. These studies highlight the importance of understanding the differences in the mechanism of action and effect of the different stimulation approaches on the immune cell compartment, especially valuable in selecting the most suitable approach for specific research or clinical purposes.

Previous studies have shown that different *in vitro* B-cell stimulation approaches induce comparable FCM cell subset distributions and/or increased fluorescence intensities of activation markers^11,12^. However, whether such phenotypic similarity reflects comparable underlying transcriptional states has not been systematically evaluated. Furthermore, B-cell focused studies investigating differences in single-cell transcriptomic profiles between T-cell independent and dependent B-cell stimulators, are currently lacking. This might enable capturing differences between cell (sub)types of interest that might be missed at protein or bulk transcriptomic levels. Taking into account the potential diversity in effects of stimulation strategies is of utmost importance in selecting the appropriate stimulator, especially in studies focusing on specific impaired cellular or molecular functions causal for (immune-related) diseases.

Therefore, in this study, we generated an unbiased, sample multiplexed scRNA-seq atlas to explore the impact of distinct *in vitro* B-cell stimulatory conditions (including cytokines, CD40L, anti-CD3/CD28, ODN, and R-848) and incubation durations (eight and 30 hours). We focused on targeted B-cell specific pathways and interactions between immune cell (sub)types in B-cell enriched human peripheral blood mononuclear cell (PBMC) samples. This finally led to the identification of stimulation-specific patterns and differences between stimulatory strategies, guiding future research that uses *in vitro* B-cell stimulation.

## RESULTS

PBMCs isolated from healthy donors (n=4; two females and two males; 27–49 years) were incubated for 8h or 30h with ODN2006, IFN-γ/R-848, MegaCD40L, or anti-CD3/CD28 (**Figure 1**, **Sup. Table S1**) and compared to unstimulated samples to assess the effects on cellular activation and lineage differentiation using multiparameter FCM and scRNA-seq. To facilitate B-cell focused profiling, samples were enriched for B cells to a proportion of up to 60%. The generated data was computationally preprocessed and analyzed, enabling in-depth understanding of (dis)similarities between stimulatory strategies (**Sup. Table S1**). Detailed description of the experimental and computational analysis steps is provided in the Materials and Method section.

**Figure 1:**
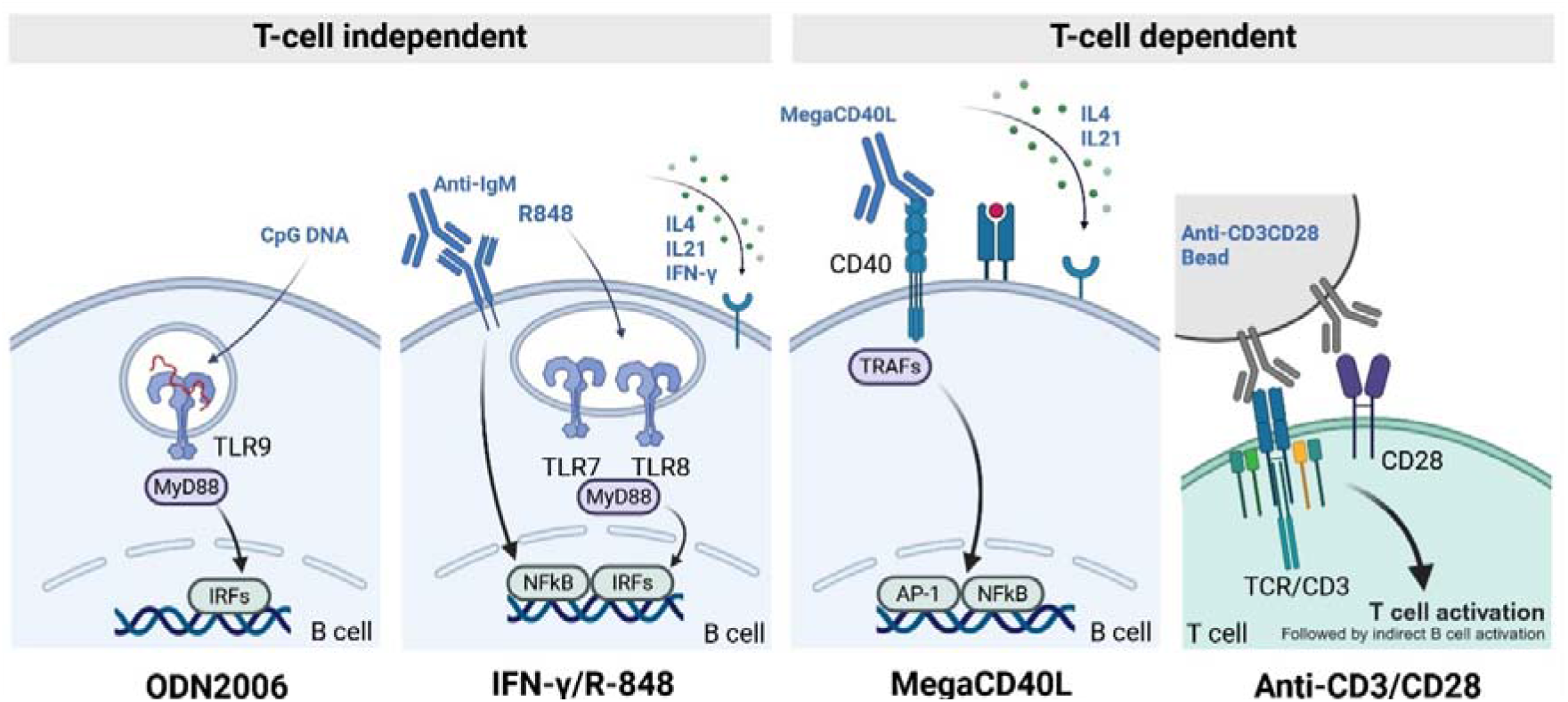
Overview of stimulatory conditions and their (intracellular) working mechanisms. Two T-cell independent (ODN2006 and IFN-γ/R-848; left) next to two T-cell dependent (MegaCD40L and anti-CD3/CD28; right) were investigated in this study. TLR, toll-like receptor; IRFs, interferon regulatory factors; IL, interleukin; TCR, T-cell receptor. Created with BioRender.com.

### Flow cytometry analysis did not reveal substantial differences between B-cell subsets across stimulatory strategies

The cell surface proteins, immune cell type decomposition and activation status of the B-cell compartment within (un)stimulated samples were examined using multiparameter FCM (**Sup. Table S2**). The flow panel targeted B-cell lineage and maturation markers (i.e. CD19, CD21, CD27, CD38, CD40 and IgD), inspired by the standardized EuroFlow Consortium panels, and activation markers (e.g., CD69, CD83, CD86) ^12,57–59^. Following data pre-processing, a total of 1.33 million CD45^+^CD19^+^ B cells (over 4 donors) were analysed and split between cells incubated for 8h (665,160 cells; 50% of total B cells) and 30h (664,311 cells; 50% of total B cells). B cells were clustered and annotated based on expression patterns of the targeted markers (**Figure 2A** and **Sup. Figure S1**), resulting in three B-cell subsets: pre-germinal center (CD27□IgD□), unswitched memory (CD27□IgD□), and switched memory B cells (CD27□IgD□). B-cell activation status was determined on increased CD69 expression. The proportions of the CD69^-^ (inactivated) and CD69^+^ (activated) subsets per sample were compared across different stimulatory conditions (**Figure 2A** and **Sup. Figure S1**).

**Figure 2:**
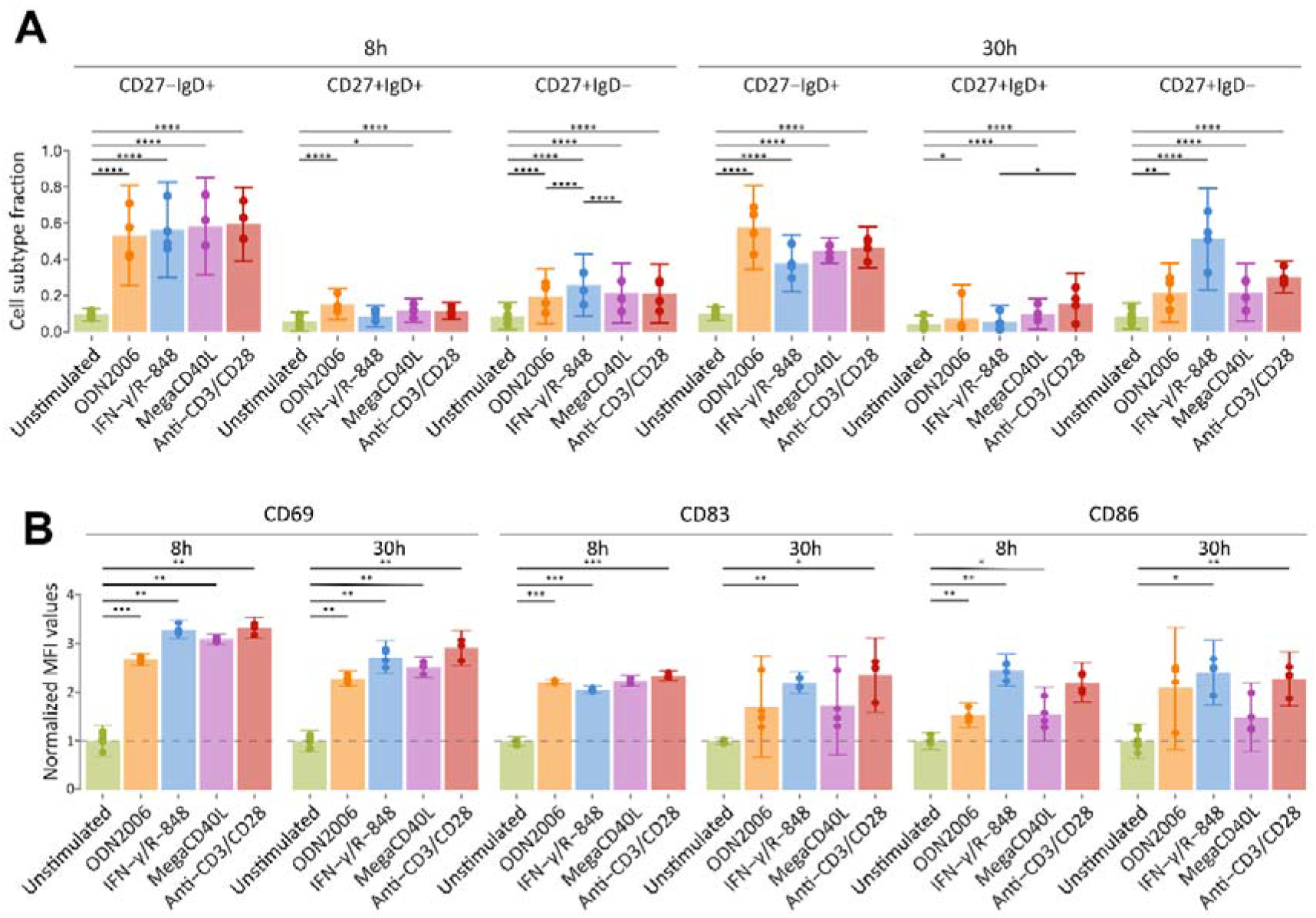
Flow cytometric analysis of *in vitro* stimulated PBMCs. (**A**) The fraction of activated (CD69^+^) B cells (CD19^+^) per healthy donor, stimulation condition, and incubation time point across B cell subsets. CD27^-^ IgD^+^ population represent pre-germinal center B cells. Unswitched and switched memory B cells are characterized by CD27^+^IgD^+^ and CD27^+^IgD^-^ phenotypes, respectively. Error bars representing 95.5% (mean ± 2*standard deviation) confidence intervals. Dots represent individual cell type fractions per healthy donor. Significant differences are indicated, ****p<0.0001, ***p<0.001, **p<0.01, *p<0.05, testing based on generalized linear mixed model, beta-family regression using a logic link function. (**B)** Bar plots of mean fluorescence intensities (MFI) of activation markers, i.e. CD69, CD83, CD86, within the CD45+ subset, normalized with respect to unstimulated samples. Error bars representing 95.5% (mean ± 2*standard deviation) confidence intervals. Dots represent individual measurements per healthy donor. Significant differences with respect to control samples are shown, ***p<0.001, **p<0.01, *p<0.05, testing with paired t-test.

Incubation with all distinct stimulatory conditions resulted in successful B-cell activation with significant increases in fraction of CD69□ activated cells compared to unstimulated controls (p_adj_<0.01), except for IFN-γ/R-848 stimulated CD27^+^IgD^+^ cells, both after 8h (|Δ_fraction_|=0.015) and 30h (|Δ_fraction_|=0.009; **Figure 2A**). The latter is possibly linked to promotion of class switching in memory B cells and induction of antigen secreting cells ^16,17^. Among the stimulatory conditions, most activated subsets showed no significant differences in proportions at either time point, with two exceptions: IFN-γ/R-848 stimulation led to a significant increase in activated switched memory B cells compared to ODN2006 (|Δ_fraction_|=0.057) and MegaCD40L (|Δ_fraction_|=0.0456) after 8h (both p_adj_<0.0001), and a decrease in activated unswitched memory B cells compared to anti-CD3/CD28 after 30h (|Δ_fraction_|=0.101; p_adj_<0.05). In addition, activation markers CD69, CD86, and CD83 exhibited comparable increases in fluorescence intensity, with average normalized values ranging from 1.49 to 3.32, across all stimulated samples relative to unstimulated controls (**Figure 2B**). Although these findings confirm successful B-cell activation under distinct stimulation conditions, they also underscore the limitations of the selected FCM markers for detailed analysis of stimulation-specific effects on (subsets) of B-cells.

### Successful B-cell enrichment uncovers stimulation strategy-driven clustering

scRNA-seq enabled in-depth and more unbiased profiling of the (un)stimulated immune cells, with the main focus on the enriched B-cell subset. Following library preparation, sequencing, data pre-processing and immune type subset annotation, a total of 43,100 B cells (46.8% of 8h stimulated cells; 53.2% of 30h stimulated cells), 15,009 CD4^+^ T cells, 9,382 CD8^+^ T cells, 1,803 other T cells, 3,901 NK cells and 5,389 myeloid cells were identified based on cell-type specific gene expression levels (**Sup. Figure S4B-C**). The B-cell subsets for 8h and 30h of stimulation were filtered out separately for downstream analysis, including Uniform Manifold Approximation and Projection (UMAP) visualisations and discrete, conventional Louvain cell clustering (**Figure 3A-D**). When correlating the within-cluster cell fractions of the stimulatory conditions, high positive Spearman correlations between MegaCD40L and anti-CD3/CD28 (0.806 at 8h; 0.757 at 30h), and between IFN-γ/R-848 and ODN2006 (0.879 at 8h; 0.280 at 30h) were observed. This suggests that cells tend to cluster according to stimulation strategy characteristics (T-cell (in)dependent) rather than maturation status (**Figure 3A-F** and **Sup. Figure S7**).

**Figure 3:**
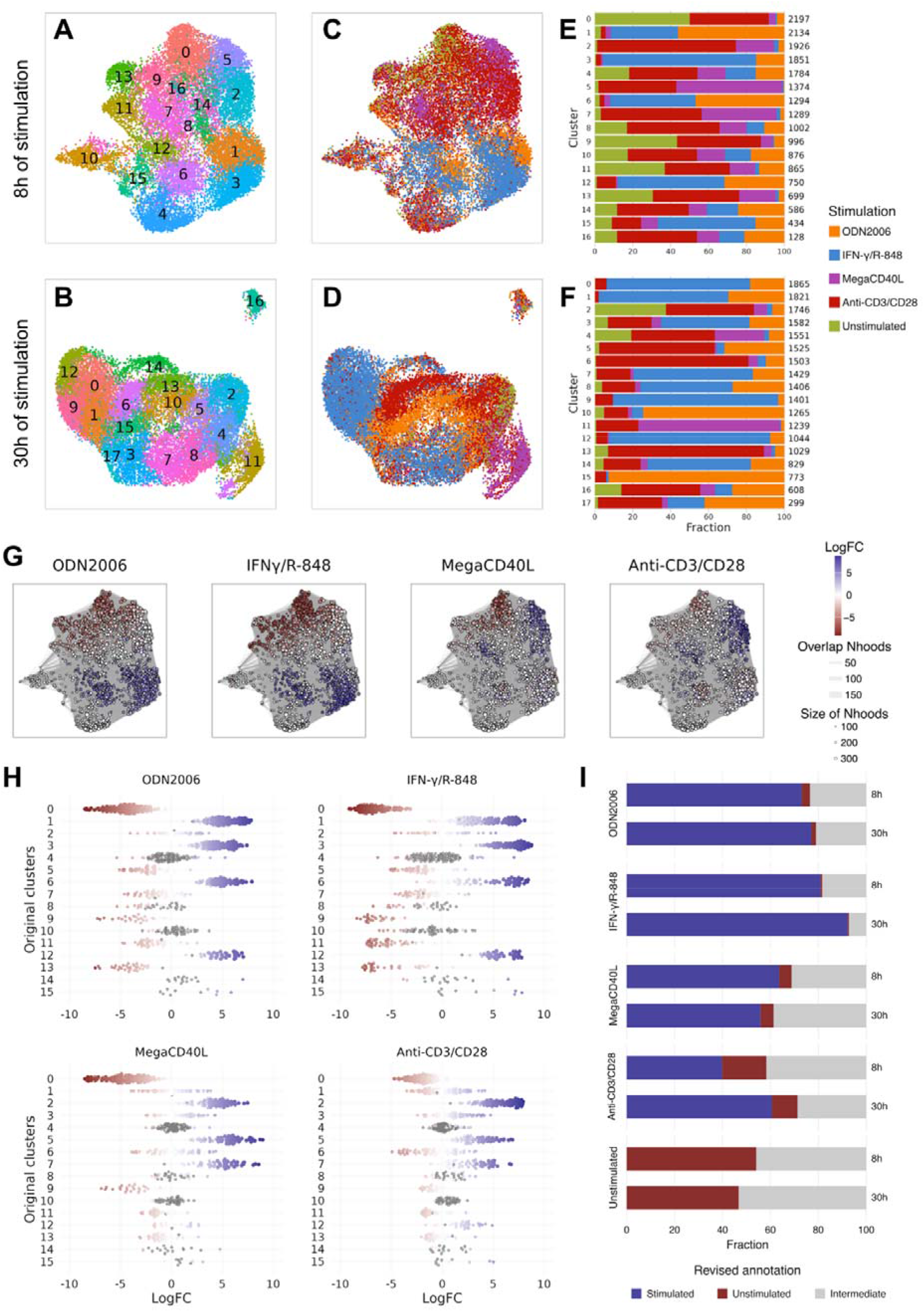
Clustering, differential abundance testing and refinement of the stimulatory status within the B-cell subset. (**A-D**) The B-cell subset after 8h (20,185 cells) and 30h (22,915 cells) of stimulation visualized in a two-dimensional UMAP, colored based on the obtained conventional Louvain cell clusters (**A, B**) or the stimulatory condition at sample-level (**C, D**). (**E, F**) Bar plots showing stimulation distributions across the conventional Louvain clusters for each of the incubation time points separately, with the total number of cells per cluster shown at the right-hand side of each bar. (**G**) Neighbourhood (Nhood) graph based on the UMAP visualization of the cells stimulated for 8h per stimulatory condition, show the results of the differential abundance testing with unstimulated cells as reference using MiloR. Each Nhood is represented as a dot of which the size is correlated to the number of cells. The Nhoods are colored based on LogFC values of differential abundance testing: red indicates increased abundance of unstimulated cells, blue indicates increased abundance of stimulated cells and white indicate no significant differences in abundance. (**H**) Beeswarm plots for eight hours of stimulation showing the distribution of LogFC after abundance testing per Nhood (dots) per Louvain cell cluster (y-axis), performed for each of the four stimulations separately. Full colored dots represent the Nhoods selected for downstream analysis, same coloring scheme was used as described in panel **G**. (**I**) Comparison of the fractions of the refined activation status annotation per stimulatory condition and incubation time point.

### Differential abundance testing hints towards B-cell transcriptional patterns linked to stimulatory working mechanism

The rigid conventional clustering approach, based on stringent cutoff definitions, results in heterogeneous clusters (**Figure 3E-F**). Therefore, unambiguous annotation and prioritization of interesting cell subsets enriched for specific (combinations of) conditions is difficult. To overcome these limitations, MiloR was used to identify overlapping regions of transcriptomically similar cells, referred to as neighbourhoods (Nhoods) ^48^. These Nhoods were subsequently subjected to differential abundance testing (**Figure 3G** and **Sup. Figure S5A**), comparing each of the stimulated cells to the unstimulated reference cells. This pinpoints Nhoods predominantly composed of unstimulated cells, and other Nhoods strongly enriched for stimulated cells, clearly delineating stimulated and unstimulated regions projected on the UMAP (Figure 3**G**) ^48^. Interestingly, differential abundant Nhood patterns induced by TLR agonists (IFN-γ/R-848 and ODN2006) were highly similar to each other, as were those induced by MegaCD40L and anti-CD3/CD28 (**Figure 3G-H** and **Sup**. **Figure S5A**). These findings are in line with the cluster-based observations, and confirm the presence of stimulus-specific transcriptional programs in B cells, highly related to the working mechanisms and targeted pathways for each of the stimulatory conditions.

### Annotation refinement reveals a substantial fraction of unstimulated cells in MegaCD40L and anti-CD3/CD28

Next to similar patterns between stimulatory conditions, the differential abundance testing also revealed higher abundant unstimulated regions when focusing on MegaCD40L and anti-CD3/CD28 stimulated cells, after both 8h and 30h of stimulation (**Figure 3G** and **Sup**. **Figure S5A**). This suggests a lower stimulation performance compared to the TLR agonists IFN-γ/R-848 and ODN2006. To evaluate the truly stimulated cell fraction, the annotation of the stimulation status of the cells was refined, for both incubation time points separately. This involved a two-step selection procedure of the Nhoods, as described in the Materials and Method section. Selected Nhoods with positive differential abundance values were annotated as ‘truly stimulated’, while negative differential abundance values were linked to ‘truly unstimulated’ cells (**Figure 3I** and **Sup**. **Figure S5B**). Remaining cells that could not be linked to ‘truly stimulated’ or ‘truly unstimulated’ Nhoods, were annotated as ‘intermediate’ (white Nhoods in **Figure 3G**) and consisted of 32.9% (6,631 cells; 8h) and 21.9% (5,019 cells; 30h) of the total B cell subsets (**Figure 3**I and **Sup. Figure S6).** These cells are believed to reflect a transitional state between unstimulated and stimulated status and were not considered for downstream analysis.

Annotation refinement of the stimulation status leads to an average of 39.9%, 73.1%, 81.0% and 63.7% of ‘truly stimulated’ cells in anti-CD3/CD28, ODN2006, IFN-γ/R-848 and MegaCD40L conditions, respectively (8h). After 30h of stimulation, similar results were observed for ODN2006 (77.1%) while a significant increase was found in IFN-γ/R-848 stimulated samples (92.4%, p_adj_<0.001). The average stimulated fraction for MegaCD40L stimulated samples decreased to 55.9% (p_adj_>0.05). For anti-CD3/CD28, this fraction significantly increased up to 60.7% (p_adj_<0.001) when samples were incubated for 30h, indicating that stimulation for a longer time period is beneficial for the activity performance of anti-CD3/CD28. This condition also showed the highest fraction of unstimulated cells (18.5% and 10.7% after 8h and 30h, respectively) in comparison to the other conditions (<6%; p_adj_<0.001 except for ODN2006 30h (p_adj_=0.029) and MegaCD40L 30h (p_adj_>0.05)) and can be explained by the required B- and T-cell interaction of this stimulatory condition. The smallest fraction of unstimulated cells in stimulated samples was observed in the IFN-γ/R-848 condition (0.7% and 0.5% after 8h and 30h, respectively; significant differences p_adj_<0.04 compared to all other conditions), implying that the majority of the B cells within these samples differed substantially from the unstimulated B cells, indicating high activity performance.

Focusing on IFN-γ/R-848 and ODN2006 separately, the same regions and cell densities were observed on the UMAP visualization when comparing the original metadata-based to the refined Nhood-based annotation projected (**Sup. Figure S8**). This shows that the majority of the regions linked to these stimulatory conditions are retained after refinement, underlining the observation of a high stimulated sample fraction. In the MegaCD40L and anti-CD3/CD28 stimulated samples, only the densest regions of the metadata-based annotation were retained as truly stimulated after annotation refinement (**Sup. Figure S8**). Enrichment for truly stimulated cells thus reduces the confounding of unstimulated and intermediate cells and improves the performance of downstream analysis steps.

### Differential gene expression indicates overlap and uniqueness related to conditions and incubation duration

The refined stimulation status was used to identify differentially expressed genes (DEGs) in stimulatory conditions with unstimulated cells as a reference for both incubation times separately. This analysis resulted for IFN-γ/R-848, ODN2006, anti-CD3/CD28 and MegaCD40L in 1,060, 629, 318 and 178 DEGs after 8h, and 455, 258, 213 and 152 DEGs after 30h, respectively (**Figure 4A-B, Sup. Material D1** and **D2**). The overlap between all four stimulatory conditions revealed 74 (20 up-and 54 downregulated; 8h) and 31 DEGs (14 up- and 17 downregulated; 30h), including genes related to immune activation (e.g. *CXCR4*, *CD27*, *CCR6*, *CD79B*, *TNFRSF4*), cell proliferation (e.g. *CDC45*) and stress response (e.g. *GPAT2, ZFP36L1*).

**Figure 4:**
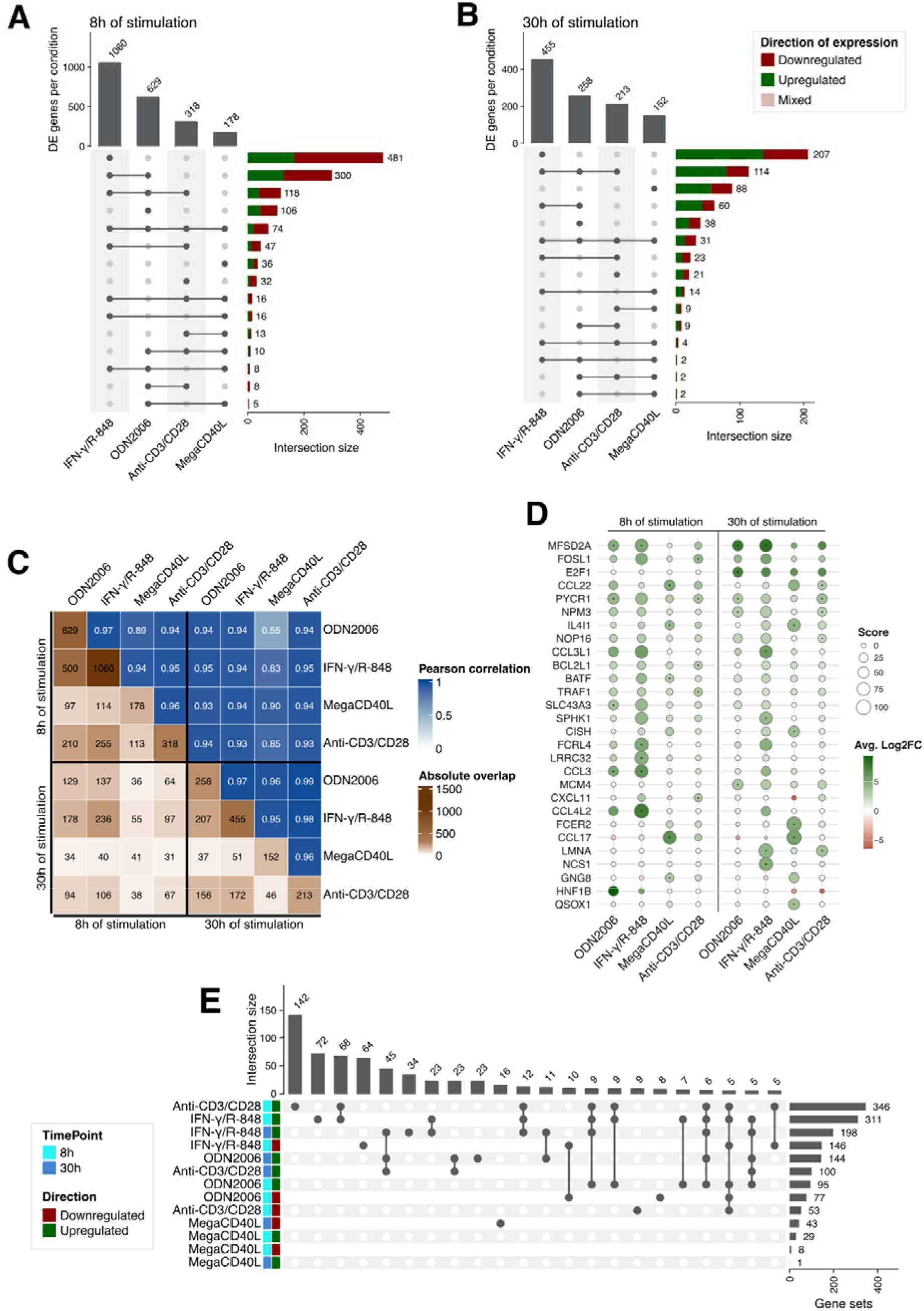
Differential gene expression per condition and incubation time point. (**A-B**) Upset plot of differentially expressed (DE) genes for the four conditions compared to the unstimulated cells for 8h (**A**) and 30h (**B**) of stimulation. The vertical bar plot on top visualizes the total number of significant DE genes (p_adj_<0.05) per stimulatory condition. The horizontal bar plot indicates the size of the overlap in DE genes for the indicated combination of stimulatory conditions and are colored based on consistency in expression direction (“Downregulated” for log2FC<-2; “Upregulated” for log2FC>2). If discrepancies in expression direction between conditions is observed, the label “Mixed” is used. (**C**) A two triangle heatmap visualizing the absolute overlap in detected significantly expressed genes between stimulatory conditions and incubation time points (lower triangle), and the Pearson correlation levels of the expression values (log2FC) of these overlapping genes (upper triangle). (**D**) Gene expression values (log2FC-based coloring) of the top five upregulated genes per stimulatory condition and incubation time point. The size of the dots represents the absolute gene score (see Materials&Methods section for detailed information). The presence of a small dot in the center of the larger dots, indicates that the gene is one of the top five DE genes in that particular condition. (**E**) Upset plot representing the overlap in significantly enriched Gene Ontology (GO) gene sets between stimulatory conditions and incubation time points. The horizontal bar plot represents the absolute number of significantly enriched GO gene sets per stimulatory condition split by direction of regulation. The vertical bar plot represents the size of the overlap in significantly enriched GO gene sets.

Unique sets of DEGs were found for each of the single stimulation mixes. The largest unique set of DEGs was observed for IFN-γ/R-848 stimulation, consisting of 481 (35% up-, 65% downregulated) after 8h and 207 genes (66% up-, 34% downregulated) after 30h, highlighting a large diversity of genes affected by IFN-γ/R-848 in comparison to other stimulatory conditions. The top five unique upregulated DEGs for this condition highlighted *HSPA1B*, *RSAD2*, *CD274* (*PD-L1*), *ID2* and *PDE4DIP* after 8h, and *NCS1*, *SPHK1*, *DUSP4*, *ZBTB32* and *ID2* after 30h of stimulation (**Sup. Figure S9C, Sup. Material D1** and **D2**). Only *ID2* was found at the two timepoints in the top five upregulated genes. *ID2* is an intrinsic negative regulator of B-cell development, suggesting a shift towards a more mature B-cell state is blocked^60,61^. Focusing on the top five downregulated genes for IFN-γ/R-848, *FTH1*, *CD37*, *JUND*, *SH3BGRL3* and *PDE4D* were observed after 8h, and *RCSD1*, *CD79B*, *ANKRD12*, *PDE7A* and *SEL1L3* after 30h of stimulation (**Sup. Figure S9C, Sup. Material D1** and **D2)**. Various genes associated to the cAMP-related (3’,5’-cyclic adenosine monophosphate) adaptive immune response, e.g. *PDE4D*, *PDE7A*, were observed as downregulated upon IFN-γ/R-848 stimulation. These genes are extensively described in activated T cells, but also linked to B-cell malignancies, such as diffuse large B-cell lymphoma (DLBCL)^62–64^. A link with DLBCL is also described for deficiencies in the cell surface protein CD37, predominantly expressed in B cells^65–67^. Knock outs of *JUND*, encoding for a protein involved in the AP-1 complex, were found to be strongly involved in apoptotic death and proliferative defects in response to stress^68,69^. Next to IFN-γ/R-848, the other conditions also showed unique sets of DEGs. In ODN2006 stimulated samples, this resulted in a set of 106 (42% up-, 58% downregulated) after 8h and 38 unique genes (55% up-, 45% downregulated) after 30h. For MegaCD40L 36 (8h; 56% up-, 44% downregulated) and 88 genes (30h; 63.5% up-, 37.5% downregulated) were found to be unique for this condition. The smallest set of unique DEGs was linked to anti-CD3/CD28 stimulation, resulting in only 32 (25% up-, 75% downregulated) and 21 unique DEGs (57% up-, 43% downregulated) for 8h and 30h, respectively (**Figure 4A-B**). The top five up- and downregulated genes of these unique subset are highlighted in **Sup. Figure S9 (Sup. Material D1** and **D2)**.

After 8h of stimulation, the overlap in DEG between T-cell independent stimulatory conditions IFN-γ/R-848 and ODN2006 (300 DEGs) is larger than after 30h (60 DEGs) (**Figure 4A-B**). Interestingly, *MFSD2A* (both time points) and *CCL3* (only in 8h) are common genes present in the top five upregulated genes of both IFN-γ/R-848 and ODN2006 stimulated samples (**Figure 4D)**. *MFSD2A* is mainly described in blood-brain barrier focused research and linked back to activated T cells, but is lacking in B-cell focused research^70,71^. *CCL3* is a well-known chemokine linked to B-cell receptor activation and DLBCL^72^. 74 DEGs after 8h and 31 DEGs after 30h were shared between all four conditions. Exclusion of MegaCD40L showed that the number of genes overlapping between IFN-γ/R-848, ODN2006 and anti-CD3/CD28 was stable comparing 8h (118 DEGs) and 30h (114 DEGs), however, only 16 genes were consistently expressed at both time points (**Figure 4A-C**). The decrease in shared genes between all four conditions when moving from 8h to 30h of incubation, and the consistency in shared genes between ODN2006, IFN-γ/R-848 and anti-CD3/CD28, implies divergence of MegaCD40L from the other stimulatory conditions when samples are stimulated longer. This is also confirmed by the increasing number of, mainly upregulated, DEGs uniquely associated with 30h compared to 8h of MegaCD40L: 36 to 88 DEGs (**Figure 4A-B**, **Sup. Figure S9A-B**).

### Concordance of expression direction among shared DEGs between conditions

As expected, the direction of expression (upregulation for positive log2FC and downregulation for negative log2FC) was consistent along the stimulatory conditions for both time points, except for three genes: *IFIT1, PPP1R14A* and *TMEM273* (**Figure 4A-B, Sup. Figure S9**). *IFIT1,* encoding for an anti-viral protein, was found to be upregulated (p<0.0001) after stimulation with ODN2006 (log2FC=2.17) and downregulated when stimulated with MegaCD40L (log2FC= -2.92) and anti-CD3/CD28 (log2FC= -2.82), after 8h. This is in line with Kato *et al.* (2003), who reported increased expression levels of *IFIT1* after 6h and 24h of stimulation with ODN2006 ^73^. The downregulation of the interferon-induced *IFIT1* gene in the MegaCD40L stimulated samples might be linked to the presence of IL-4 in this stimulation mix, a cytokine that is known to inhibit IFN-γ production in immune cells^74^. *PPP1R14A*, encoding for phosphatase inhibitor protein, and *TMEM273,* encoding for transmembrane protein, were both upregulated (p_adj_<0.0001) in MegaCD40L-stimulated samples (log2FC_PPP1R14A_=2.61; log2FC_TMEM273_=2.44), but downregulated after stimulation with ODN2006 (log2FC_PPP1R14A_= -3.63; log2FC_TMEM273_= -2.16), after 8h. This again highlights that, if differences are observed, it can be linked back to the T-cell independent and dependent nature of the stimulatory conditions. No DEGs with inconsistent expression direction were observed after 30h of stimulation. When comparing conditions and incubation time points side by side, the consistency of gene expression is retained as shown by the Pearson correlation scores based on the log2FC expression levels of the overlapping genes (**Figure 4C, Sup. Figure S10**).

### Enriched GO gene sets are uniquely linked to specific conditions and incubation time points

To gain insight into the functional roles of the DEGs across the conditions, Gene Ontology (GO) enrichment analysis focusing on biological processes (available at MSigDB^50^) was performed on the extracted sets of up- and downregulated DEGs per condition and time point separately. The majority of differential GO gene sets, both up- and downregulated, was uniquely linked to a specific condition and incubation time point (**Figure 4E**). A large number of differential GO gene sets were observed after anti-CD3/CD28 and IFN-γ/R-848 stimulation (8h): 346 and 311, respectively (**Figure 4E**). 142 and 72 GO gene sets were uniquely linked to upregulated DEGs after 8h of anti-CD3/CD28 and IFN-γ/R-848 stimulation, respectively, while 68 GO gene sets were shared between these two subsets. Furthermore, 45 of the identified GO gene sets were observed after 8h of IFN-γ/R-848, ODN2006 and anti-CD3/CD28 stimulation (upregulated DEGs only; **Figure 4E**). Most of the GO gene sets significantly enriched after MegaCD40L stimulation were not observed in other conditions. In general, GO gene sets identified after 8h of stimulation were mainly linked to immune system processes and response, and (immune) cell activation, while GO gene sets associated with 30h of stimulation were mainly linked to metabolic, cell cycle and transition/migration processes (**Sup. Figure S11**).

### IFN-**γ**/R-848 and anti-CD3/CD28 induce numerous cell-cell interactions, while targeting different cell types

The B-cell compartment was the primary focus of the previous analysis sections. However, PBMCs

contain other immune cells (NK, CD4^+^ T, CD8^+^ T and myeloid cells, **Sup. Figure S4B**) which are also present in approximately 40% of the total profiled cells in this scRNA-seq data set. Their presence is presumed to be importance in indirect B-cell stimulatory conditions, but nonetheless other immune cell interactions might occur upon or after *in vitro* stimulation, even in the direct conditions. Using MultiNicheNetR ^51^, a total of 5,311 and 4,746 interactions were inferred for 8h and 30h of stimulation, respectively. Ranking all of these interactions, based on the final score provided by MultiNicheNetR, resulted in an overall top 50 list dominated by IFN-γ/R-848 and anti-CD3/CD28 induced interactions (**Figure 5A**). Within this top 50, only three interactions were linked to MegaCD40L and two to ODN2006 stimulated samples (**Figure 5A**). Similar results were observed after 30h of stimulation (**Sup. Figure S12**). This indicates that IFN-γ/R-848 (probably cytokine driven) and anti-CD3/CD28 (indirect stimulator) indeed seem to have the strongest inductive effect on cell-cell interactions between immune cells, independent of the incubation duration.

**Figure 5:**
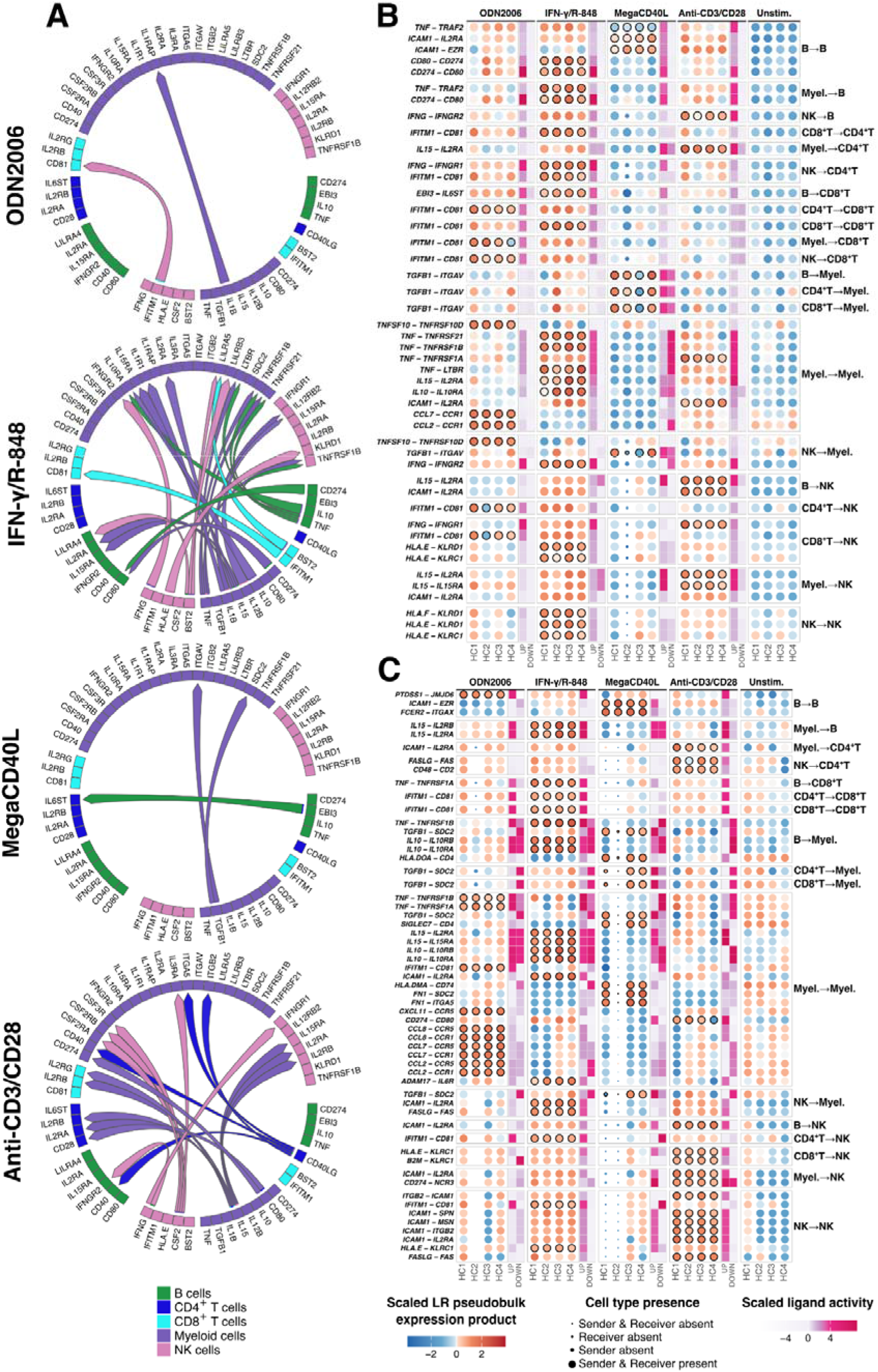
Cell-cell communication per stimulatory condition compared to the unstimulated samples after 8h and 30h of stimulation. (**A**) Chord diagrams showing the overall top 50 of highly ranked upregulated ligand-receptor pairs associated to the interacting cell types, split over stimulatory conditions (8h). The arrows, colored based on the sender cell type, connect the ligand in a specific sender cell type with the receptor in a specific receiver cell type. If arrows are absent between ligands and receptor, the combination of interaction and stimulatory condition was not present in the overall top 50 list for that specific condition. (**B-C**) Dot and tile plot combining the 46 and 57 prioritized interactions for 8h (**B**) and 30h (**C**) of incubation, respectively. The vertical axis represents ligand-receptor (LR) interactions at the left and cell type interaction at the right; sample conditions are indicated at the top. The dot plot shows a scaled (pseudobulk) expression of the LR combination per sample (HC1-HC4, bottom), while the dot size represents the presence or absence (after filtering, see Materials&Methods) of the sender and/or receiver cell type within the individual samples. The presence of a solid black stroke indicates prioritization of the interaction within that specific stimulatory condition. The tile plot shows the scaled ligand activity, indicating the enrichment of expected up- and downregulated target genes. Higher ligand activities (dark pink) suggests functional interaction affecting the downstream target genes in the receiver cell.

Focusing on the top 50 highest ranked interactions per stimulatory condition separately, revealed that the majority of interactions involved myeloid cells (e.g. dendritic cells and monocytes), either as sender, receiver or both (**Sup. Figure S13**-**S20**). For IFN-γ/R-848 and anti-CD3/CD28 (8h only) stimulated samples, interactions between B and myeloid cells mainly involved cytokines, such as IL-10, IL-15 and Tumor Necrosis Factor (TNF), while a variety of ligand/receptor pairs were observed in the other conditions. Beside myeloid driven interactions, anti-CD3/CD28 and MegaCD40L were associated with numerous interactions involving T and NK cells, mainly interacting with myeloid or B cells (**Figure 5B-C**, **Sup. Figure S15**, **S16**, **S19** and **S20**). Remarkably, the T-cell dependent interaction between CD40 and CD40L was observed in the highest ranked interactions after 8h of anti-CD3/CD28 stimulation (e.g. CD4^+^T:B), but lacking after 30h of incubation (**Sup. Figure S16, S20**). These interactions were absent in other conditions (**Sup. Figure S13**-**S20**), which might explain the similarities observed between anti-CD3/CD28 and MegaCD40L in the differential abundance and gene expression analysis after 8h of stimulation only.

### Expression direction of interaction-driven target genes differs between stimulation strategies

Remarkably, the stimulations did not only impact the ligand/receptor pairing between distinct cell types, but also induced differences in expression patterns of the downstream target genes linked to highly ranked cell-cell interactions. For example, IL-15 interacts with IL-2RA and IL-2RB in B:NK and Myeloid:NK communications and are highly ranked in samples stimulated for 8h with anti-CD3/CD28. Despite their absence in the top 50 ranking of other conditions, these interactions also seem to be present in IFN-γ/R-848 samples (high ligand-receptor pseudobulk expression) with a lower rank. Most remarking is the discrepancy in downstream gene expression pattern in the receiver cell: upregulation after anti-CD3/CD28 stimulation and downregulation after IFN-γ/R-848 stimulation. A similar observation is seen for TNF-involved interactions between B and myeloid cells at 8h of incubation, which induced upregulation in the receiver cell after ODN2006, IFN-γ/R-848, and anti-CD3/CD28 stimulation, while target genes are downregulated in MegaCD40L stimulated samples (**Figure 5B-C**, **Sup. Figure S13**-**S20**). After 30h, however, a switch to a downregulated pattern of these genes was also observed after anti-CD3/CD28 stimulation.

### In-depth prioritization of cell-cell interactions highlights unique, stimulation-induced communications

More in-depth prioritization of the cell-cell interactions was introduced by considering multiple criteria, including coherent up- or downregulation of target genes in the receiver cell, to highlight specific interactions that are more likely functionally active. For IFN-γ/R-848, anti-CD3/CD28, ODN2006 and MegaCD40L this lead to 20, 10, 9 and 7 prioritized interactions after 8h, and 19, 15, 11 and 12 prioritized interactions after 30h of stimulation, respectively (**Figure 5B-C**).

This prioritization approach highlighted CD274:CD80 in B:B communication as one of the top interactions in IFN-γ/R-848 (8h) stimulated samples (**Figure 5B**). Consistently, *CD274* or *PD-L1*, known as immune checkpoint, has also been identified in our atlas as one of the top five upregulated genes in B cells uniquely linked to IFN-γ/R-848 stimulation (**Sup. Figure S9**). In literature, CD274:CD80 interactions are mainly reported as cis (proteins present at the same cell membrane), while the trans-interaction (adjacent cells interacting) has been elusive^75,76^. However, CD274:CD80 was also inferred in the direct Myeloid:B trans-interaction with high activity (upregulation) of target genes of this ligand-receptor pair (IFN-γ/R-848; 8h), hinting towards the presence of possible trans-interactions in our samples. Furthermore, the interaction between IFITM1 (interferon-induced transmembrane protein) and CD81 was prioritized in both IFN-γ/R-848 and ODN2006 stimulated samples at both incubation points, mainly linked to interactions including CD4^+^/CD8^+^ T and/or NK cells and strong upregulation of downstream target genes (**Figure 5B-C**, **Sup. Figure S13**, **S14**, **S17** and **S18**). CD4^+^T:CD8^+^T and CD4^+^T:NK pairing via interaction between IFITM1 and CD81 specifically were consistently prioritized in 8h of ODN2006 and 30h of IFN-γ/R-848 stimulation, while being absent for T-cell dependent conditions (**Figure 5B-C, Sup. Figure S15**, **S16**, **S19** and **S20**). The latter observation is in line with strong downregulation of *IFITM1* observed in B cells following MegaCD40L (**Sup. Figure S9**). The IFITM protein family has been reported to have anti-viral activities by restricting the viral entry in many cells, which links back to R-848 and ODN2006 targeting TLR7/8 and TLR9, respectively^77,78^. This again underlines the highly similar characteristics and downstream intracellular effect of IFN-γ/R-848 and ODN2006, strongly differing from the T-cell dependent conditions.

Interaction involving monocyte chemotactic proteins (MCP) such as CCL2, CCL7 and CCL8 were prioritized only after ODN2006 stimulation (**Figure 5B-C**)^79,80^. Fleige and Capellino ^81^ reported an increase of CCL2 secretion following ODN2006 stimulation of PBMCs, but showed that B cells did neither produce, nor secrete this chemokine. Consistent with this observation, the inferred interactions were only restricted to Myeloid:Myeloid interactions, highlighting the secondary effects on the non-B cell immune compartment. Only two interactions (CCL7:CCR1 and CCL2:CCR1) were observed after 8h of stimulation, whereas four additional MCP-driven interactions were prioritized after 30h, suggesting that chemokine-mediated communication becomes more prominent over time (**Figure 5B-C**).

Zooming in on the interactions prioritized in anti-CD3/CD28 uncovered IFN-γ interactions with the receptors IFNGR1 (CD8^+^T:NK) and IFNGR2 (NK:B) after 8h of stimulation (**Figure 5B-C**). As (activated) T and NK cells are reported as the major IFN-γ producing cell population within the immune system, the inference of this interaction under anti-CD3/CD28 stimulation aligns with the direct T-cell activation character of this condition^82^. Since overlap between anti-CD3/CD28 and IFN-γ/R-848 in other analysis steps in this study was mainly observed after 30h of incubation, this interaction might indicate that pathways and interactions induced by IFN-γ/R-848 stimulation mimics the delayed IFN-γ production post anti-CD3/CD28 stimulation, finally leading to similar biological processes at different stimulation durations. Furthermore, interactions between intercellular cell adhesion molecule 1 (ICAM1) and diverse receptors were prioritized in anti-CD3/CD28 stimulated samples at both incubation times, confirming the importance of cell-cell interactions linked to this indirect stimulator^83,84^. It primarily involved IL2RA, referring to the alpha chain of IL-2 receptor, as a receptor including among others B:B, B:NK and NK:NK interactions (**Figure 5B-C**). While IL2RA (CD25) is mainly described in T-cell focused research, recent scientific evidence on interaction with ICAM1 between immune cells seems to be lacking^85–87^.

Finally, the majority of prioritized interactions linked to MegaCD40L stimulation after 8h of stimulation involved TGF-β1 as a ligand (**Figure 5B-C**). This cytokine is known to maintain the balance between cell proliferation and apoptosis, and is reported as a suppressor of CD40-mediated immune cell activation^88,89^. This might indicate that it tempers the activation of antigen presenting cells, such as myeloid and B cells, upon CD40-activation. This suppressive mechanism is less abundant in the prioritized interactions after 30h of stimulation with MegaCD40L, which might hint towards a delay in immune activation in MegaCD40L stimulated samples (**Figure 5B-C**, **Sup. Figure S15** and **S19**).

## DISCUSSION

In this study, we examined the effects of distinct *in vitro* B-cell stimulatory strategies on healthy PBMCs, including two T-cell independent (IFN-γ/R-848, ODN2006) and two T-cell dependent (MegaCD40L, anti-CD3/CD28) stimulatory conditions, using FCM and scRNA-seq. We focused on 8h and 30h of incubation, which are representable for the generally applied stimulation durations in literature ranging from a couple of hours to multiple days^11–13,16–19^. In line with recent studies on *in vitro* B cell stimulation, our FCM analysis of stimulated PBMCs did not result in substantial differences in cell type composition and activation marker fluorescence intensities between conditions^11,20,90^. The predefined panel, although chosen according to our best knowledge and analysed with advanced methods, might be too limited to pinpoint (subtle) stimulation specific changes. Therefore, we profiled the stimulated cells with a more unbiased single-cell transcriptomics approach as (1) no prior knowledge is required (whole transcriptome sequencing), (2) heterogeneous cell population (diverse cell states and lineages) are profiled in one sample, (3) cell lineages are described more accurately and (4) a broader overview of functional intra- and intercellular processes or interactions are obtained^13,91^. We optimized this approach to reduce technical bias, minimize processing time, and increase sample throughput by enriching for the B-cell compartment, and including nuclei hashing-based sample multiplexing. These experimental characteristics, next to the comparison of distinct stimulation strategies (conditions and incubation time points) makes our study unique in the field of *in vitro* B-cell stimulation.

Differential abundance testing (MiloR) and stimulation status annotation refinement of the cells was facilitated better capturing of subtle differences in transcriptional responses associated to a more continuous trajectory of (post-stimulation) B-cell development^48^. The differential abundant Nhoods indicated a remarkable overlap between IFN-γ/R-848 and ODN2006 (T-cell independent), and between MegaCD40L and anti-CD3/CD28 (T-cell dependent), showing the important role of stimulation mechanisms in the regulation towards the final cell fate. Annotation refinement highlighted significant differences between conditions in actual stimulation performance at specific incubation time points. IFN-γ/R-848 was observed to be the most effective stimulator due to showing the largest fraction of stimulated cells, even after a short incubation time, while anti-CD3/CD28 requires more time to (indirectly) stimulate higher fractions of cells, possibly even more than the longest included incubation time point (30h). This difference could be explained by the direct versus indirect mode of action of the respective stimuli. Cells annotates as ‘intermediate’ represented a heterogeneous population that dynamically evolved along the continuum between unstimulated and stimulated states. Although their enriched genes and pathways might provide insight into the transitional intracellular processes, this was beyond the scope of this study.

In a second phase, we defined the genes and pathways that are common and specific to each of the stimuli. This resulted in minor overlap when comparing the two time points within conditions, hinting towards a transition in transcriptional state depending on stimulation duration. The majority of DEGs was observed in IFN-γ/R-848 stimulated cells after 8h of stimulation but decreased after 30h. A similar trend was observed for ODN2006 and anti-CD3/CD28, although with less genes, while the number of detected DEGs in MegaCD40L stimulation was stable across both incubation time points. This might hint towards the end of stimulation potential of MegaCD40L around or after 30h, while for the other conditions shorter incubation durations suffice to reach full stimulation potential at transcriptomic level. We assessed specific genes that are extensively described in and related to (immune) diseases, such as DLBCL. Other genes might indirectly target specific pathways known to be involved in disease pathogenesis. *JUND*, for example, was uniquely downregulated in IFN-γ/R-848. It is strongly involved in the formation of the AP-1 complex, which is a crucial transcription factor influencing the Fas-mediated activation-induced cell death^68,69,92^. Alterations in this mechanism have been linked to various diseases such as inborn errors of immunity (e.g. autoimmune lymphoproliferative syndrome (ALPS), common variable immunodeficiency (CVID), systemic lupus erythematosus (SLE), autoimmune diseases, and cancer^93^. *In vitro* stimulation strategies for examination and determination of specific disease pathogenesis in which specific intracellular pathways are of interest, should therefore be selected with care. Next to the sets of genes uniquely linked to one of the stimuli, an overlap (including consistency in up- and downregulation) was observed between all four conditions showing genes known to be related to immune activation, cell proliferation and stress response. GO enrichment analysis finally revealed condition- and time-specific pathways, highlighting immune-linked processes and cell activation after 8h, and cell survival processes after 30h.

Finally, cell-cell communication analysis resulted in interesting interactions of which myeloid-involving interactions formed a major part^51^. The presence and importance of myeloid-B cell communication has been shown in previous studies, mainly linked to bone marrow tissue but also to circulating peripheral blood cells^91,94^. Bae *et al.* (2022)^94^ demonstrated the interaction between antigen-presenting and B cells based on extracted droplets displaying typical signatures for both cell types (doublets). In our analysis approach, multiplets were filtered out as distinguishing the (cell-)origin of the RNA fragments in these events remains challenging in a single-cell set-up, but it is not excluded that they may hold interesting information on closely interacting cells. IFN-γ/R-848 and anti-CD3/CD28 stimulated samples were associated to the majority of highly prioritized interactions. To our knowledge, no published studies are available describing the downstream and biological impact on cell-cell interactions of *in vitro* stimulation in detail, so we can only speculate on whether the interactions are directly induced by these conditions in the targeted cells, or are rather a cascade reaction of downstream triggered pathways in multiple activated cells. We also highlighted the differences in effect on downstream target gene expression regulation (up- or downregulation) of the interactions when comparing stimulations and the appearance of trans-interactions not yet described in literature.

Some caution should however be taken when interpreting cell-cell communication results. First, the quality scores, outputted during the preprocessing steps, were slightly higher for some of the samples than the predetermined thresholds. This was mainly observed in the B-cell subset and IFN-γ/R-848 stimulated samples and might be due to two main characteristics of the experimental set-up: (1) enrichment of the B-cell subset resulting in aberrant immune cell fractions compared to standard immune cell samples and (2) (B-cell) targeted stimulation of PBMCs resulting in upregulation of genes in specific cell subtypes only. Besides, we only focused on prioritized interactions in the top 50 ranking list per stimulation strategy, while a broader view might be interesting to reveal additional or extensive communication pathways. It should also be noted that this tool infers cell-cell communication based on transcriptomic profiles only, so experimental validation of the ligand-receptor interactions between cells, using for example proximity ligation or co-localization assays, will be needed to confirm these findings^95,96^.

Mainly cost and technical limitations inherent to single-cell workflows necessitated a balanced experimental approach regarding stimulation strategies, limiting the number of stimuli, concentrations and incubation time points that could be assessed. Of the four included stimuli, two conditions reflected the combined effect of multiple stimulation agents in one mixture, since MegaCD40L and IFN-γ/R-848 mixes included additional interleukins (IL-4 and IL-21) and anti-IgM antibodies (**Sup. Table S1**) following recommendations from previous studies to promote B-cell proliferation and cell stimulation^11,12^. However, in literature, it is also shown that these additional components are directly involved in B or broader immune cell stimulation, resulting in differences in gene expression, but these effects could not be uniquely filtered out in our data^26,59^. Our design only included 8h and 30h of stimulation, so very early and late induced genes and activated processes are most likely underrepresented in our atlas, underlining the need for additional time points to provide a broad longitudinal overview of the condition-specific intracellular effects. Another limitation of our study is that it is focused solely on transcriptomic profiling. In contrast to a scRNA-seq analysis, the scCITE-seq (single-cell Cellular Indexing of Transcriptomes and Epitopes by Sequencing) approach combines transcriptomic profiling with surface marker identification (cfr. FCM) to provide a multi-layered view at single-cell level^13,18,26,97^. This additional modality could facilitate more refined immune cell (sub)type annotation, especially of interest in the detailed annotation and characterization of B-cell subsets. Besides, it might provide insights in improving (novel) marker selection for the FCM panel that potentially pick up immune cell activation specific signals. Nonetheless, the transcriptome-focused approach still enabled an unbiased characterization of cell types and determination of their responses after stimulation with distinct conditions.

In conclusion, our data emphasizes that the choice of *in vitro* immune stimulation strategies, marked by the stimulatory condition and the incubation duration, is critical when targeted genes and cell-cell interactions, among others, are of interest. Rodríguez-Ubreva *et al.* (2022)^98^ emphasized the need for B-cell *in vitro* stimulation in the quest to understanding CVID pathogenesis in order to enable early diagnosis and improved patient treatment, while similar approaches were used in other research focusing on the (impaired) immune compartment^99–102^. Although, to our knowledge, no clear consensus on the use of specific stimulation strategies is available. Therefore, this study provides a valuable single-cell B-cell stimulation atlas in which we demonstrate fundamental differences on immune cell behaviour between distinct stimulation approaches, particularly with respect to T-cell (in)dependent stimulatory conditions and incubation time points. It offers a detailed documentation of targeted (intracellular) pathways and induced cell-cell interactions, functioning as a reference for selecting the most appropriate *in vitro* B-cell stimulation strategy and refinement of the experimental design when aiming to understand B-cell (dys)regulation and development in healthy individuals and disease conditions.

## MATERIALS AND METHODS

### Study participants details

Peripheral blood samples were collected from four healthy individuals (two males and two females, aged 27 to 49 years, Caucasian) at Ghent University Hospital, Ghent, Belgium. Blood samples were collected in accordance with the approved protocols of the Ethical Committee of Ghent University Hospital (EC number BC-10582). Written informed consent was obtained from each donor prior to sample collection.

### B-cell stimulation mixes

A panel of commonly used direct and indirect B-cell stimulators, representing both T-cell dependent and T-cell independent mechanisms (**Figure 1** and **Sup. Table S1**)^11,12,15,27^. T-cell independent stimulations included TLR9 agonist CpG oligodeoxynucleotide 2006 (ODN2006), and a combination of interferon-γ (IFN-γ) and the TLR7/8 agonist R-848 (Resiquimod). T-cell dependent stimulations included MegaCD40L (a CD40 ligand) anti-CD3/CD28 antibody-coated beads.

The stimulatory mixes included 20µL of anti-CD3/CD28 Dynabeads (Invitrogen, 11161D) per 1 million (M) Peripheral blood mononuclear cells (PBMCs), 0.1µg/mL CD40L (MEGACD40L® Protein, ENZO, ALX-522-110-C010), 50ng/mL IL-21 (Tebu-bio, 200-21-B), 50ng/mL IL-4 (Tebu bio, 200-04-A), 2.5µg/mL unmethylated CpG (ODN2006, ENZO, ALX-746-006-C100), 5µg/mL anti-IgM (Southern Biotech™, 202201), 50ng/mL IFN-γ (R&D Systems, 285-IF), and/or 5µg/mL R-848 (Resiquimod) (InvivoGen, tlrl-r848) (**Sup. Table S1**). The different combinations of B-cell stimulators are further referred to as “IFN-γ/R-848” (IFN-γ/R-848/anti-IgM/IL-4/IL-21), “ODN2006” (ODN2006), “MegaCD40L” (MegaCD40L/IL-4/IL-21), and “Anti-CD3/CD28” (anti-CD3/CD28 Dynabeads).

### PBMC isolation, thawing and *in vitro* stimulation

PBMCs were isolated within three hours of blood collections (K_2_EDTA anticoagulant) using a Ficoll-Paque density separation strategy and stored at -150°C (**Sup. Material D3**). Frozen PBMCs samples were thawed (**Sup. Material D3**) prior incubation at 37°C and 5% CO_2_ with different stimuli (**Sup. Table S1**) for 8h and 30h in 96-well plates. For flow cytometry, 1M PBMCs per healthy donor, condition and time point were incubated. For single-cell RNA sequencing (scRNA-seq) a total of 6M PBMCs per healthy donor, condition and time point, divided over 6 wells containing 1M PBMCs per well, were incubated. Cells incubated in complete Roswell Park Memorial Institute medium (RPMI) with B-cell stimulators were called ’stimulated’ samples, while those incubated in complete RPMI without stimuli were labeled as ’unstimulated’ samples. A total of 40 PBMC samples (ten stimulation conditions, four donors per condition) were analysed at 8h and 30h post-stimulation using flow cytometry (FCM; **Sup. Figure S1**) and scRNA-seq (**Sup. Table S3**).

### Flow cytometry workflow

#### FACS staining and acquisition

Following *in vitro* PBMC stimulation, cells were harvested, washed twice with phosphate-buffered saline (PBS) + 1% Bovine Serum Albumin (BSA), and centrifuged (5 minutes - 540g - 4°C). For staining purposes, eleven fluorochrome-conjugated monoclonal antibodies targeting surface maturation and activation markers were used to stain the (un)stimulated cells (**Sup. Table S2**). Unstained samples were diluted in 50µL of PBS-BSA. Cell suspensions were incubated in the dark for 20-30 minutes at room temperature before being washed and centrifuged (5 min – 400g – RT). After that, the stained cell suspensions were washed with PBS + 1% BSA and resuspended in 500µL PBS-BSA. Data acquisition was performed using a BD FACSLyric™ instrument (BD Biosciences, San José, CA), with up to 300,000 events recorded for each sample.

#### FACS data analysis

In R (v4.3.1), .fcs files were preprocessed to remove doublets (based on FCS-A and FCS-H), erroneous events, and events outside the cytometer’s detection range using the PeacoQC package^28^, with compensation and data transformation performed using flowCore^29^, and debris removal based on FCS-A and SSC-A characteristics using FlowSOM^30^. Preprocessed files were mapped onto a generated FlowSOM model (**Sup. Material D3**) finally resulting in cell phenotype annotation based on CD19, CD27, CD69, CD80, CD83, CD86 and IgD expression. B-cell maturation markers (CD27 and IgD) were used for B-cell annotation, while activation markers (e.g., CD69, CD83, CD86) distinguished activated from inactivated cells. Per activated (CD69^+^) and inactivated (CD69^-^) B cell subsets (CD27^-^IgD^+^, CD27^+^IgD^+^ and CD27^+^IgD^-^), cell fractions were calculated per sample (characterized by healthy donor, stimulatory condition and incubation time point), while mean fluorescence intensity (MFI) values of surface markers were normalized with respect to the unstimulated samples (**Sup. Material D3**).

### Single-cell RNA-seq experimental workflow

Stimulated PBMCs were tagged with cell hashing antibodies for sample multiplexing, enriched for B cells using Fluorescence-Activated Cell Sorting (FACS), and pooled equimolarly into six batches (three pools per incubation time point, **Sup. Table S3**) loading on a microfluidics device for scRNA-seq profiling.

#### Cell hashing, FACS sorting and sample pooling

After *in vitro* stimulation of approximately 6M PBMCs per condition, cells were collected, washed twice with an in-house buffer -i.e., phosphate-buffered saline (PBS) (Gibco, ThermoFisher Scientific,14190-169) + 0,2% bovine serum albumin (BSA) (Sigma-Aldrich, A7284-1L) + 0,09% sodiumazide (Sigma-Aldrich, 2002-500G)-, and centrifuged at (5 min – 540g – 4°C). Staining was performed on ice with 5µL CD19-PECy7 (J3-119) (Analis; IM3628) and 1µL Fixable Viability dye eFluor506 (Fisher Scientific; 65-0866-14). TotalSeq™-C anti-human hashtags (2µL, n=7), conjugated to hashtag oligonucleotide (HTO) uniquely associated to a specific sample (**Sup. Table S3**), were added for sample multiplexing and incubated for 30 minutes at 4°C in the dark. Cells were then washed and resuspended in 200µL PBS+BSA. CD19^+^ and CD19^-^ living cells were sorted using BD FACSymphony™ S6, BD FACSMelody™ Cell Sorter, or BD FACSVULCAN™ (**Sup. Table S3**).

B-cell enrichment (expected fraction of 5-17% of lymphocytes in healthy adults) using FACS sorting was added to the scRNA-seq workflow due to the B-cell focus of this study^31^. HTO-tagged FACS-sorted cell fractions (CD19^+^ B cells and CD19^-^ non-B cells) were pooled at a 60/40 ratio. PBMCs from four healthy controls and five ((un)stimulated) conditions were pooled into three groups, with up to seven samples per incubation time point (CEV017-CEV022; **Sup. Table S3**). **Sup. Table S3**includes the number of sorted CD19^+^ cells per sample and pool (349,390 to 733,143) and additional pooling and sorting. This strategy ensured each pool uniquely represented a combination of control samples and stimulation conditions, facilitating data demultiplexing based on single nucleotide polymorphism (SNP) genotyping and HTO cell hashing, and removal of technical bias.

#### Single-cell library preparation and pooling

Sample pools (n=6, CEV017-CEV022) were counted with a Luna-FL Cell Counter, and all showed >95% viability. Cells were centrifuged (6 min – 400g – 4°C) and resuspended to enable loading of 50,000 cells onto the Chromium Controller (10X Genomics). Single-cell library preparation was performed using Chromium Next GEM Single Cell 5’ Reagent Kits v2 (10X Genomics), aiming for the generation of nanoliter-scale gel beads-in-emulsion (GEMs) with one cell and one bead per droplet. RNA from the cells was captured by the oligonucleotide-coated beads, and dual indexing with i5 and i7 indices was used to differentiate sample pools (CEV017-CEV022), finally resulting in a 10X Genomics 5’ cDNA library (Dual Index Kit TT). The final library (average fragment length of 422bp, diluted to 4nM) was sequenced in a single run on an Illumina Novaseq 6000 using a Novaseq 6000 S2 Reagent Kit v1.5 (100 cycles; 20028316) for paired-end, dual-indexed sequencing (i5 and i7 indices), involving 26-10-10-90 cycles (R1-I7-I5-R2) with a final loading concentration of 0.35nM (1% PhiX). A total of 2.34 billion reads (passing filter) were obtained with a high base calling accuracy (95.24% >Q30) and an overall alignment of 9.63% of PhiX fragments.

### Single-cell RNA-seq data-analysis workflow

#### Data preprocessing

All following data and statistical analysis steps were performed in R (v4.4.1) using the High Performance Computing infrastructure of Ghent University^32^. After sequencing, the 10X Genomics CellRanger v7.0.0^33^ pipeline was used to demultiplex the samples (CEV017-CEV022; cellranger mkfastq –barcode-mismatches=0), align cDNA reads to the GRCh38 genome (v100)^34^, and link the reads to the associated cell barcode (cellranger multi). Count matrices for cDNA were corrected for amplification bias using unique molecular identifiers (UMIs). A total of 19,892 to 36,563 droplets were captured per sample pool with an average of 253,814,172 reads (range: 153,957,721 – 304,378,211) per pool. Metrics per cell across pools were similar: median reads (mean = 5,547, range: 4,618 – 6,335), unique molecular identifiers (UMI; mean = 2,074, range: 1,682 – 2,668), and gene counts (mean = 995 range: 831 – 1,227). The results were integrated into a Seurat object using Seurat V5 (version 5.1.0), using Seurat::Read10X() and Seurat::CreateSeuratObject(), for feature extraction, cell labeling, and visualization^35^. Cells were filtered based on unique RNA molecules (<200) and mitochondrial gene expression (<25%).

#### Sample demultiplexing

For sample demultiplexing, two methods were combined to achieve consensus on cell annotation. First, the reference-free freemuxlet() function from the Popscle package ^36^ was used to demultiplex based on SNP profiles, with an adapted and filtered input reference VCF file of the 1000 Genomes project (v100)^37^. Unambiguous cells were annotated as “singlets”, linked to a specific donor, or “doublet”. The remaining cells stayed “non-defined”. Second, sample demultiplexing and multiplet detection based on HTO counts were performed with the Seurat::HTODemux() function (**Sup. Figure S2**). A consensus on final cell annotation was achieved by combining the results from both methods. On average, 15,493 events per pool were recovered (range: 10,019 – 19,267). Details on the composition of each pool in fractions of singlets linked to healthy control, stimulatory condition, or HTO are shown in **Sup. Figure S3**.

#### Data integration

The preprocessed Seurat objects from the six sample pools (CEV017-022) were integrated using appropriate Seurat functions (**Sup. Material D3**)^35^. A Canonical Correlation Analysis (CCA) integration approach was used taking into account the biological variance explained by the combination of healthy individual and stimulatory condition (e.g. *HD1_MegaCD40L*), prior dimensionality reduction and conventional Louvain clustering of the cells (**Sup. Material D3**).

#### Doublet detection and cell annotation based on the Azimuth PBMC reference

Doublet detection in the integrated Seurat object was performed using the scDblFinder package^38^ (**Sup. Material D3**). After removing 12.7% of the 92,958 total events annotated as multiplets, a final dataset of 81,160 singlets was obtained (**Sup. Figure S4A**). These singlets were then annotated (1) by mapping the transcriptome profiles on a multi-modal human PBMC reference atlas (Azimuth^39^) and (2) based on expression levels of immune cell-type specific genes as reported elsewhere (**Sup. Figure S4A**)^40–47^. The former approach consists of a transformation step (Seurat::SCTransform()) and anchor extraction between the reference and query data (Seurat::FindTransferAnchors()) prior to reference label mapping (Seurat::MapQuery()). Finally, a consensus label (B, CD4^+^ T, CD8^+^ T, other T, NK, and myeloid cells) between both approaches was assigned to each of the 28 conventional clusters, obtained after full data set integration, and visualized using a UMAP approach (**Sup. Figure S4B-C**).

B cells made up 56.3% of the total cells (n=45,676), meeting the enrichment goal of 60%. Other cell types included CD4+ T cells (18.5%, n=15,009), CD8+ T cells (11.6%, n=9,382), other T cells (2.2%, n=1,803), NK cells (4.8%, n=3,901), and myeloid cells (6.6%, n=5,389, consisting of dendritic cells and monocytes) (**Sup. Figure S4B**). The fractions of B cells by stimulation condition and incubation time were consistent with expectations (**Sup. Figure S4D**). B cells (n=45,676) were further divided into those incubated for 8h (45.8%, n=20,937) or 30h (54.2%, n=24,739). After B-cell extraction, CCA integration, dimensionality reduction and conventional Louvain clustering was applied on both 8h and 30h stimulated cell subsets separately to remove technical variability in a similar approach as described for the total immune cell data set (**Figure 3A-B, Sup. Material D3**).

### In-depth analysis of B-cell subsets

#### B-cell subset selection, filtering and integration

For 8h and 30h of stimulation, a total of 24 and 26 initial clusters, respectively, were observed. An additional filtering step of these clusters based on cell quality and cell type purity (**Sup. Material D3**) resulted in final filtered B-cell subsets of 20,185 and 22,915 for 8h and 30h of stimulation, respectively, that underwent a final round of CCA integration, dimensionality reduction and conventional Louvain clustering prior to downstream analysis (**Sup. Material D3**).

#### Generation and annotation of neighbourhoods

The filtered B-cell subsets (8h and 30h of stimulation separately) was used as input for the functions of the miloR^48,49^ package to finally obtain differential abundance (DA) regions for the distinct stimulatory conditions (**Sup. Material D3**). This analysis included generation of a k-nearest nearest neighbour graph enabling identification of neighbourhoods (Nhoods), which is a unique set of highly similar cells. Since one cell can be linked to multiple Nhoods, the continuum of cell lineages is conserved. Differential abundance testing was performed by defining specific contrasts for every stimulation separately, in which the unstimulated cells were used as reference (e.g. ‘StimulationMegaCD40L – StimulationUnstimulated’), and inputted in the miloR::testNhoods(…, fdr.weighting=‘graph-overlap’) function. Obtained Nhoods were then visualized on the CCA integrated UMAP embeddings and annotated with miloR::annotateNhoods() using the original B-cell clusters as reference labels. Since most of the Nhoods contain a mixture of cells linked to distinct clusters, the final label is the majority cluster defined as the cluster with the highest fraction in that specific Nhood. If the highest fraction is smaller than 70%, the Nhood is annotated as “Mixed” and is not considered for further analysis. Finally, beeswarm plots per contrast were obtained with the miloR::plotDAbeeswarm(…, alpha=0.1).

This analysis was performed for the filtered B-cell subset for 8h and 30h of stimulation separately (**Sup. Material D3**).

#### Selection of differentially abundant (un)stimulated neighbourhoods

(Un)stimulated Nhoods were selected using two criteria. First, for each of the time points and conditions separately, Nhoods linked to specific Louvain algorithm-based clusters were selected. The Louvain clusters were selected if they showed enrichment for highly differentially abundant Nhoods based on the generated beeswarm plots (**Sup. Material D3**, **Figure 3G-H** and **Sup. Figure S5**). Secondly, the selected Nhoods were only retained for downstream analysis if the differential abundance test resulted in a |logFC|>3. This can be interpreted as a twenty-fold change (e^3^) between stimulated and unstimulated cells. To ensure an informative subset of unstimulated Nhoods for downstream analysis steps, overlapping Nhoods selected in at least three conditions were finally retained. If a cell was linked to a stimulated and an unstimulated Nhoods, it was finally annotated as ‘Unstimulated’ (**Sup. Material D3**). This selection was visualized using an in-house adapted version of the miloR::plotDAbeeswarm(…, alpha=0.1) by representing the retained and ignored Nhoods as solid dots and empty circles, respectively (**Figure 3H** and **Sup. Figure S5B**).

#### Refinement of cell annotation related to stimulation status

A refinement of the annotation related to cell stimulation status was performed based on the selected Nhoods per condition and the overlap with the metadata annotation of the sample that the cells originate from (**Sup. Material D3**). This resulted in ‘truly stimulated’ and ‘truly unstimulated’ cells identified per stimulatory condition and incubation time point. Cells belonging to Nhoods that were not retained at the end of the selection process were annotated after refinement as ‘intermediate’. This refined stimulation status annotation of the cells is used in the downstream analysis steps. The unstimulated cells, a unique subset per incubation time point, were used as reference for differential gene analysis in each of the conditions from (**Sup. Material D3**).

#### Fraction of truly stimulated cells per condition

The fraction of truly stimulated cells per condition was obtained by dividing the number of stimulated cells based on the selected Nhoods and cell metadata (as described above) by the total number of cells in the data set (solely based on cell metadata and independent of the selected Nhoods) with respect to the condition, incubation time points and healthy donors. As a reference for natural stimulation of cells incubated in wells without an *in vitro* stimulus, the fraction of unstimulated cells present in the selected stimulated Nhoods was calculated in a similar approach. For example, after 8h of stimulation, 2,428 cells were annotated as truly stimulated with IFN-γ/R-848 for the first healthy donor. The total number of cells recovered for this donor and associated to 8h of incubation with IFN-γ/R-848 was 2,799 (solely based on metadata information), resulting in a fraction of 86.7% of the starting cells that was truly seen as stimulated (**Figure 2I, Sup. Figure S6**). Within the selected Nhoods finally annotated as truly stimulated with IFN-γ/R-848, 21 cells originated from an unstimulated sample. The total number of cells recovered for the first healthy donor and associated to 8h of incubation in the unstimulated condition was 855, resulting in a fraction of 2.5% of the starting cells that have been ‘naturally stimulated’ in a similar way as IFN-γ/R-848, despite the absence of this stimulator in the incubation well (**Sup. Figure S6**). Finally, an average fraction over healthy donors was obtained, indicating stimulation potential of the stimulatory conditions per incubation time point (**Figure 2I**).

#### Differential gene expression analysis

The subsets of (un)stimulated cells per incubation time point were used as input for the differential expression analysis using the Seurat::FindMarkers(…, min.pct = 0, logfc.threshold = 0, ident.1 = <STIM>, ident.2 = “Unstimulated”) where <STIM> is adapted for each of the four stimulatory conditions so that these conditions are analysed separately, and the unstimulated cells are used as a reference. The arguments min.pct and logfc.threshold are set to zero to ensure the entire set of genes is tested and outputted, necessary for downstream gene set enrichment testing (hypergeometric testing, HGT). This results in output values including avg_logFC, p_val_adj (based on Bonferroni correction), pct.1 and pct.2 (percentage of cells where the gene is detected in the test and reference group, respectively). An additional normalized expression metric was calculated and annotated as “gene score”, of importance in gene prioritization, by implementing the following formula: Score = (pct.1 / (pct.2 + 0.01)) * avg_log2FC). Genes were annotated as “upregulated” and “downregulated” if the adjusted p-value was smaller than 0.05 and the log2FC was larger than 2 or smaller than -2, respectively. An overview of all genes is available in **Sup. Material D1** and **D2**, for both 8h and 30h of incubation respectively. Remaining genes were annotated as “NS” (non-significant). For visualization, only the top five genes after arranging based on the gene score were shown (**Figure 4D** and **Sup. Figure S9**).

#### Gene set enrichment analysis and biological interpretation

Gene set enrichment analysis was performed using the Gene Ontology (GO) gene set for biological processes, specific for Homo sapiens, available in the fifth category of the Molecular Signature Database (MSigDB) collections^50^. These gene sets were loaded using msigdbr::msigdbr(species = ‘Homo sapiens’, category = ‘C5’, subcategory = ‘BP’). Hypergeometric testing was performed on subsets of differentially expressed genes by splitting based on stimulatory condition, time point of incubation and expression of direction (up- or downregulated). Statistical testing was performed using the stats::phyper (…, lower.tail=FALSE) expression and multiple testing was accounted for using the stats::p.adjust(…, method = ‘BH’).

### Cell-cell communication between large immune cell subsets

Differential cell-cell communication analysis was performed using the MultiNicheNetR^51,52^ package conform the described procedure by Browaeys *et al.*^51^ for the 8h and 30h stimulated cells separately (**Sup. Material D3**). The contrasts were defined by comparing the stimulatory condition to the unstimulated samples (e.g. ‘IFN-γ/R-848’-‘Unstimulated’), and all cell types were considered as possible senders and receivers (B, CD8+ T, CD4+ T, NK and Myeloid cells). MultiNicheNetR was then run including selection of sufficiently abundant cell types (minimal ten cells) and genes (fraction_cutoff=0.05 and min_sample_prop=0.5), calculation of pseudobulk expression, differential expression analysis, gene set assessment (logfc_threshold=0.5, p_val_adj=FALSE, p_val_threshold=0.05), ligand activity prediction and cell-cell communication pattern determination (top_n_target=250), and interaction scoring (ligand_activity_down=FALSE, scenario=’lower_DE’). The Other T cell subset did not meet the filtering requirements and was therefore left out for downstream analysis. This finally resulted in 5,398-8,135 retained genes per incubation time point and cell subtype used for cell-cell communication analysis To get a general overview, the inferred interactions were ranked based on the obtained score, enabling the extraction of the top 50 overall interactions (**Figure 5A and Sup. Figure S12**) and the top 50 interactions per stimulatory condition (**Sup. Figure S13-S20**). More in-depth prioritization of the top 50 interactions per stimulatory conditions was performed based on three criteria: (1) ligand activity, (2) prior knowledge linking ligand-receptor pairs to downstream target genes, and (3) concordance between upstream ligand-receptor presence and downstream target genes expression (absolute value of Pearson correlation > 0.8). This resulted in seven to twenty highly trustworthy, prioritized interactions for each of the stimulatory condition (**Figure 5B-C**). This analysis is performed for subsets obtained after 8h and 30h of stimulation separately.

### Visualizations

UMAPs were plotted using the Seurat::DimPlot() function. Upset plots were generated using the ComplexHeatmap^53^ R package. Other figures were created with the ggplot2 package^54^, or related packages such as GGally^55^. Graphical summaries were generated with BioRender.. Circos diagrams and bubble plots representing the cell-cell interactions were generated using the MultiNicheNetR package^51,52^.

### Quantification and statiscal analysis

For statistical testing of obtained fractions in the flow cytometry data (**Figure 2** and **Sup. Figure S1B**) and single-cell data after refinement of the stimulation status (**Figure 3I**), a generalized linear mixed model (GLMM) based on beta-family regression with the logit-link function was used: glmmTMB(…, family=glmmTMB::beta_family(link=“logit”)^56^. It should be noted that a donor effect was observed when comparing cell fractions between samples in both datasets (**Sup. Figure S6**), which is accounted for in this statistical model approach. For the flow cytometry data, the formula ‘Fraction ∼ Condition × Phenotype + HealthyControl × Phenotype + TimePoint × Condition was used. For the fractions obtained after refinement of the stimulation status in the single-cell dataset, a division based on final annotation (Stimulated or Unstimulated) of the data was performed prior to GLMM training. The model, that accounts for multiple testing correction, was finally built based on the formula ‘Fraction ∼ TimePoint × Condition + HealthyControl’. Contrasts were finally defined for each of the datasets separately and tested using the multcomp::glht() function with the corresponding trained model. Significance of the differences was tested using a significance level α=0.05

## Supporting information

This document contains supplementary tables and figures referred to in the manuscript.

List of significantly differentially expressed genes obtained after 8 hours of stimulation (referred to as Sup. Material D1 in the manuscript).

List of significantly differentially expressed genes obtained after 30 hours of stimulation (referred to as Sup. Material D2 in the manuscript).

Supplementary information on the Materials and Method section (referred to as Sup. Material D3 in the manuscript).

## DATA AVAILABILITY

Raw sequencing FastQ files and processed data (including CellRanger output files) of all samples are published in the European Genome-Phenome Archive (EGA) with accession ID EGAD50000002998.

## CODE AVAILABILITY

The main code for data analysis and visualization, using existing packages and software tools, are available at GitHub (https://github.com/VIBTOBIlab/scRNAseq_InVitro_ImmuneStimulation).

## ACKNOWLEDGEMENTS

The authors would like to thank the healthy blood donors for participating in the study. We acknowledge the VIB Single Cell Core and VIB Single Cell Flow Core facilities (Zwijnaarde, Belgium) for their support and assistance during the experimental work. We also acknowledge Malicorne Buysse, Karlien Claes, Sofie Roelandt, and Alvaro Marcos Rubio for their technical support and assistance in blood sampling and sample processing. Furthermore, we are grateful for the support of Ruth Seurinck in the cell-cell communication data analysis and Leander Meuris in the statistical analysis. The computational resources (Stevin Supercomputer Infrastructure) and services used in this work were provided by the VSC (Flemish Supercomputer Center), funded by Ghent University, FWO and the Flemish Government – department EWI.

## AUTHOR CONTRIBUTIONS

Tine D’hamers: data curation, formal analysis, methodology, visualization, writing – original draft. Jana Neirinck: conceptualization, data curation, funding acquisition, methodology, project administration, writing – original draft. Malicorne Buysse: data curation, methodology. Ciel De Vriendt, Tessa Kerre and Filomeen Haerynck: resources, writing – review and editing. Mattias Hofmans, Carolien Bonroy and Celine Everaert: conceptualization, funding acquisition, methodology, supervision, writing – review and editing.

## ETHICAL STATEMENT

The study is approved by the Ethical Committee of Ghent University Hospital (EC number BC-10582). Informed consent was obtained from all subjects involved in the study.

## REFERENCES

1. Charles A Janeway, J., Travers, P., Walport, M. & Shlomchik, M. J. B-Cell Activation by Armed Helper T Cells. (Garland Science, New york, 2001).

2. Recaldin, T. & Fear, D. J. Transcription factors regulating B cell fate in the germinal centre. Clin. Exp. Immunol. 183, 65–75 (2016).

3. Akkaya, M. et al. Second signals rescue B cells from activation-induced mitochondrial dysfunction and death. Nat. Immunol. 19, 871–884 (2018).

4. Tsai, D.-Y., Hung, K.-H., Chang, C.-W. & Lin, K.-I. Regulatory mechanisms of B cell responses and the implication in B cell-related diseases. J. Biomed. Sci. 26, 64 (2019).

5. Doherty, D. G., Melo, A. M., Moreno-Olivera, A. & Solomos, A. C. Activation and Regulation of B Cell Responses by Invariant Natural Killer T Cells. Front. Immunol. 9, (2018).

6. Bogaert, D. J. et al. The immunophenotypic fingerprint of patients with primary antibody deficiencies is partially present in their asymptomatic first-degree relatives. Haematologica 102, 192–202 (2017).

7. Le Coz, C. et al. Common variable immunodeficiency-associated endotoxemia promotes early commitment to the T follicular lineage. J. Allergy Clin. Immunol. 10.1016/j.jaci.2019.08.007 (2019) doi:10.1016/j.jaci.2019.08.007.

8. Driessen, G. J. et al. Common variable immunodeficiency and idiopathic primary hypogammaglobulinemia: two different conditions within the same disease spectrum. Haematologica 98, 1617–23 (2013).

9. Song, S. & Matthias, P. D. The Transcriptional Regulation of Germinal Center Formation. Front. Immunol. 9, (2018).

10. Laidlaw, B. J. & Cyster, J. G. Transcriptional regulation of memory B cell differentiation. Nat. Rev. Immunol. 21, 209–220 (2021).

11. Marsman, C. et al. Optimized Protocols for In-Vitro T-Cell-Dependent and T-Cell-Independent Activation for B-Cell Differentiation Studies Using Limited Cells. Front. Immunol. 13, 815449 (2022).

12. Van Belle, K. et al. Comparative In Vitro Immune Stimulation Analysis of Primary Human B Cells and B Cell Lines. J. Immunol. Res. 2016, 5281823 (2016).

13. Lawlor, N. et al. Single Cell Analysis of Blood Mononuclear Cells Stimulated Through Either LPS or Anti-CD3 and Anti-CD28. Front. Immunol. 12, 636720 (2021).

14. Bishop, G. A. & Hostager, B. S. Signaling by CD40 and its mimics in B cell activation. Immunol. Res. 24, 97–109 (2001).

15. Keller, B., et al. The expansion of human T-bethighCD21low B cells is T cell dependent. Sci. Immunol. 6, eabh0891 (2021).

16. Zumaquero, E. et al. IFNγ induces epigenetic programming of human T-bethi B cells and promotes TLR7/8 and IL-21 induced differentiation. eLife 8, e41641 (2019).

17. Kochayoo, P. et al. Interferon-γ signal drives differentiation of T-bethi atypical memory B cells into plasma cells following Plasmodium vivax infection. Sci. Rep. 12, 4842 (2022).

18. Aguiar, V. R. C. et al. A multi-omics resource of B cell activation reveals genetic mechanisms for immune-mediated diseases. medRxiv 2025.05.22.25328104 (2025) doi:10.1101/2025.05.22.25328104.

19. Calderon, D. et al. Landscape of stimulation-responsive chromatin across diverse human immune cells. Nat. Genet. 51, 1494–1505 (2019).

20. Pierzchalski, A., Zenclussen, A. C. & Herberth, G. A comprehensive battery of flow cytometric immunoassays for the in vitro testing of chemical effects in human blood cells. Front. Immunol. 14, (2024).

21. Khoenkhoen, S., Ádori, M., Pedersen, G. K. & Karlsson Hedestam, G. B. Flow Cytometry-Based Protocols for the Analysis of Human Plasma Cell Differentiation. Front. Immunol. 11, (2020).

22. Mylka, V. et al. Comparative analysis of antibody- and lipid-based multiplexing methods for single-cell RNA-seq. Genome Biol. 23, 55 (2022).

23. Stoeckius, M. et al. Cell Hashing with barcoded antibodies enables multiplexing and doublet detection for single cell genomics. Genome Biol. 19, 224 (2018).

24. Kang, H. M. et al. Multiplexed droplet single-cell RNA-sequencing using natural genetic variation. Nat. Biotechnol. 36, 89–94 (2018).

25. Jovic, D. et al. Single-cell RNA sequencing technologies and applications: A brief overview. Clin. Transl. Med. 12, e694 (2022).

26. Cui, A. et al. Dictionary of immune responses to cytokines at single-cell resolution. Nature 625, 377–384 (2024).

27. Rodríguez-Ubreva, J. et al. Single-cell Atlas of common variable immunodeficiency shows germinal center-associated epigenetic dysregulation in B-cell responses. Nat. Commun. 13, 1779 (2022).

28. Emmaneel, A. et al. PeacoQC: Peak-based selection of high quality cytometry data. Cytom. Part J. Int. Soc. Anal. Cytol. 101, 325–338 (2022).

29. Ellis, B., et al. flowCore: flowCore: Basic Structures for Flow Cytometry Data. (2022).

30. Van Gassen, S. et al. FlowSOM: Using self-organizing maps for visualization and interpretation of cytometry data. Cytom. Part J. Int. Soc. Anal. Cytol. 87, 636–45 (2015).

31. van der Burg, M. et al. The EuroFlow PID Orientation Tube for Flow Cytometric Diagnostic Screening of Primary Immunodeficiencies of the Lymphoid System. Front. Immunol. 10, 246 (2019).

32. Team, R. C. R: A Language and Environment for Statistical Computing. https://www.R-project.org/ (2020).

33. Zheng, G. X. Y. et al. Massively parallel digital transcriptional profiling of single cells. Nat. Commun. 8, 14049 (2017).

34. Dyer, S. C. et al. Ensembl 2025. Nucleic Acids Res. 53, D948–D957 (2025).

35. Hao, Y. et al. Dictionary learning for integrative, multimodal and scalable single-cell analysis. Nat. Biotechnol. 42, 293–304 (2024).

36. statgen/popscle. Center for Statistical Genetics (2024).

37. Auton, A. et al. A global reference for human genetic variation. Nature 526, 68–74 (2015).

38. Germain, P.-L., Lun, A., Garcia Meixide, C., Macnair, W. & Robinson, M. D. Doublet identification in single-cell sequencing data using scDblFinder. F1000Research 10, 979 (2022).

39. Hao, Y. et al. Integrated analysis of multimodal single-cell data. Cell 184, 3573–3587.e29 (2021).

40. Espinoza, D. A. et al. Distinct stage-specific transcriptional states of B cells derived from human tonsillar tissue. https://insight.jci.org/articles/view/155199/figure/1 (2023) doi:10.1172/jci.insight.155199.

41. Holla, P. et al. Shared transcriptional profiles of atypical B cells suggest common drivers of expansion and function in malaria, HIV, and autoimmunity. Sci. Adv. 7, eabg8384 (2021).

42. King, H. W., et al. Single-cell analysis of human B cell maturation predicts how antibody class switching shapes selection dynamics. Sci. Immunol. 6, eabe6291 (2021).

43. Stewart, A. et al. Single-Cell Transcriptomic Analyses Define Distinct Peripheral B Cell Subsets and Discrete Development Pathways. Front. Immunol. 12, (2021).

44. Rose, J. R. et al. Distinct transcriptomic and epigenomic modalities underpin human memory T cell subsets and their activation potential. *Commun*. Biol. 6, 1–19 (2023).

45. Szabo, P. A. et al. Single-cell transcriptomics of human T cells reveals tissue and activation signatures in health and disease. Nat. Commun. 10, 4706 (2019).

46. Wang, X. et al. Reinvestigation of Classic T Cell Subsets and Identification of Novel Cell Subpopulations by Single-Cell RNA Sequencing. J. Immunol. 208, 396–406 (2022).

47. Stuart, T. et al. Comprehensive Integration of Single-Cell Data. Cell 177, 1888–1902.e21 (2019).

48. Dann, E., Henderson, N. C., Teichmann, S. A., Morgan, M. D. & Marioni, J. C. Differential abundance testing on single-cell data using k-nearest neighbor graphs. Nat. Biotechnol. 40, 245–253 (2022).

49. Morgan, D. M. & Dann, E. miloR: Differential neighbourhood abundance testing on a graph. (2025).

50. Liberzon, A. et al. The Molecular Signatures Database (MSigDB) hallmark gene set collection. Cell Syst. 1, 417–425 (2015).

51. Browaeys, R. et al. MultiNicheNet: a flexible framework for differential cell-cell communication analysis from multi-sample multi-condition single-cell transcriptomics data. 2023.06.13.544751 Preprint at 10.1101/2023.06.13.544751 (2023).

52. Browaeys, R., Saelens, W. & Saeys, Y. NicheNet: modeling intercellular communication by linking ligands to target genes. Nat. Methods 17, 159–162 (2020).

53. Gu, Z., Eils, R. & Schlesner, M. Complex heatmaps reveal patterns and correlations in multidimensional genomic data. Bioinformatics 32, 2847–2849 (2016).

54. Wickham, H. *Ggplot2*. (Springer International Publishing, Cham, 2016). doi:10.1007/978-3-319-24277-4.

55. Schloerke, B., et al. GGally: Extension to ‘ggplot2’. (2025).

56. Brooks, M. E. et al. glmmTMB Balances Speed and Flexibility Among Packages for Zero-inflated Generalized Linear Mixed Modeling. R J. 9, 378–400 (2017).

57. van Dongen, J. J. M. et al. EuroFlow-Based Flowcytometric Diagnostic Screening and Classification of Primary Immunodeficiencies of the Lymphoid System. Front. Immunol. 10, (2019).

58. Neirinck, J., et al. The Euroflow PID Orientation Tube in the diagnostic workup of primary immunodeficiency: Daily practice performance in a tertiary university hospital. Front. Immunol. 13, (2022).

59. Miller, R. A. et al. Anti-CD73 antibody activates human B cells, enhances humoral responses and induces redistribution of B cells in patients with cancer. J. Immunother. Cancer 10, e005802 (2022).

60. Becker-Herman, S., Lantner, F. & Shachar, I. Id2 Negatively Regulates B Cell Differentiation in the Spleen1. J. Immunol. 168, 5507–5513 (2002).

61. Ji, M. et al. Id2 intrinsically regulates lymphoid and erythroid development via interaction with different target proteins. Blood 112, 1068–1077 (2008).

62. Zaccolo, M., Zerio, A. & Lobo, M. J. Subcellular Organization of the cAMP Signaling Pathway. Pharmacol. Rev. 73, 278–309 (2021).

63. Bielenberg, M., Kurelic, R., Frantz, S. & Nikolaev, V. O. A mini-review: phosphodiesterases in charge to balance intracellular cAMP during T-cell activation. Front. Immunol. 15, 1365484 (2024).

64. Cooney, J. D. & Aguiar, R. C. T. Phosphodiesterase 4 inhibitors have wide-ranging activity in B-cell malignancies. Blood 128, 2886–2890 (2016).

65. Bertoni, F. & Stathis, A. Staining the target: CD37 expression in lymphomas. Blood 128, 3022–3023 (2016).

66. Xu-Monette, Z. Y. et al. Assessment of CD37 B-cell antigen and cell of origin significantly improves risk prediction in diffuse large B-cell lymphoma. Blood 128, 3083–3100 (2016).

67. Winde, C. M. de et al. Tetraspanin CD37 protects against the development of B cell lymphoma. J. Clin. Invest. 126, 653–666 (2016).

68. Weitzman, J. B., Fiette, L., Matsuo, K. & Yaniv, M. JunD Protects Cells from p53-Dependent Senescence and Apoptosis. Mol. Cell 6, 1109–1119 (2000).

69. Wu, Y. et al. B cell function in patients with systemic lupus erythematosus is regulated by the upregulation of JunD. Heliyon 10, e35949 (2024).

70. Li, C., Ibrahim, M. M. & Fang, C. MFSD2A: a molecular nexus linking blood–brain barrier, lipid metabolism, and ischemia–reperfusion injury. Cell. Mol. Biol. Lett. 30, 148 (2025).

71. Piccirillo, A. R. et al. The Lysophosphatidylcholine Transporter MFSD2A Is Essential for CD8+ Memory T Cell Maintenance and Secondary Response to Infection. J. Immunol. 203, 117–126 (2019).

72. Takahashi, K. et al. CCL3 and CCL4 are biomarkers for BCR pathway activation and prognostic serum markers in diffuse large B cell lymphoma (DLBCL). Br. J. Haematol. 171, 726–735 (2015).

73. Kato, A. et al. Interferon-α/β receptor-mediated selective induction of a gene cluster by CpG oligodeoxynucleotide 2006. BMC Immunol. 4, 8 (2003).

74. Wurtz, O., Bajénoff, M. & Guerder, S. IL□4□mediated inhibition of IFN□γ production by CD4+ T cells proceeds by several developmentally regulated mechanisms. Int. Immunol. 16, 501–508 (2004).

75. Zhao, Y. et al. PD-L1:CD80 Cis-Heterodimer Triggers the Co-stimulatory Receptor CD28 While Repressing the Inhibitory PD-1 and CTLA-4 Pathways. Immunity 51, 1059–1073.e9 (2019).

76. Sugiura, D. et al. Restriction of PD-1 function by cis-PD-L1/CD80 interactions is required for optimal T cell responses. Science 364, 558–566 (2019).

77. Bailey, C. C., Zhong, G., Huang, I.-C. & Farzan, M. IFITM-Family Proteins: The Cell’s First Line of Antiviral Defense. Annu. Rev. Virol. 1, 261–283 (2014).

78. Carty, M. & Bowie, A. G. Recent insights into the role of Toll-like receptors in viral infection. Clin. Exp. Immunol. 161, 397–406 (2010).

79. She, S. et al. Functional Roles of Chemokine Receptor CCR2 and Its Ligands in Liver Disease. Front. Immunol. 13, (2022).

80. Gschwandtner, M., Derler, R. & Midwood, K. S. More Than Just Attractive: How CCL2 Influences Myeloid Cell Behavior Beyond Chemotaxis. Front. Immunol. 10, 2759 (2019).

81. Fleige, L. & Capellino, S. Simultaneous Stimulation of Peripheral Blood Mononuclear Cells with CpG ODN2006 and α-IgM Antibodies Leads to Strong Immune Responses in Monocytes Independent of B Cell Activation. Cells 13, 1822 (2024).

82. Alspach, E., Lussier, D. M. & Schreiber, R. D. Interferon γ and Its Important Roles in Promoting and Inhibiting Spontaneous and Therapeutic Cancer Immunity. Cold Spring Harb. Perspect. Biol. 11, a028480 (2019).

83. Bui, T. M., Wiesolek, H. L. & Sumagin, R. ICAM-1: A master regulator of cellular responses in inflammation, injury resolution, and tumorigenesis. J. Leukoc. Biol. 108, 787–799 (2020).

84. Guerra-Espinosa, C., Jiménez-Fernández, M., Sánchez-Madrid, F. & Serrador, J. M. ICAMs in Immunity, Intercellular Adhesion and Communication. Cells 13, 339 (2024).

85. Burton, J., Goldman, C. K., Rao, P., Moos, M. & Waldmann, T. A. Association of intercellular adhesion molecule 1 with the multichain high-affinity interleukin 2 receptor. Proc. Natl. Acad. Sci. 87, 7329–7333 (1990).

86. Li, Y., Li, X., Geng, X. & Zhao, H. The IL-2A receptor pathway and its role in lymphocyte differentiation and function. Cytokine Growth Factor Rev. 67, 66–79 (2022).

87. Malek, T. R. & Castro, I. Interleukin-2 Receptor Signaling: At the Interface between Tolerance and Immunity. Immunity 33, 153–165 (2010).

88. Zhou, Y., Richmond, A. & Yan, C. Harnessing the Potential of CD40 Agonism in Cancer Therapy. Cytokine Growth Factor Rev. 75, 40–56 (2024).

89. Deng, Z. et al. TGF-β signaling in health, disease and therapeutics. Signal Transduct. Target. Ther. 9, 61 (2024).

90. Rincon-Arevalo, H. et al. Differential response of IgM and IgG memory B cell populations to CD40L: insights of T cell – memory B cell interactions. Front. Immunol. 15, (2024).

91. Yang, X. et al. Single-cell analysis reveals multi-faceted features of B cell development, together with age-associated B cell subpopulations. *Commun*. Biol. 9, 240 (2026).

92. Kasibhatla, S. et al. DNA Damaging Agents Induce Expression of Fas Ligand and Subsequent Apoptosis in T Lymphocytes via the Activation of NF-κB and AP-1. Mol. Cell 1, 543–551 (1998).

93. Consonni, F., Gambineri, E. & Favre, C. ALPS, FAS, and beyond: from inborn errors of immunity to acquired immunodeficiencies. Ann. Hematol. 101, 469–484 (2022).

94. Bae, S. et al. Existence of blood circulating immune-cell clusters (CICs) comprising antigen-presenting cells and B cells. 2022.01.26.477834 Preprint at 10.1101/2022.01.26.477834 (2022).

95. Sable, R. et al. Proximity ligation assay to study protein–protein interactions of proteins on two different cells. Biotechniques 65, 149–157 (2018).

96. Armingol, E., Officer, A., Harismendy, O. & Lewis, N. E. Deciphering cell–cell interactions and communication from gene expression. Nat. Rev. Genet. 22, 71–88 (2021).

97. Stoeckius, M. et al. Simultaneous epitope and transcriptome measurement in single cells. Nat. Methods 14, 865–868 (2017).

98. Rodríguez-Ubreva, J. et al. Single-cell Atlas of common variable immunodeficiency shows germinal center-associated epigenetic dysregulation in B-cell responses. Nat. Commun. 13, 1779 (2022).

99. Rodríguez-Ubreva, J. et al. Single-cell Atlas of common variable immunodeficiency shows germinal center-associated epigenetic dysregulation in B-cell responses. Nat. Commun. 13, 1779 (2022).

100. Maglione, P. J. et al. IRAK-4 and MyD88 deficiencies impair IgM responses against T-independent bacterial antigens. Blood 124, 3561–3571 (2014).

101. Marasco, E. et al. B-cell activation with CD40L or CpG measures the function of B-cell subsets and identifies specific defects in immunodeficient patients. Eur. J. Immunol. 47, 131–143 (2017).

102. Steiner, S., Wittke, K., Bauer, S., Scheibenbogen, C. & Hanitsch, L. G. Antibody Secretion Capacity in CVID Patients: Immunoglobulin Isotypes and Antigen Specificities After T-Cell-Dependent In Vitro Stimulation. J. Clin. Med. 14, (2025).

