## Supplementary material for "Stimulation strategy matters: A B-cell focused single-cell transcriptomic study on *in vitro* stimulated immune cells": This document contains supplementary tables and figures referred to in the manuscript.

### Sup. Table S1: Composition of stimulation mixes

| Mix | Stimulation component | Concentration | Factory information |
|---|---|---|---|
| 1 | ODN2006 | 0.5 µg/200 µL | ENZO Life sciences, ALX-746-006-C100 |
| 2 | MegaCD40L | 0.1 µg/mL | ENZO Life Sciences, ALX-522-110-C010 |
| | IL-4 | 50 ng/mL | tebu-bio, 200-04-A |
| | IL-21 | 50 ng/mL | tebu-bio, 200-21-B |
| 3 | Dynabeads (anti-CD3/CD28) | 20 µL/million PBMCs | Thermo Fisher Scientific |
| 4 | IFN-γ | 50 ng/mL | R&D systems, 285-IF |
| | R-848 | 5 µg/mL | InvivoGen, tlrl-r848 |
| | IL-4 | 50 ng/mL | tebu-bio, 200-04-A |
| | IL-21 | 50 ng/mL | tebu-bio, 200-21-B |
| | Anti-IgM | 5 µg/mL | Southern Biotech, 202201 |

### Sup. Table S2 Antibody panel used for FACS staining and acquisition for the FACS workflow

| Labeled antibody | Clone | Volume per sample (µL) | Factory information |
|---|---|---|---|
| **CD69-PE** | FN50 | 1.5 | BioLegend, 310906 |
| **CD86-BV711** | FUN-1 | 1.5 | BD Biosciences, 563158 |
| **CD40-APC** | 5C3 | 5 | BD Biosciences, 555591 |
| **CD80-FITC** | L307.4 | 5 | BD Biosciences, 557226 |
| **CD83-BV421** | HB15e | 1 | BD Biosciences, 562630 |
| **CD45-PacO** | HI30 | 1 | Life Technologies, MHCD4530 |
| **CD19-PECy7** | J3-119 | 5 | Analis, 134668 |
| **CD38-BV605** | HB7 | 2.5 | BD Biosciences, 562665 |
| **CD27-PerCPCy5.5** | M-T271 | 1.5 | BD Biosciences, 560612 |
| **CD21-R718** | B-ly4 | 2.5 | BD Biosciences, 752117 |
| **IgD-APC-H7** | IA6-2 | 2.5 | BD Biosciences, 561305 |

**Sup. Table S3: Overview of the cell hashing, FACS sorting and sample pooling for the single cell RNA-sequencing workflow.**

| Sample name | Healthy donor | Condition | Incubation time | Sorter | Sorted B cells | HTO | Fraction |
|---|---|---|---|---|---|---|---|
| **CEV017_1** | HC1 | Unstimulated | 8h | S6 | 92907 | TotalSeq-C0252 | 0.20 |
| **CEV017_2** | HC3 | Unstimulated | 8h | Vulcan | 43835 | TotalSeq-C0253 | 0.09 |
| **CEV017_3** | HC4 | Unstimulated | 8h | Vulcan | 55360 | TotalSeq-C0254 | 0.12 |
| **CEV017_4** | HC1 | MegaCD40L | 8h | Vulcan | 100498 | TotalSeq-C0255 | 0.21 |
| **CEV017_5** | HC3 | MegaCD40L | 8h | Vulcan | 30722 | TotalSeq-C0256 | 0.07 |
| **CEV017_6** | HC2 | IFNgamma | 8h | Melody | 12930 | TotalSeq-C0257 | 0.03 |
| **CEV017_7** | HC4 | IFNgamma | 8h | Melody | 13138 | TotalSeq-C0258 | 0.03 |
| **CEV017** | | | | | **349390** | | |
| **CEV018_1** | HC1 | ODN2006 | 8h | S6 | 120000 | TotalSeq-C0252 | 0.21 |
| **CEV018_2** | HC2 | ODN2006 | 8h | Vulcan | 18171 | TotalSeq-C0253 | 0.03 |
| **CEV018_3** | HC4 | ODN2006 | 8h | Vulcan | 71261 | TotalSeq-C0254 | 0.13 |
| **CEV018_4** | HC2 | MegaCD40L | 8h | Vulcan | 36397 | TotalSeq-C0255 | 0.06 |
| **CEV018_5** | HC2 | CD3/CD28 beads | 8h | Vulcan | 47946 | TotalSeq-C0256 | 0.08 |
| **CEV018_6** | HC3 | CD3/CD28 beads | 8h | Melody | 84480 | TotalSeq-C0257 | 0.15 |
| **CEV018_7** | HC4 | CD3/CD28 beads | 8h | Melody | 92375 | TotalSeq-C0258 | 0.16 |
| **CEV018** | | | | | **470630** | | |
| **CEV019_1** | HC2 | Unstimulated | 8h | Vulcan | 13218 | TotalSeq-C0253 | 0.02 |
| **CEV019_2** | HC3 | ODN2006 | 8h | Vulcan | 38787 | TotalSeq-C0254 | 0.06 |
| **CEV019_3** | HC4 | MegaCD40L | 8h | Melody | 66348 | TotalSeq-C0255 | 0.11 |
| **CEV019_4** | HC1 | CD3/CD28 beads | 8h | Melody | 120000 | TotalSeq-C0256 | 0.19 |
| **CEV019_5** | HC1 | IFNgamma | 8h | Melody | 120000 | TotalSeq-C0257 | 0.19 |
| **CEV019_6** | HC3 | IFNgamma | 8h | Melody | 33339 | TotalSeq-C0258 | 0.05 |
| **CEV019** | | | | | **391692** | | |
| **CEV020_1** | HC1 | Unstimulated | 30h | Melody | 120000 | TotalSeq-C0252 | 0.16 |
| **CEV020_2** | HC3 | Unstimulated | 30h | Melody | 120000 | TotalSeq-C0253 | 0.16 |
| **CEV020_3** | HC4 | Unstimulated | 30h | Melody | 120000 | TotalSeq-C0254 | 0.16 |
| **CEV020_4** | HC1 | MegaCD40L | 30h | S6 | 120000 | TotalSeq-C0255 | 0.16 |
| **CEV020_5** | HC3 | MegaCD40L | 30h | S6 | 120000 | TotalSeq-C0256 | 0.16 |
| **CEV020_6** | HC2 | IFNgamma | 30h | Vulcan | 38602 | TotalSeq-C0257 | 0.05 |
| **CEV020_7** | HC4 | IFNgamma | 30h | Vulcan | 94541 | TotalSeq-C0258 | 0.13 |
| **CEV020** | | | | | **733143** | | |
| **CEV021_1** | HC1 | ODN2006 | 30h | Melody | 120000 | TotalSeq-C0252 | 0.18 |
| **CEV021_2** | HC2 | ODN2006 | 30h | Melody | 72863 | TotalSeq-C0253 | 0.11 |
| **CEV021_3** | HC4 | ODN2006 | 30h | Melody | 117150 | TotalSeq-C0254 | 0.17 |
| **CEV021_4** | HC2 | MegaCD40L | 30h | S6 | 56937 | TotalSeq-C0255 | 0.08 |
| **CEV021_5** | HC2 | CD3/CD28 beads | 30h | S6 | 86033 | TotalSeq-C0256 | 0.13 |
| **CEV021_6** | HC3 | CD3/CD28 beads | 30h | Vulcan | 111150 | TotalSeq-C0257 | 0.16 |
| **CEV021_7** | HC4 | CD3/CD28 beads | 30h | Vulcan | 120000 | TotalSeq-C0258 | 0.18 |
| **CEV021** | | | | | **684133** | | |
| **CEV022_1** | HC2 | Unstimulated | 30h | S6 | 101676 | TotalSeq-C0253 | 0.16 |
| **CEV022_2** | HC3 | ODN2006 | 30h | S6 | 95799 | TotalSeq-C0254 | 0.15 |
| **CEV022_3** | HC4 | MegaCD40L | 30h | Vulcan | 120000 | TotalSeq-C0255 | 0.19 |
| **CEV022_4** | HC1 | CD3/CD28 beads | 30h | Vulcan | 120000 | TotalSeq-C0256 | 0.19 |
| **CEV022_5** | HC1 | IFNgamma | 30h | Vulcan | 120000 | TotalSeq-C0257 | 0.19 |
| **CEV022_6** | HC3 | IFNgamma | 30h | Vulcan | 68228 | TotalSeq-C0258 | 0.11 |
| **CEV022** | | | | | **625703** | | |

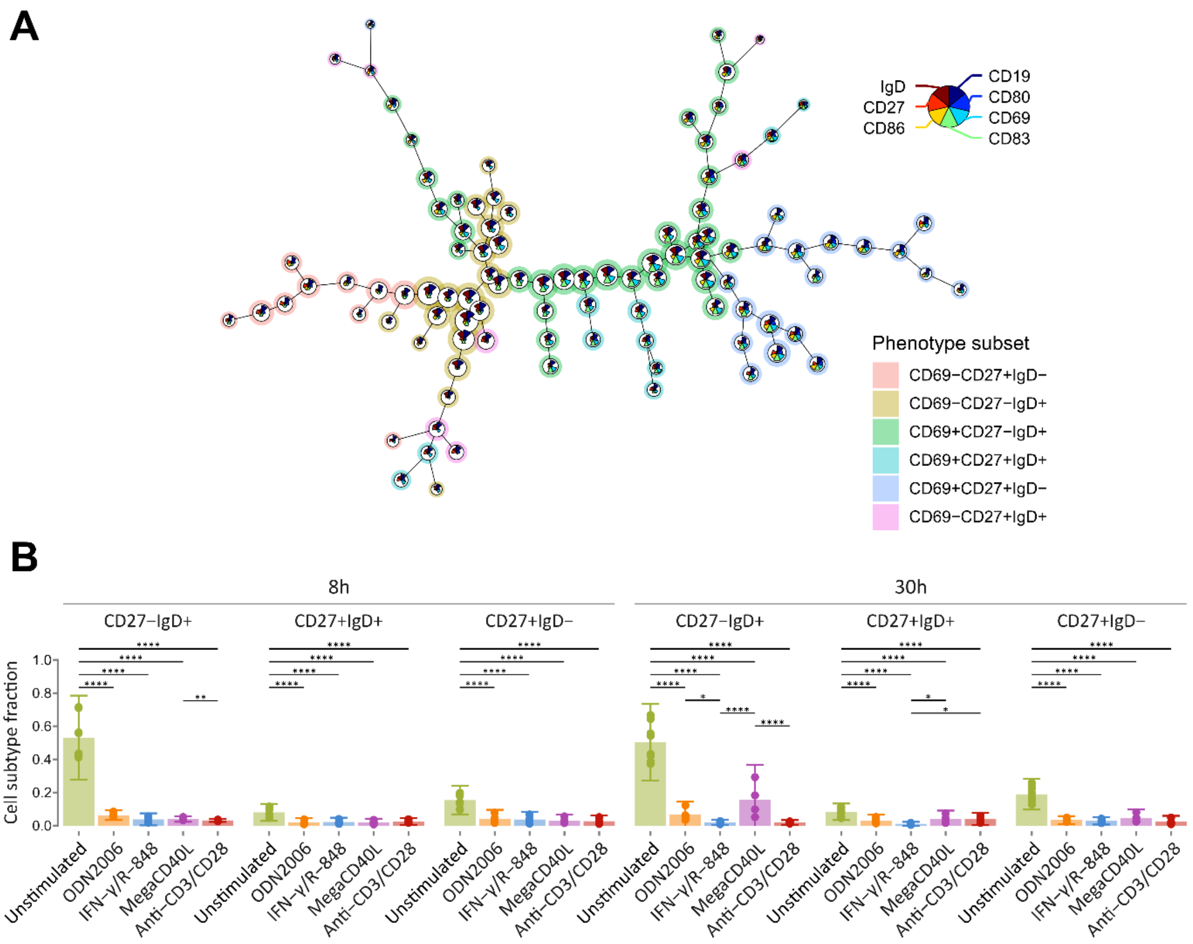


**Sup. Figure S1: FlowSOM tree and inactivated fraction of in vitro stimulated CD19+ cells.** (**A**) FlowSOM model (100 clusters and 15 metaclusters with markers CD19, CD27, CD69, CD80, CD83, CD86, IgD) generated using $CD45^{+}CD19^{+}$ *in vitro* stimulated cells. B-cell maturation markers (CD27 and IgD) were used for cell subset annotation, while CD69 distinguished activated from inactivated cells. (**B**) The fraction of inactivated ($CD69^{-}$) B cells ($CD19^{+}$) per healthy donor, stimulation condition, and incubation time point over the B cell subsets. $CD27^{-}IgD^{+}$ population represent pre-germinal center B cells. Unswitched and switched memory B cells are characterized by $CD27^{+}IgD^{+}$ and $CD27^{+}IgD^{-}$ phenotypes, respectively. Error bars representing 95.5% (mean ± 2*standard deviation) confidence intervals. Dots represent individual cell type fractions per healthy donor. Significant differences are shown, ****p<0.0001, ***p<0.001, **p<0.01, *p<0.05, generalized linear mixed model, beta-family regression using a logic link function. Non-significant differences are not indicated.

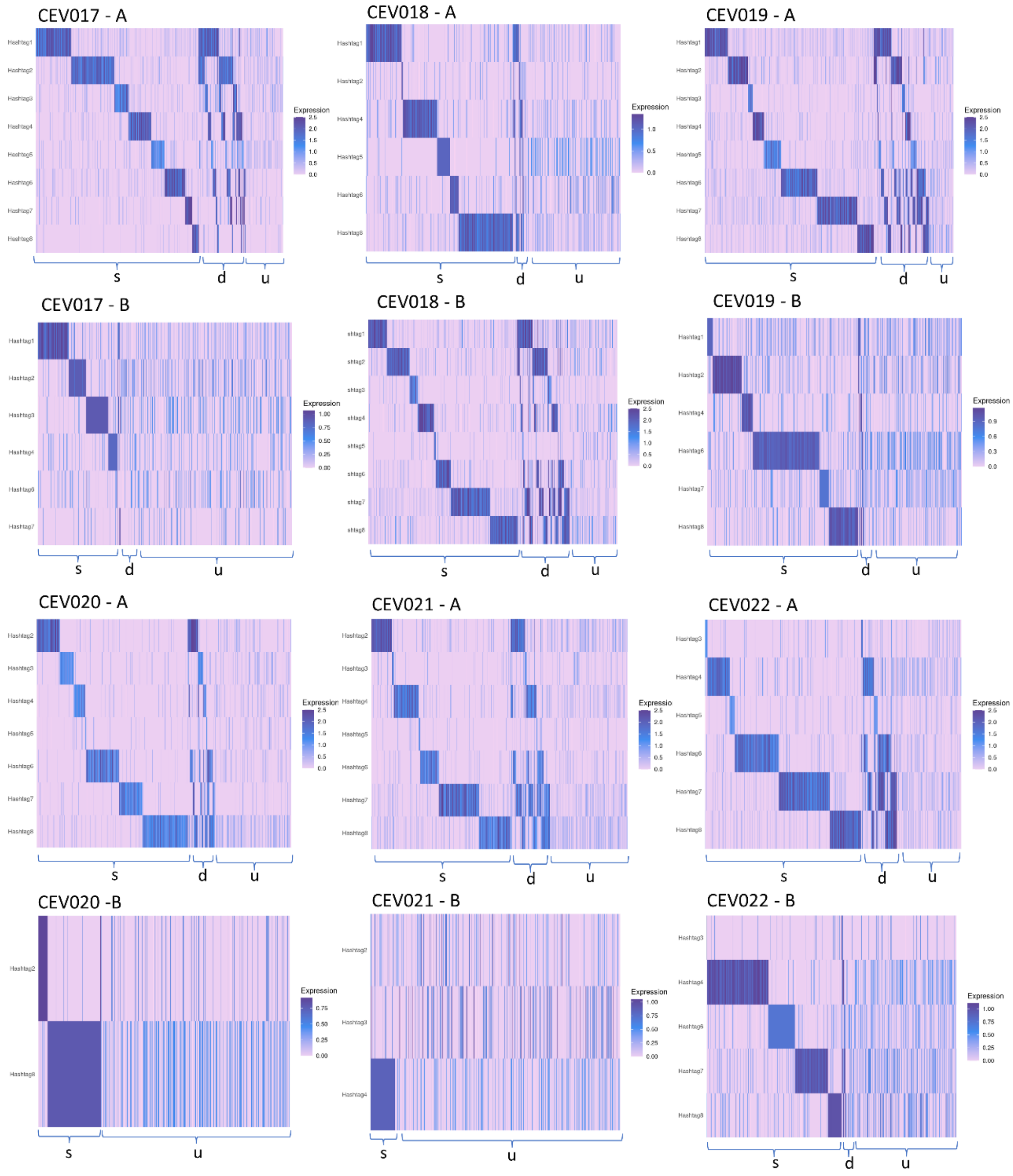


**Sup. Figure S2: Sample demultiplexing using cell hashtags (HTO) in CEV017-CEV022.** The HTO counts associated to the hashtags used to distinguish between samples in the pool are shown for all cells (panels A). Singlets (s), doublets (d) and unidentified (u) cells can be distinguished based on the HTO expression. The heatmaps obtained after an additional demultiplexing step was performed on the unidentified cells (panels B). Singlet cells were isolated resulting in higher number of retained cells after demultiplexing.

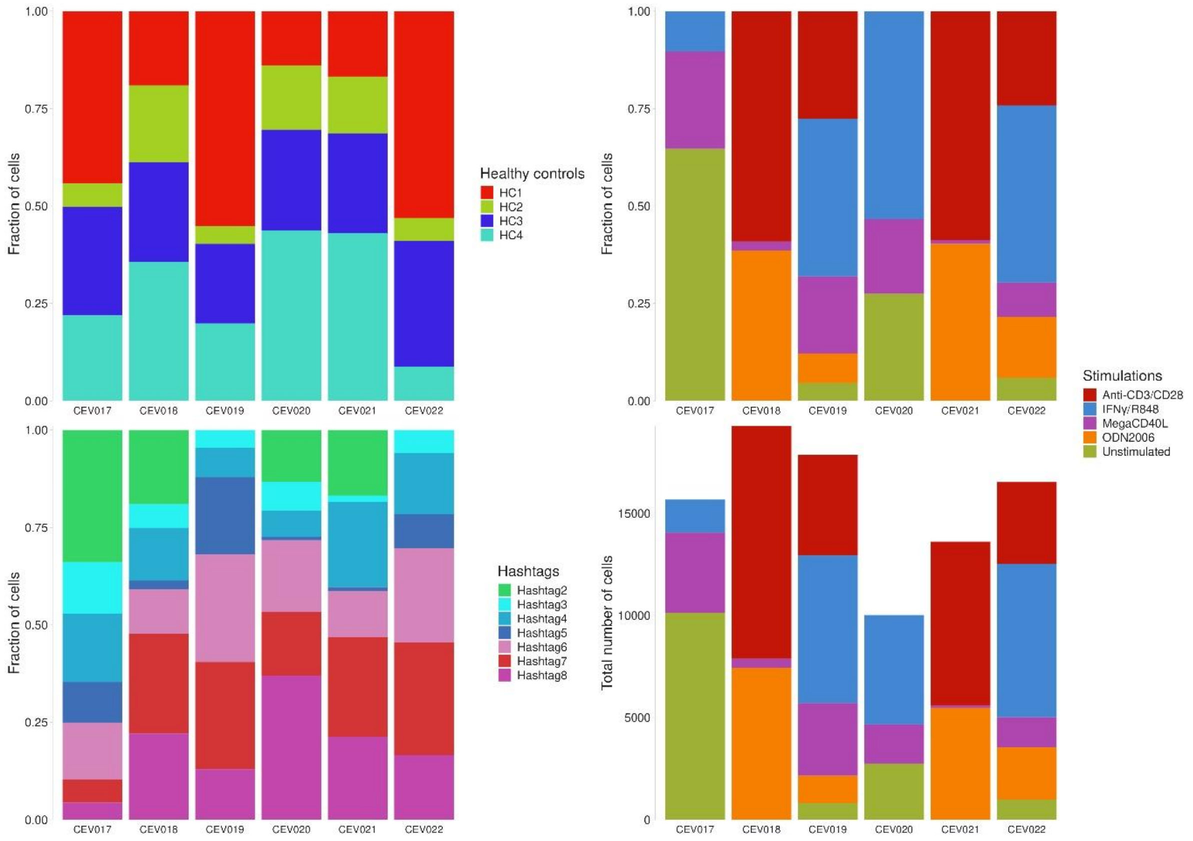


**Sup. Figure S3: Fractions of singlets per pool originating from a given healthy donor, associated to a stimulatory condition or hashtag.** The bottom right panel shows the number of total cells per pool and per stimulatory condition.

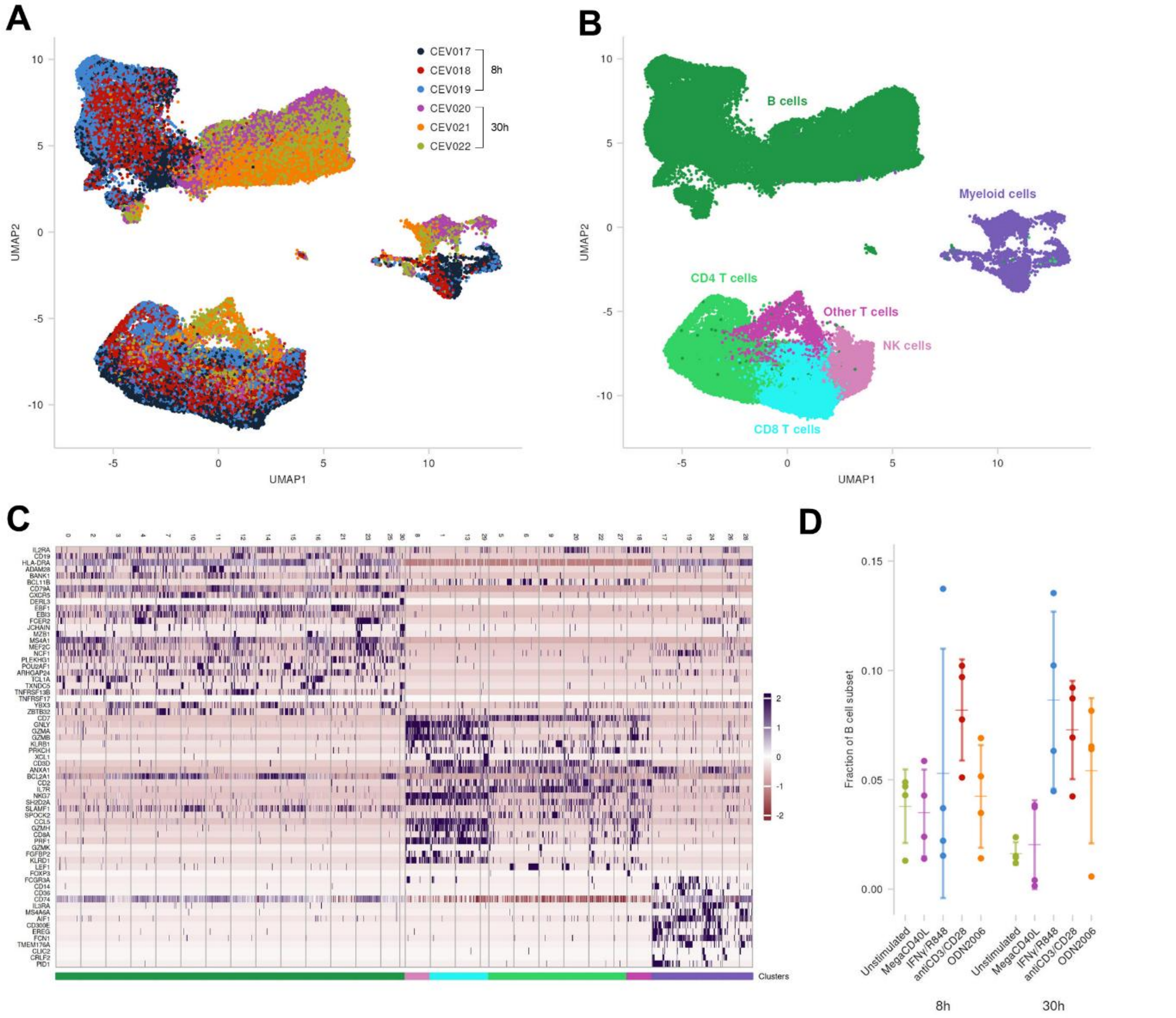


**Sup. Figure S4: Single-cell transcriptomic analysis of the final integrated data set. (A-B)** UMAP embedding of all cells (n = 81,160) derived from the integrated dataset of six samples pools (including samples from two different incubation time points, four healthy controls and five stimulatory conditions). Cells are colored by the sample pools (CEV017-CEV022) (**A**) and cell type (**B**). (**C)** Heatmap shows the expression levels of differential expressed (DE) genes in each cluster identified (n=30). The gene expression levels were used to annotate on the level of the major leucocyte cell subsets (i.e., T, B, NK and innate cells) based on expression levels of cell-specific genes, such as CD19, HLA-DR, and CD79A for the B cells (dark green), CD7, CD3D, CD8A, PRF1, and GZMA for the T (blue and light green) and NK (light pink) cells, and FCN1, CD14, CD300E, AIF1, and PID1 for the innate cell, including monocyte and dendritic cell subsets (purple) (**D**) Dot plot of fraction of the B cells per stimulatory condition and incubation time point. Each dot represents a healthy control sample. 95% confidence intervals and the mean fraction of B cells per stimulatory condition were calculated per time point represented as error bars.

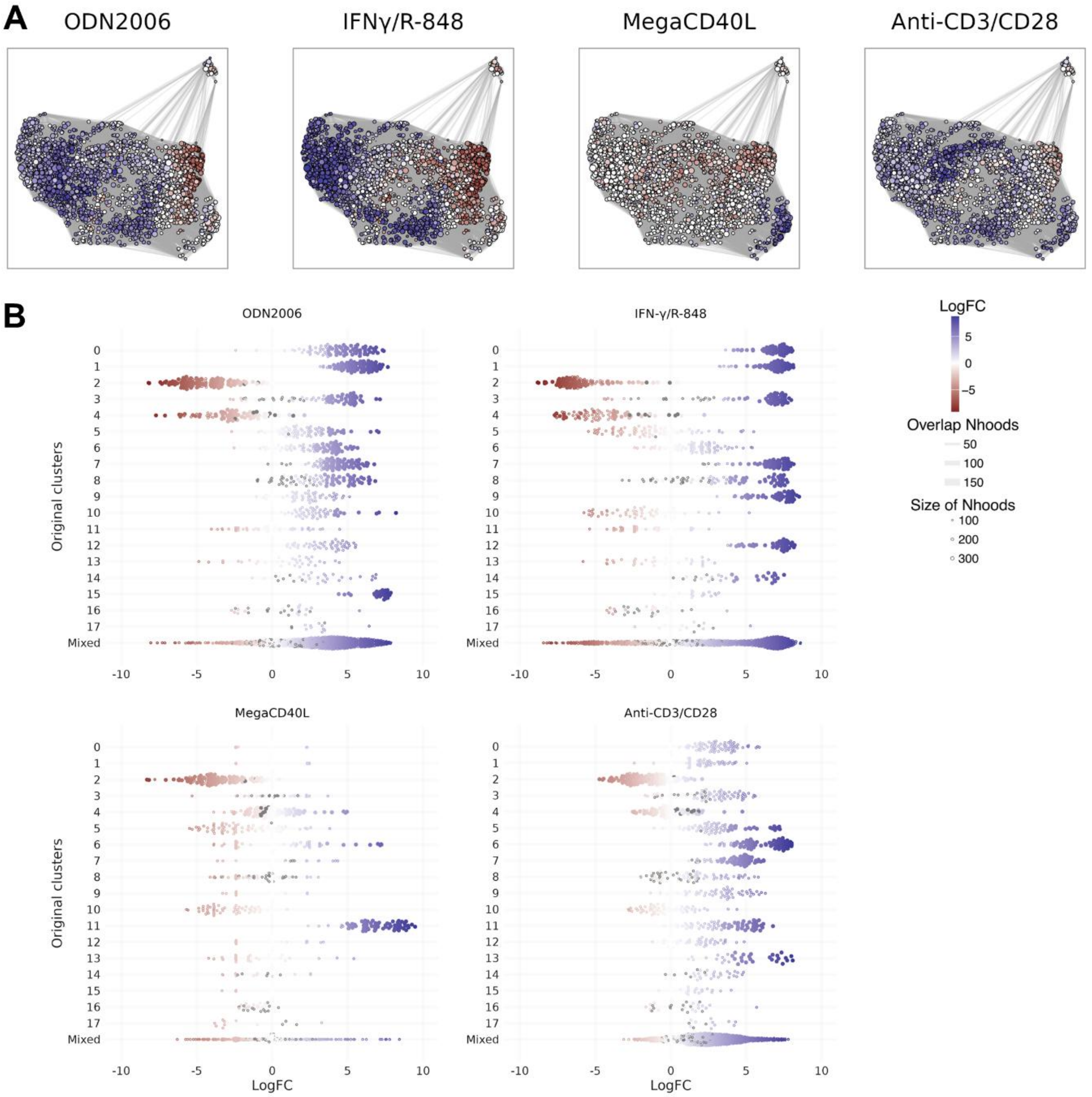


**Sup. Figure S5: Neighbourhood analysis and differential abundance testing for B-cell subset stimulated for 30h. (A)** Neighbourhood (Nhood) graph based on the UMAP visualization of the cells stimulated for 30h per stimulatory condition, show the results of the differential abundance testing with unstimulated cells as reference using MiloR. Each Nhood is represented as a dot of which the size is correlated to the number of cells. The Nhoods are colored based on LogFC values of differential abundance testing: red indicates increased abundance of unstimulated cells, blue indicates increased abundance of stimulated cells and white indicate no significant differences in abundance. (**B**) Beeswarm plots for 30 hours of stimulation showing the distribution of LogFC after abundance testing per Nhood (dots) per Louvain cell cluster (y-axis), performed for each of the four stimulations separately. Full colored dots represent the Nhoods selected for downstream analysis, same coloring scheme was used as described in panel **A**.

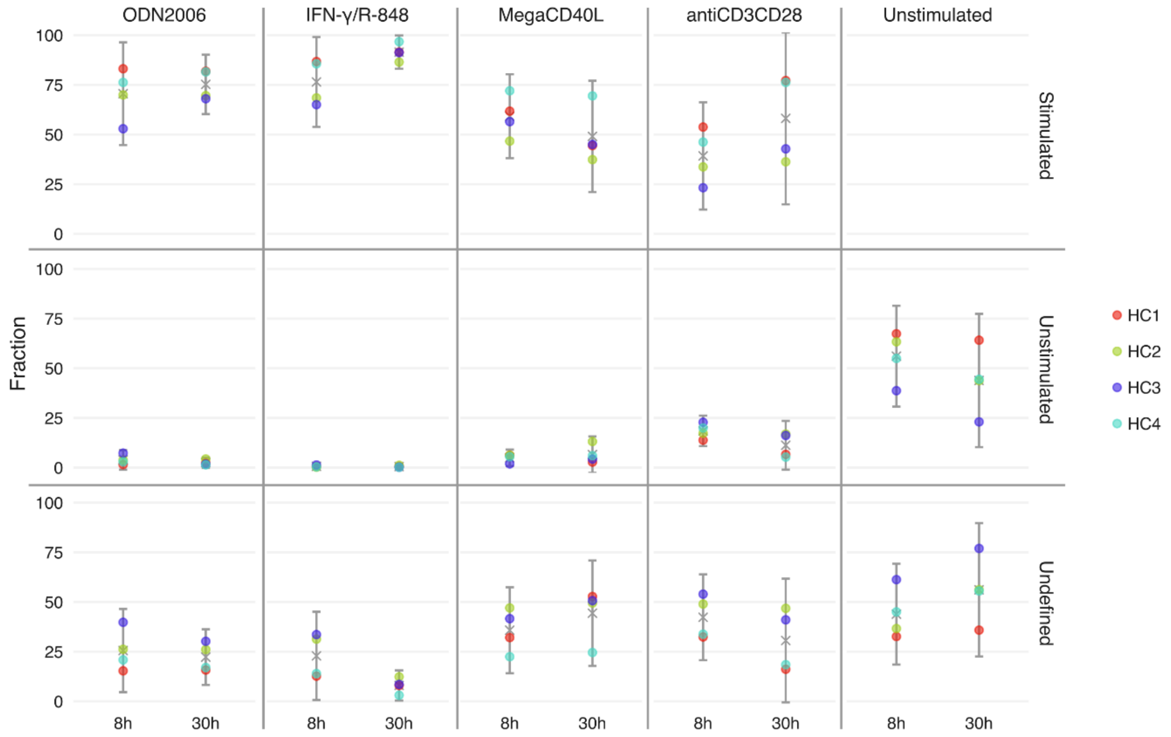


**Sup. Figure S6*: In-depth comparison of the refined stimulation status annotations based on neighbourhood (Nhood) selection (columns) with the original metadata-based annotation (rows) of the cells per stimulatory condition, incubation time point and healthy donor.*** Fractions were calculated relative to the original (metadata-based) number of cells associated with a specific stimulatory condition, with respect to the healthy donor and incubation time point of the sample. The errorbars show the 95.5% confidence interval (mean+/- 2*sd). Dots are colored with respect to the healthy control of which the cells originate from.

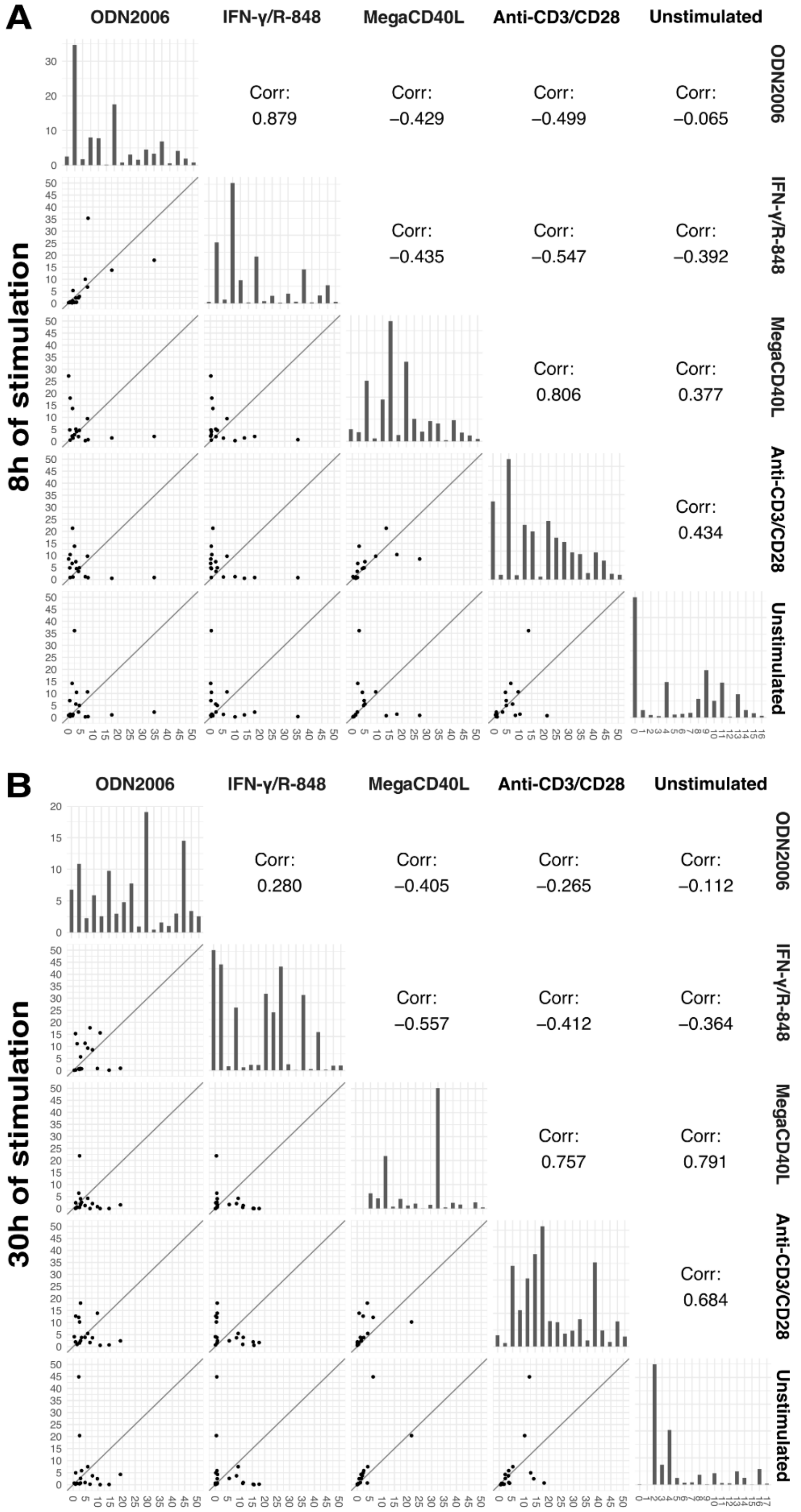


**Sup. Figure S7: Correlation plot of fractions (in percent) per condition within Louvain clusters identified in the distinct time point subsets.** The scatter plots in the lower triangle visualize the fractions within each of the seventeen (8h, **A**) or eighteen (30h, **B**) clusters, split over the five conditions. On the diagonal, a bar plot shows the fractions (vertical axis) of a specific condition within each of these clusters (horizontal axis). The upper triangle shows the Spearman correlation of the fractions shown in the scatter plots. This is shown for both 8h (**A**) and 30h (**B**) of stimulation.

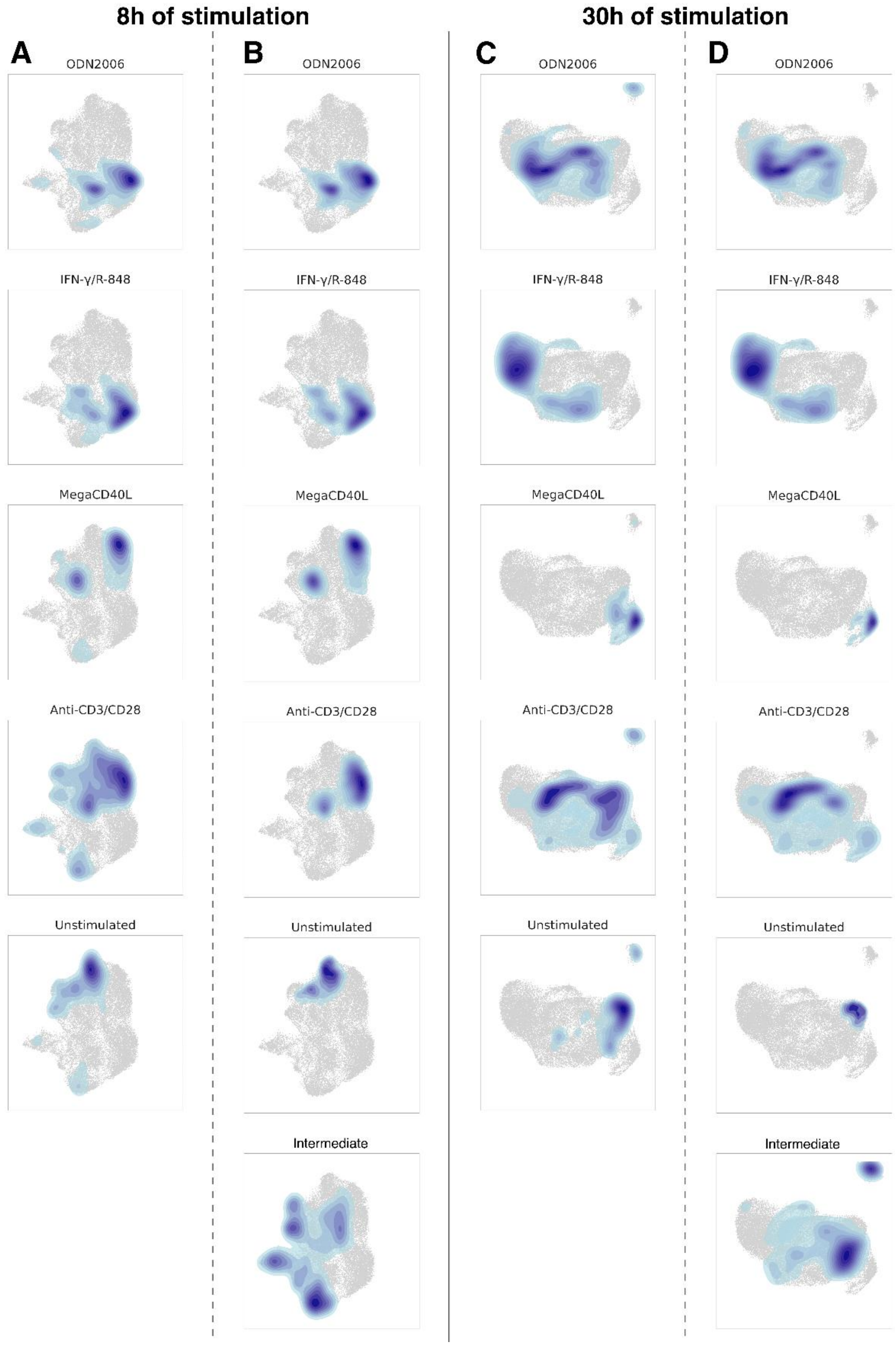


**Sup. Figure S8: Density plots of the annotation of cells regarding the stimulatory conditions.** UMAP-visualizations are shown for cells stimulated for 8h (columns **A**, **B**) and 30h (columns **C**, **D**). The contour lines show the density of the cells per stimulatory condition: the darker the indicated region, the higher the abundance of cells annotated with a specific condition as mentioned above each of the plots separately; uncolored regions did not contain cells with the annotation of interest. Columns **A** and **C** indicate the density plots based on the cell annotation using the original metadata (original sample annotation of the experiment), while columns **B** and **D** visualize the density plots using the refined stimulation status annotation based on the selected Nhoods per stimulatory condition and time point.

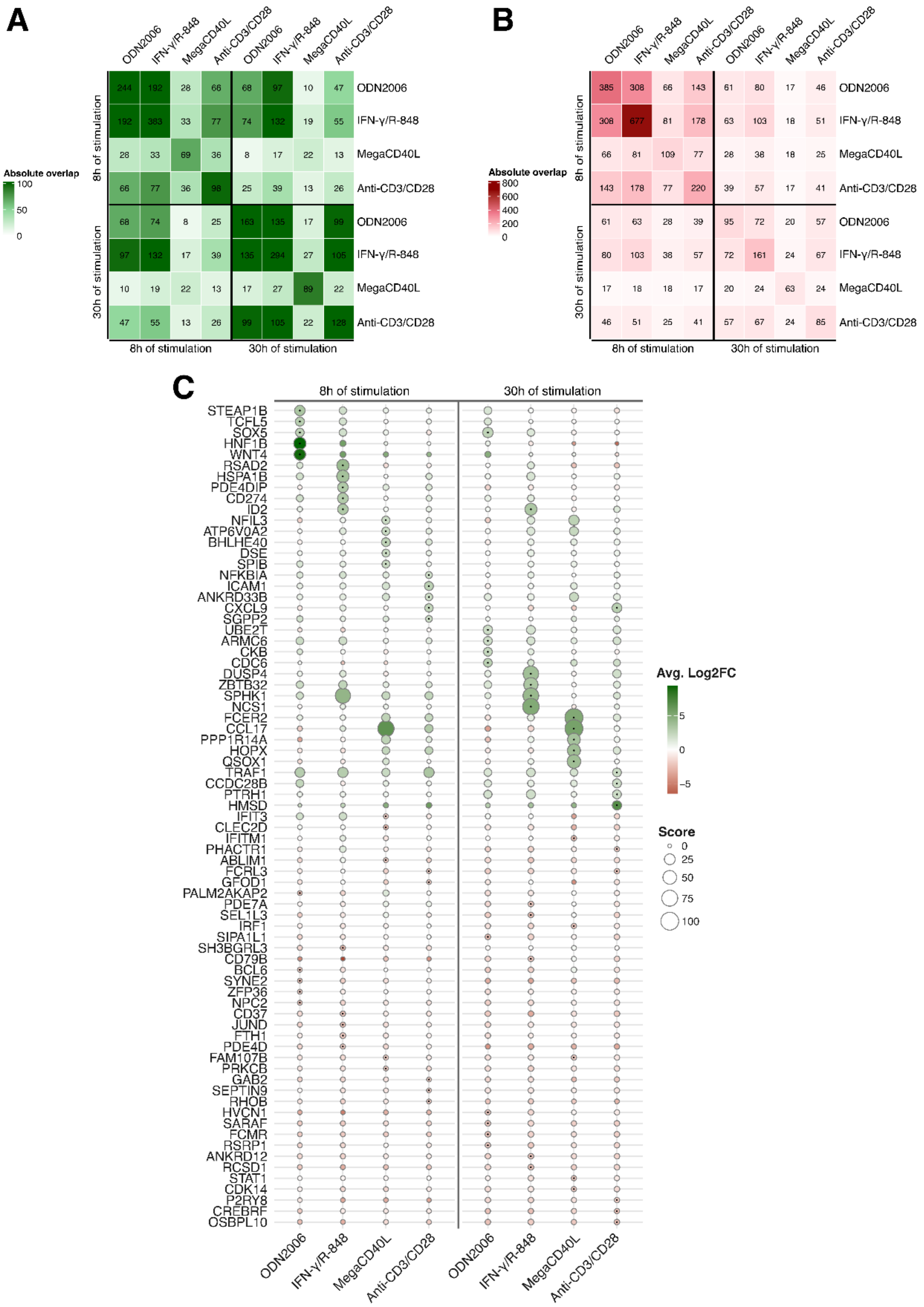


**Sup. Figure S9: Overlapping and unique differentially expressed genes per stimulatory condition and incubation time point.** (**A-B**) Heatmap showing the overlap in significant differentially expressed up- (**A**) and downregulated (**B**) genes between conditions and incubation time points. The diagonal element represents the number of differentially expressed genes per combination of stimulatory condition and incubation time point, while the off-diagonal elements represent the overlap (**C**) Dot plot representing the gene expression levels of the top five up- and downregulated genes uniquely differentially expressed in a specific stimulatory condition and at a specific incubation time point. The size of the dots represents the absolute gene score of each gene (see M&M section for detailed information). The presence of a small dot in the center of the larger dots, indicates that the gene is one of the top five differentially expressed genes in that stimulation strategy.

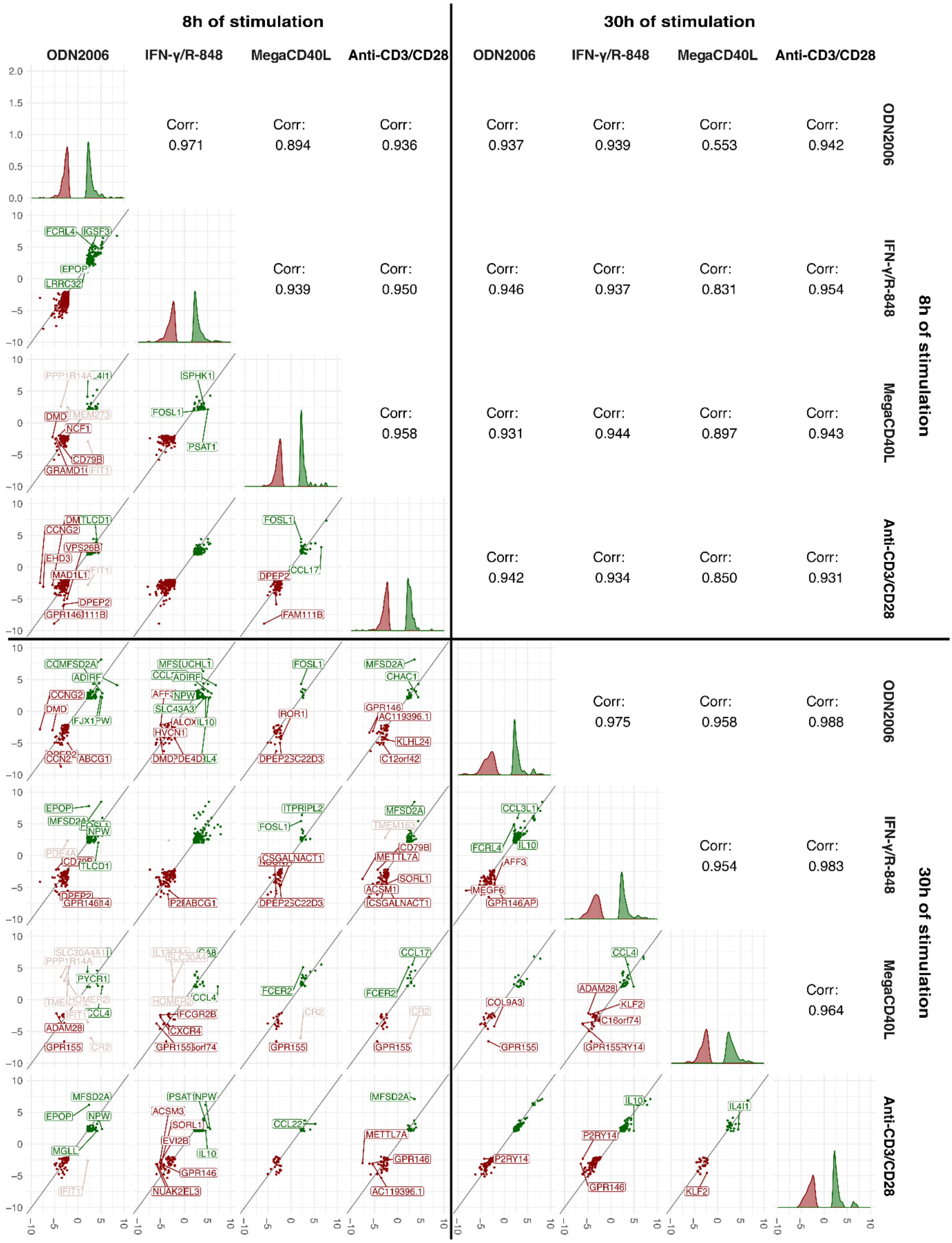


**Sup. Figure S10: Correlation plot of average log2FC expression values of overlapping genes between conditions and incubation time points.** The scatter plots in the lower triangle visualize the log2FC of the overlapping differentially expressed genes. Genes are colored regarding the (in)consistency in expression direction between the compared samples: upregulated (green), downregulated (red) and mixed (almond pink). Genes showing large deviations from the bisector (difference between samples in log2FC larger than 2) are shown with a label. On the diagonal, the density plots of the log2FC values per sample (stimulatory condition and incubation time point) are plotted. The upper triangle shows the Pearson correlation of the expression levels of the overlapping genes shown in the scatter plots.

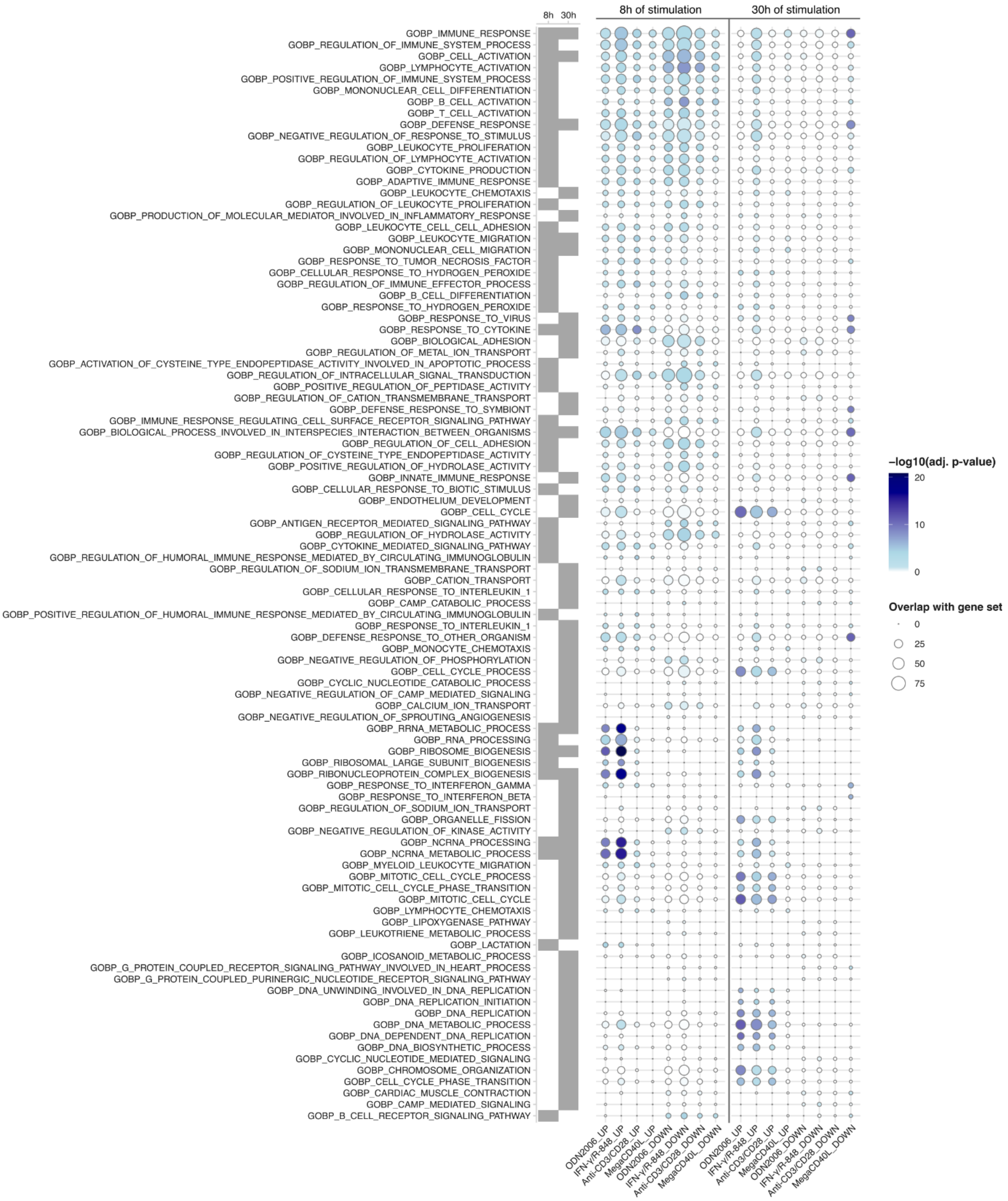


**Sup. Figure S11: Gene set enrichment analysis based on Gene Ontology Biological Process (GOBP) gene sets.** The top 10 significantly enriched GOBP (vertical axis) per combination of stimulatory condition, incubation time point and direction of expression (UP for upregulation and DOWN for downregulation, horizontal axis) are shown in this dot plot. The size of the dots represents the overlap of differentially expressed genes found in the GOBP gene set of interest, while the color indicated the adjusted p-value after hypergeometric statistical testing of this overlap. The tile plot at the left of the graph indicates if the GOBP gene set was observed to be significantly enriched in at least one differentially expressed gene set after 8h, 30h or at both time points (grey).

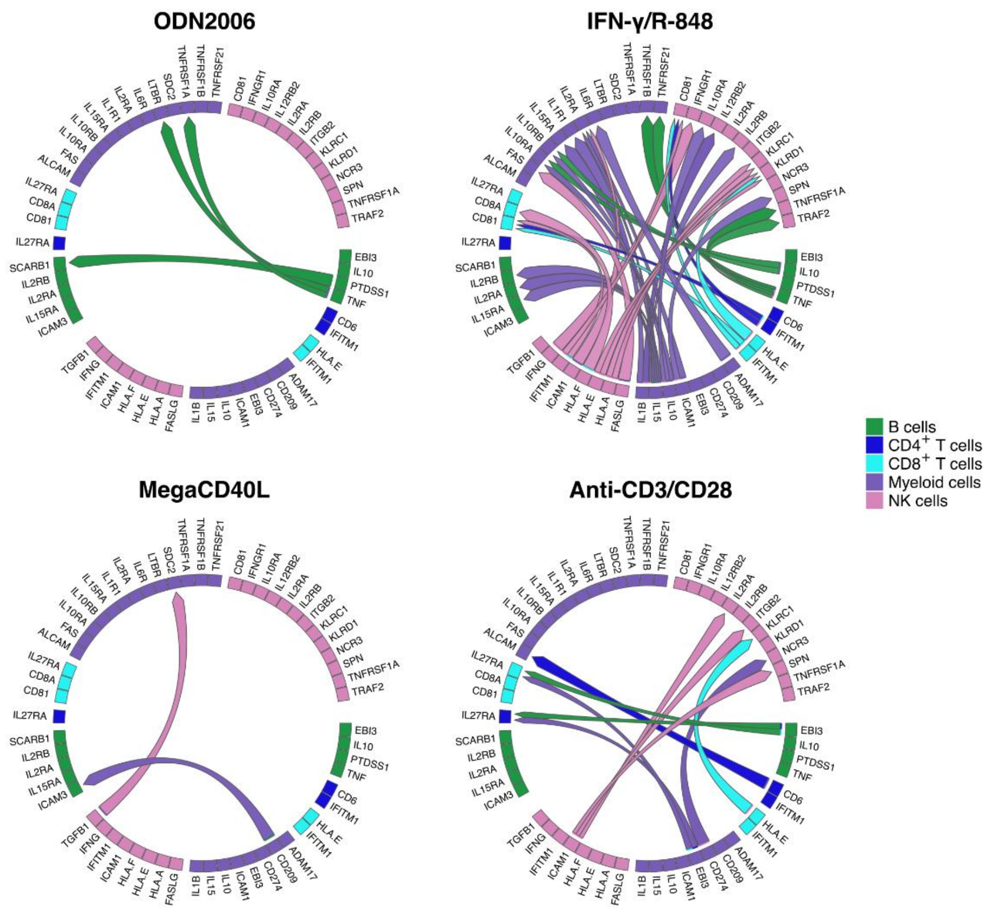


**Sup. Figure S12: Chord diagrams representing overall top 50 ligand-receptor interactions per stimulatory condition compared to the unstimulated samples 30h of stimulation.** The arrows, colored based on the sender cell type, connect the ligand in a specific sender cell type with the receptor in a specific receiver cell type. If arrows are absent between ligands and receptor, the combination of interaction and stimulatory condition was not present in the overall top 50 list for that specific condition.

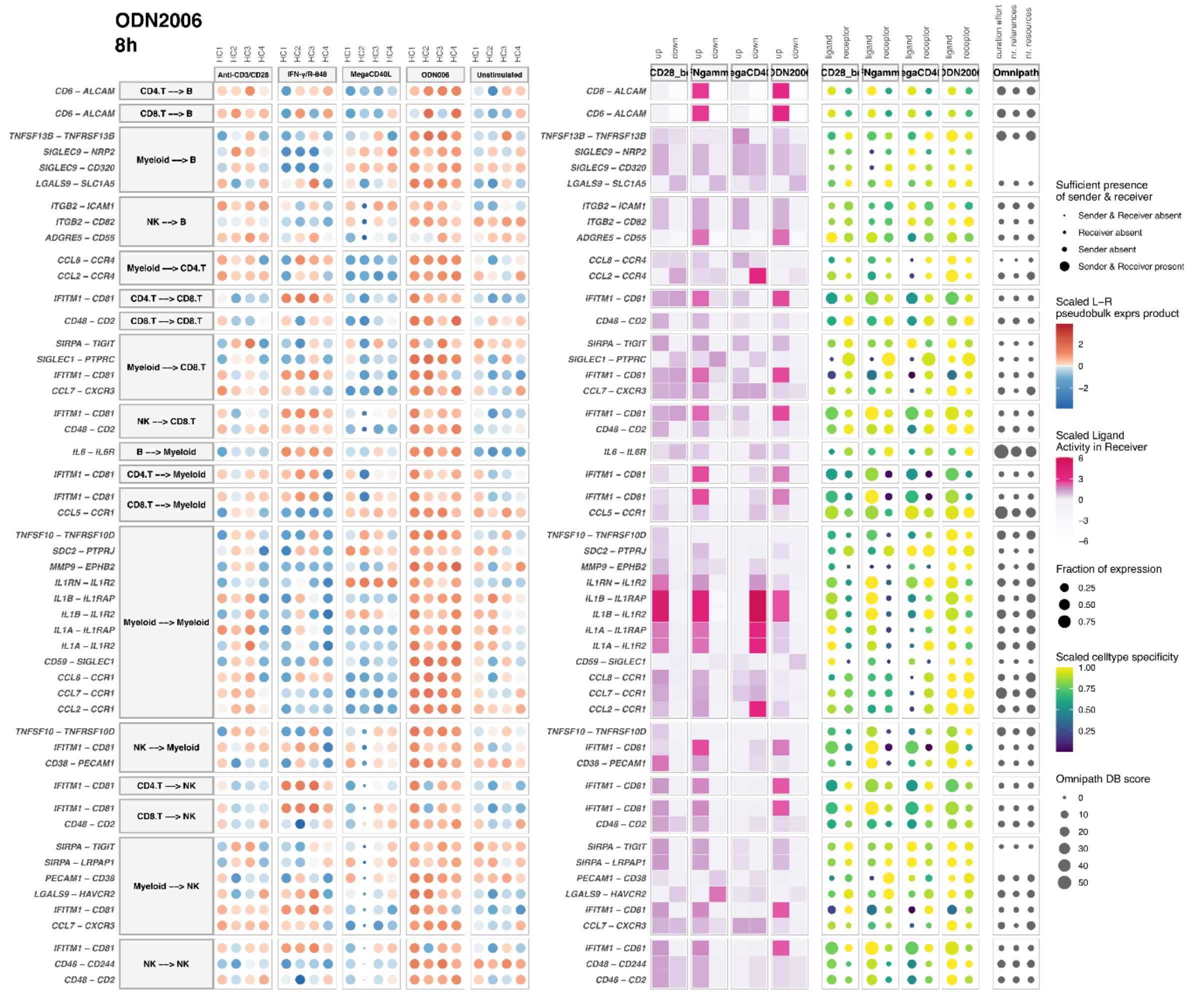

**Sup. Figure S13: Top 50 interactions observed in the cell-cell communication output of ODN2006 after 8h of stimulation, with the unstimulated samples as reference.** Cell-cell interactions through specific ligand-receptor pairs are shown at the vertical axis. The dot plot at the left shows the scaled (pseudobulk) expression product of the ligand-receptor (LR) combination per sample (HC1-HC4, horizontal axis). The size of the dots represents the presence or absence (after filtering, see Materials&Methods) of the sender and receiver cell type within the individual samples. The tile plot in the middle shows the scaled ligand activity in the receiver cell type, split over up- and down-regulated target genes of the ligand-receptor pair (horizontal axis). Higher ligand activities (dark pink) suggest functional interaction with an effect on the downstream target gene regulation in the receiver cell. The colored dot plot at the right shows the cell type specificity of the ligand and receptors present in the interactions. The size of the dots indicates the expression fraction of the ligands and the receptors in the sender and receiver cells, respectively. The dot plot at the left represents the score of the interaction with the Omnipath database (DB) as reference.

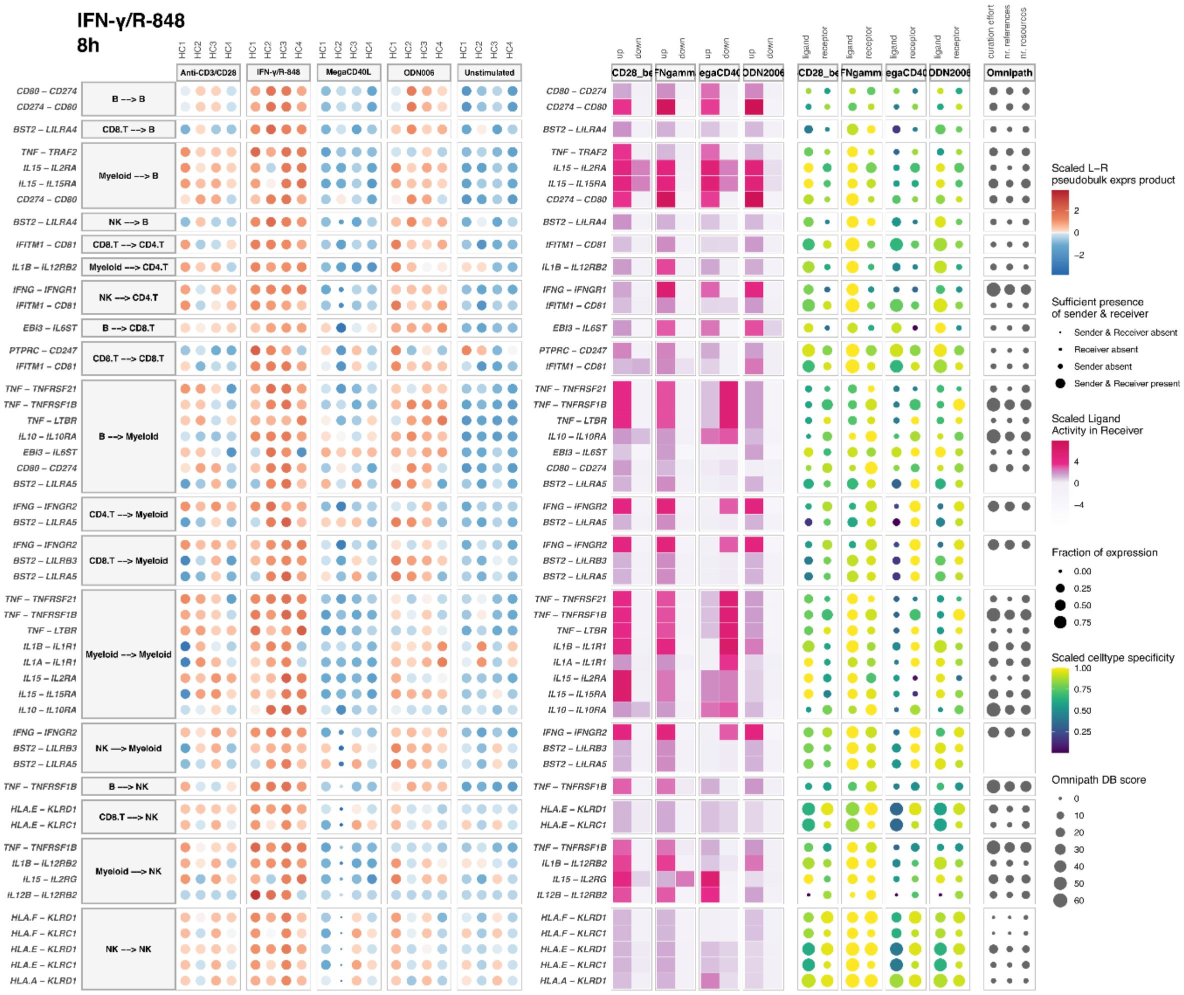


**Sup. Figure S14: Top 50 interactions observed in the cell-cell communication output of INF-γ/R-848 after 8h of stimulation, with the unstimulated samples as reference.** Cell-cell interactions through specific ligand-receptor pairs are shown at the vertical axis. The dot plot at the left shows the scaled (pseudobulk) expression product of the ligand-receptor (LR) combination per sample (HC1-HC4, horizontal axis). The size of the dots represents the presence or absence (after filtering, see Materials&Methods) of the sender and receiver cell type within the individual samples. The tile plot in the middle shows the scaled ligand activity in the receiver cell type, split over up- and down-regulated target genes of the ligand-receptor pair (horizontal axis). Higher ligand activities (dark pink) suggest functional interaction with an effect on the downstream target gene regulation in the receiver cell. The colored dot plot at the right shows the cell type specificity of the ligand and receptors present in the interactions. The size of the dots indicates the expression fraction of the ligands and the receptors in the sender and receiver cells, respectively. The dot plot at the left represents the score of the interaction with the Omnipath database (DB) as reference.

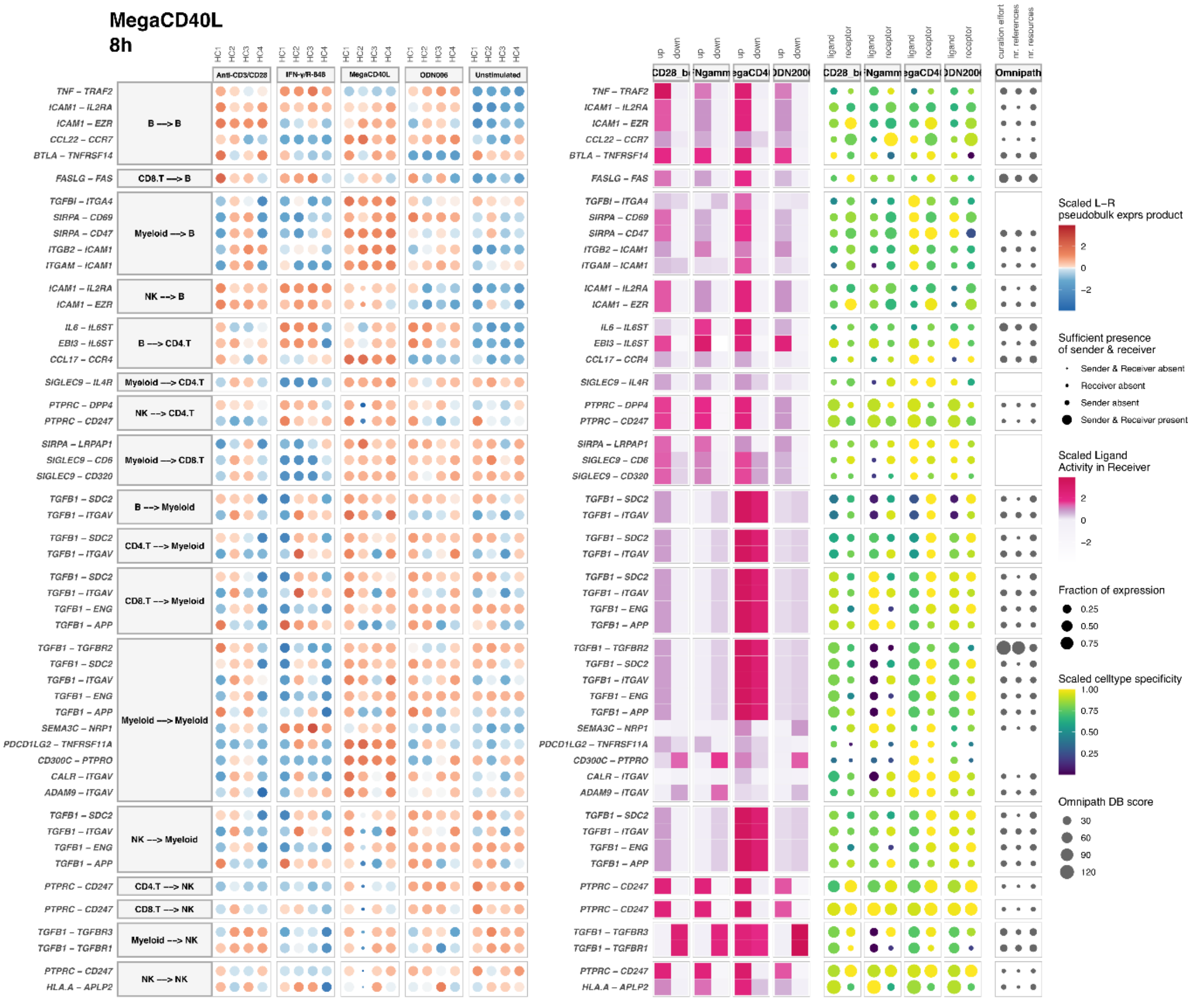


**Sup. Figure S15: Top 50 interactions observed in the cell-cell communication output of MegaCD40L after 8h of stimulation, with the unstimulated samples as reference.** Cell-cell interactions through specific ligand-receptor pairs are shown at the vertical axis. The dot plot at the left shows the scaled (pseudobulk) expression product of the ligand-receptor (LR) combination per sample (HC1-HC4, horizontal axis). The size of the dots represents the presence or absence (after filtering, see Materials&Methods) of the sender and receiver cell type within the individual samples. The tile plot in the middle shows the scaled ligand activity in the receiver cell type, split over up- and down-regulated target genes of the ligand-receptor pair (horizontal axis). Higher ligand activities (dark pink) suggest functional interaction with an effect on the downstream target gene regulation in the receiver cell. The colored dot plot at the right shows the cell type specificity of the ligand and receptors present in the interactions. The size of the dots indicates the expression fraction of the ligands and the receptors in the sender and receiver cells, respectively. The dot plot at the left represents the score of the interaction with the Omnipath database (DB) as reference.

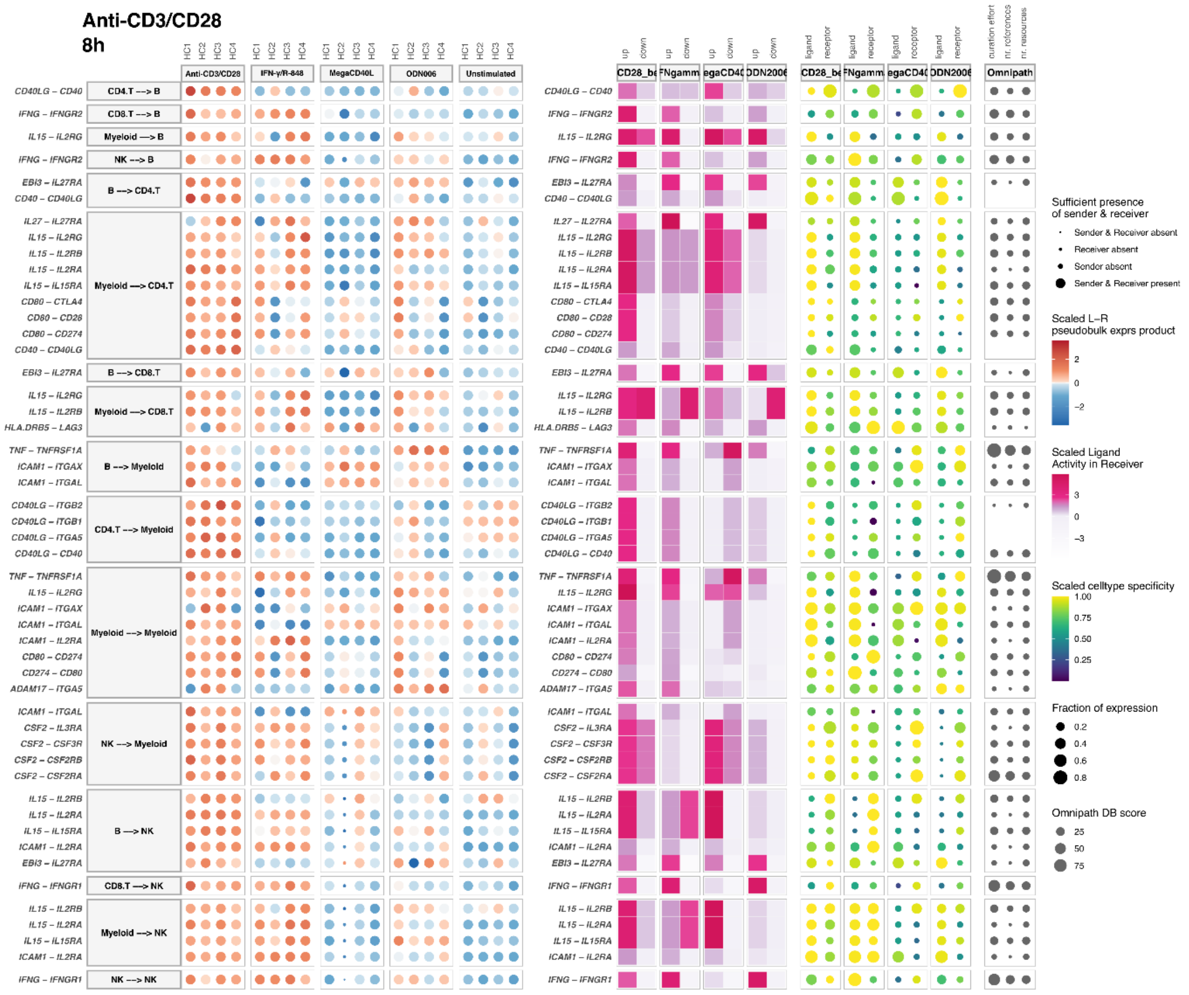


**Sup. Figure S16: Top 50 interactions observed in the cell-cell communication output of anti-CD3/CD28 after 8h of stimulation, with the unstimulated samples as reference.** Cell-cell interactions through specific ligand-receptor pairs are shown at the vertical axis. The dot plot at the left shows the scaled (pseudobulk) expression product of the ligand-receptor (LR) combination per sample (HC1-HC4, horizontal axis). The size of the dots represents the presence or absence (after filtering, see Materials&Methods) of the sender and receiver cell type within the individual samples. The tile plot in the middle shows the scaled ligand activity in the receiver cell type, split over up- and down-regulated target genes of the ligand-receptor pair (horizontal axis). Higher ligand activities (dark pink) suggest functional interaction with an effect on the downstream target gene regulation in the receiver cell. The colored dot plot at the right shows the cell type specificity of the ligand and receptors present in the interactions. The size of the dots indicates the expression fraction of the ligands and the receptors in the sender and receiver cells, respectively. The dot plot at the left represents the score of the interaction with the Omnipath database (DB) as reference.

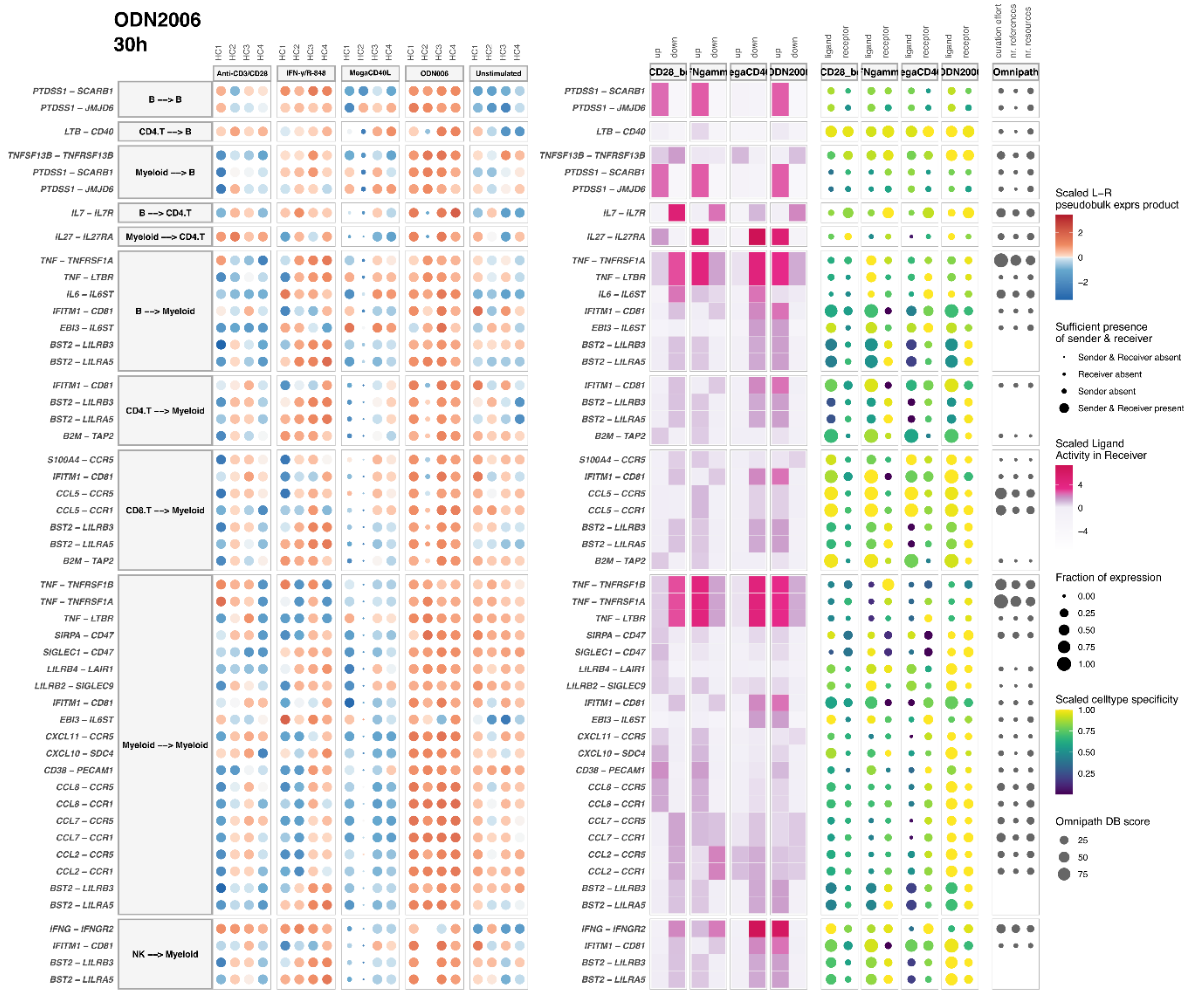


**Sup. Figure S17: Top 50 interactions observed in the cell-cell communication output of ODN2006 after 30h of stimulation, with the unstimulated samples as reference.** Cell-cell interactions through specific ligand-receptor pairs are shown at the vertical axis. The dot plot at the left shows the scaled (pseudobulk) expression product of the ligand-receptor (LR) combination per sample (HC1-HC4, horizontal axis). The size of the dots represents the presence or absence (after filtering, see Materials&Methods) of the sender and receiver cell type within the individual samples. The tile plot in the middle shows the scaled ligand activity in the receiver cell type, split over up- and down-regulated target genes of the ligand-receptor pair (horizontal axis). Higher ligand activities (dark pink) suggest functional interaction with an effect on the downstream target gene regulation in the receiver cell. The colored dot plot at the right shows the cell type specificity of the ligand and receptors present in the interactions. The size of the dots indicates the expression fraction of the ligands and the receptors in the sender and receiver cells, respectively. The dot plot at the left represents the score of the interaction with the Omnipath database (DB) as reference.

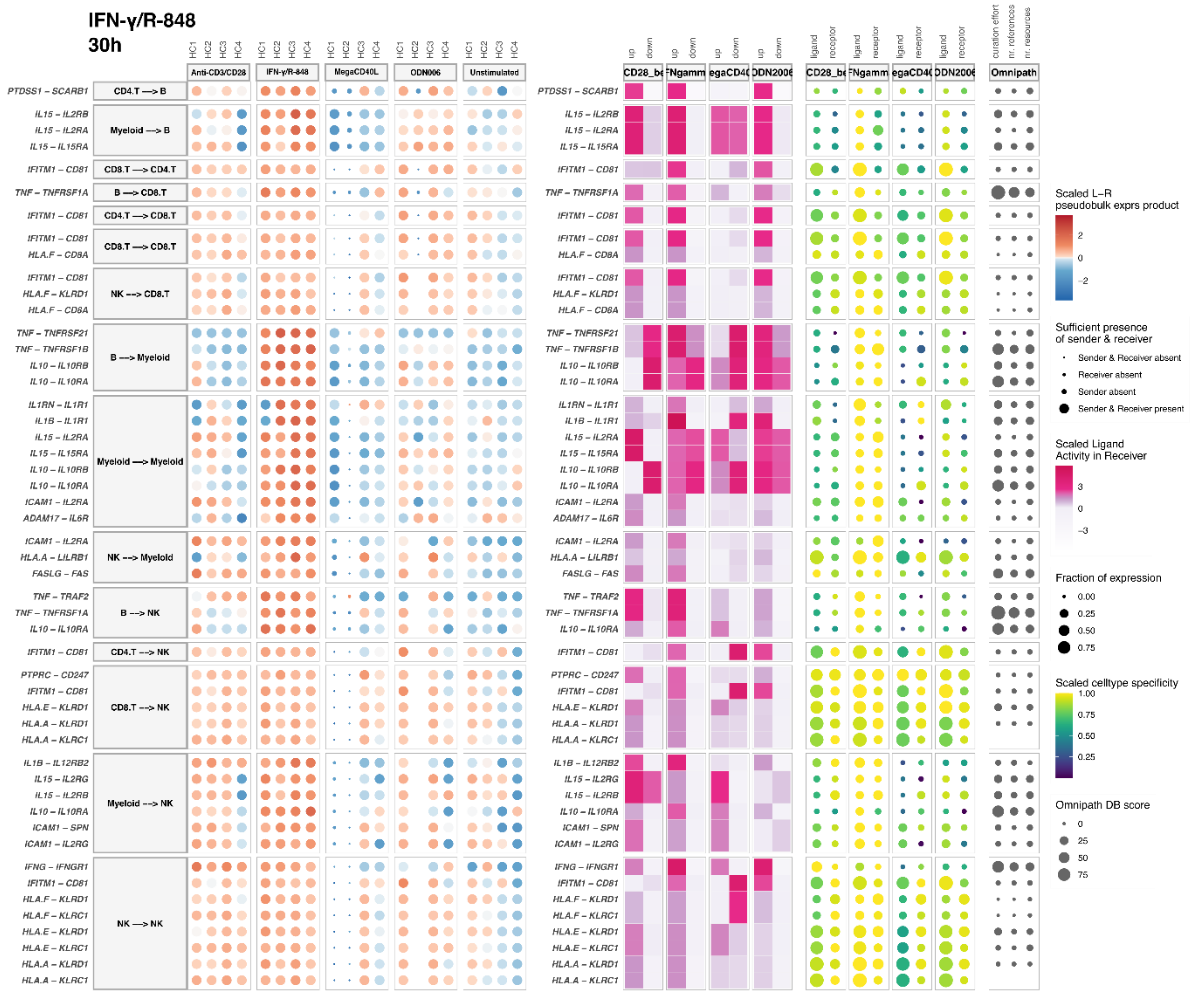


**Sup. Figure S18: Top 50 interactions observed in the cell-cell communication output of IFN-γ/R-848 after 30h of stimulation, with the unstimulated samples as reference.** Cell-cell interactions through specific ligand-receptor pairs are shown at the vertical axis. The dot plot at the left shows the scaled (pseudobulk) expression product of the ligand-receptor (LR) combination per sample (HC1-HC4, horizontal axis). The size of the dots represents the presence or absence (after filtering, see Materials&Methods) of the sender and receiver cell type within the individual samples. The tile plot in the middle shows the scaled ligand activity in the receiver cell type, split over up- and down-regulated target genes of the ligand-receptor pair (horizontal axis). Higher ligand activities (dark pink) suggest functional interaction with an effect on the downstream target gene regulation in the receiver cell. The colored dot plot at the right shows the cell type specificity of the ligand and receptors present in the interactions. The size of the dots indicates the expression fraction of the ligands and the receptors in the sender and receiver cells, respectively. The dot plot at the left represents the score of the interaction with the Omnipath database (DB) as reference.

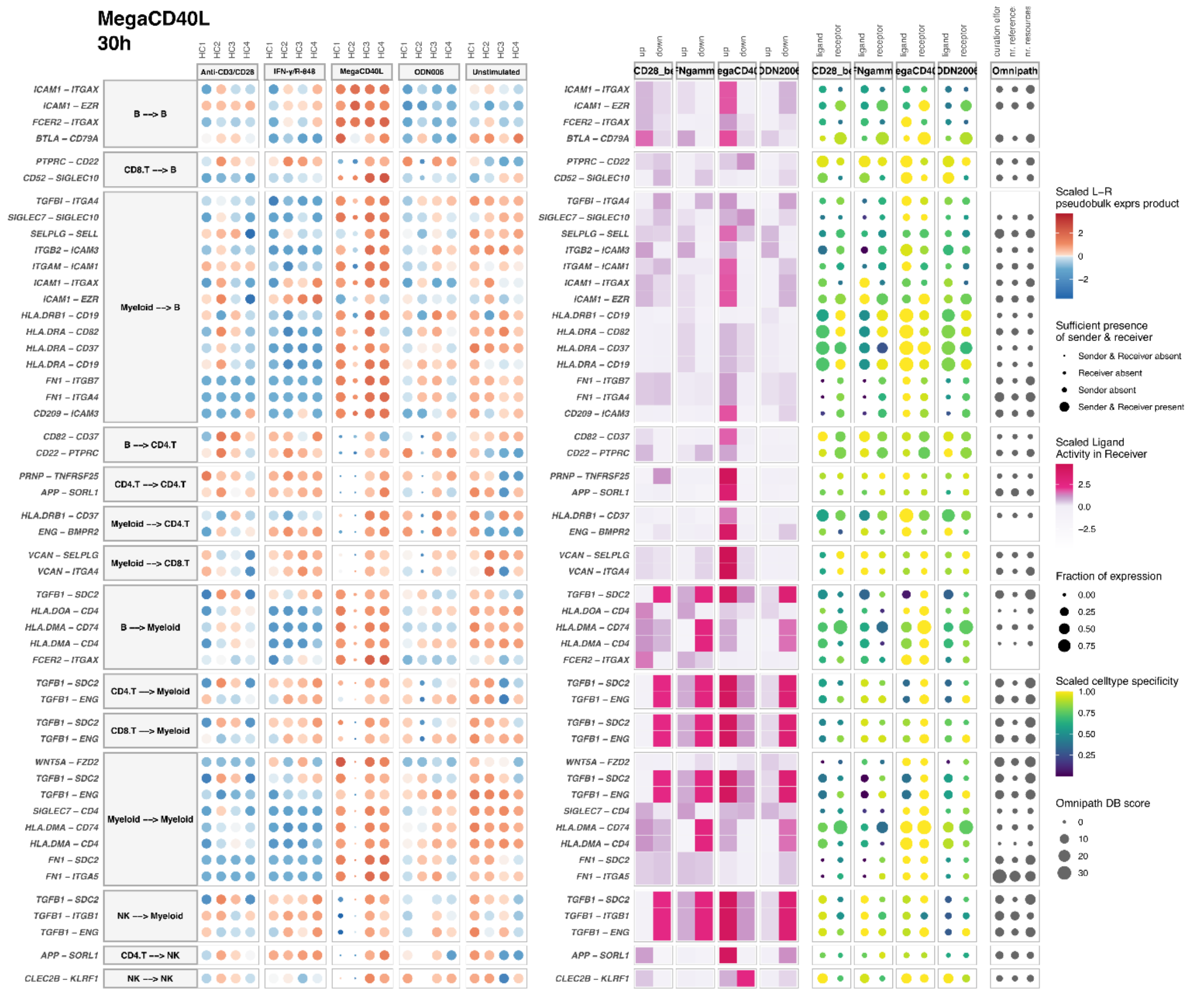


**Sup. Figure S19: Top 50 interactions observed in the cell-cell communication output of MegaCD40L after 30h of stimulation, with the unstimulated samples as reference.** Cell-cell interactions through specific ligand-receptor pairs are shown at the vertical axis. The dot plot at the left shows the scaled (pseudobulk) expression product of the ligand-receptor (LR) combination per sample (HC1-HC4, horizontal axis). The size of the dots represents the presence or absence (after filtering, see Materials&Methods) of the sender and receiver cell type within the individual samples. The tile plot in the middle shows the scaled ligand activity in the receiver cell type, split over up- and down-regulated target genes of the ligand-receptor pair (horizontal axis). Higher ligand activities (dark pink) suggest functional interaction with an effect on the downstream target gene regulation in the receiver cell. The colored dot plot at the right shows the cell type specificity of the ligand and receptors present in the interactions. The size of the dots indicates the expression fraction of the ligands and the receptors in the sender and receiver cells, respectively. The dot plot at the left represents the score of the interaction with the Omnipath database (DB) as reference.

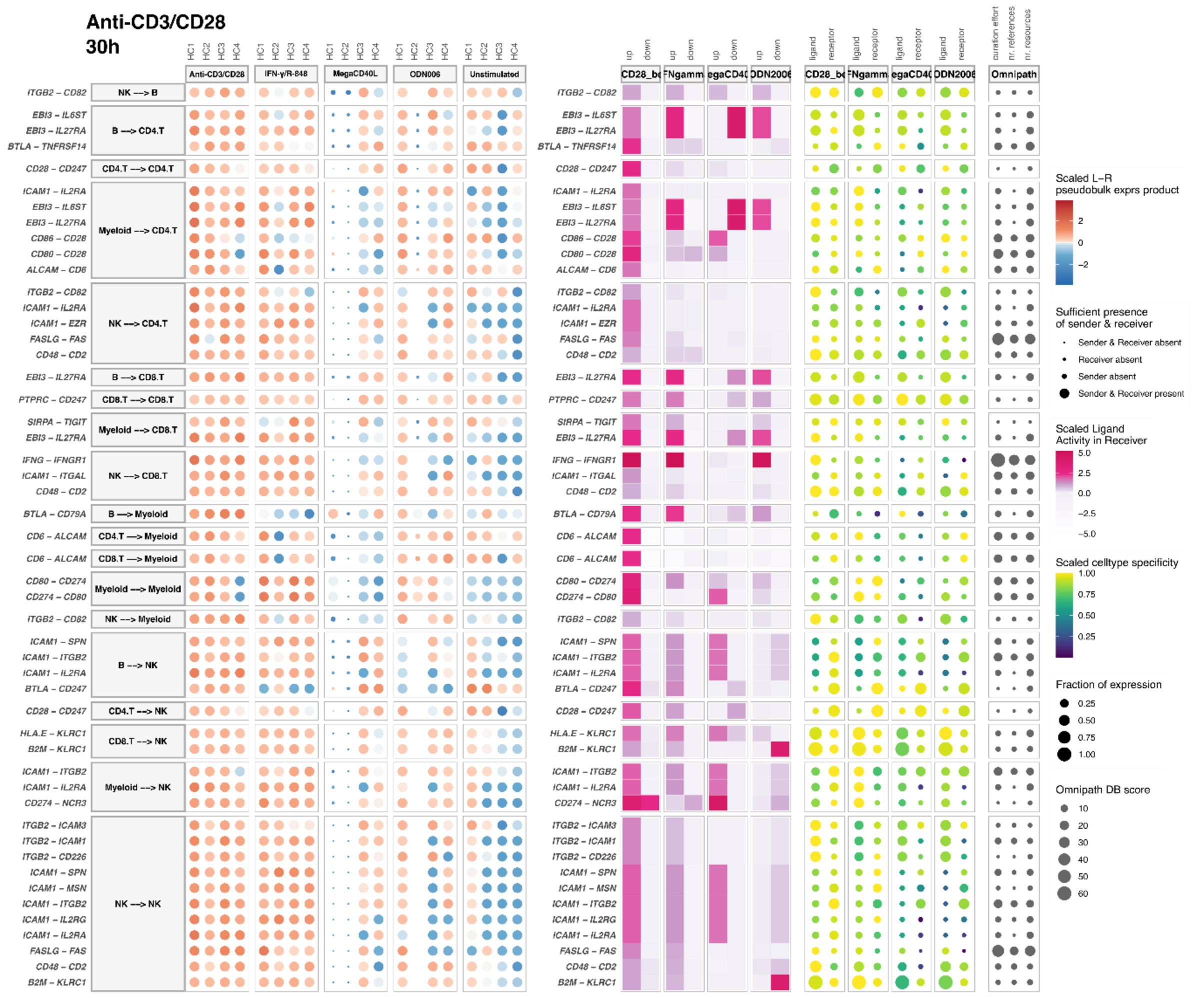

**Sup. Figure S20: Top 50 interactions observed in the cell-cell communication output of anti-CD3/CD28 after 30h of stimulation, with the unstimulated samples as reference.** Cell-cell interactions through specific ligand-receptor pairs are shown at the vertical axis. The dot plot at the left shows the scaled (pseudobulk) expression product of the ligand-receptor (LR) combination per sample (HC1-HC4, horizontal axis). The size of the dots represents the presence or absence (after filtering, see Materials&Methods) of the sender and receiver cell type within the individual samples. The tile plot in the middle shows the scaled ligand activity in the receiver cell type, split over up- and down-regulated target genes of the ligand-receptor pair (horizontal axis). Higher ligand activities (dark pink) suggest functional interaction with an effect on the downstream target gene regulation in the receiver cell. The colored dot plot at the right shows the cell type specificity of the ligand and receptors present in the interactions. The size of the dots indicates the expression fraction of the ligands and the receptors in the sender and receiver cells, respectively. The dot plot at the left represents the score of the interaction with the Omnipath database (DB) as reference.
