## Supplementary information on the Materials and Method section (referred to as Sup. Material D3 in the manuscript). for "Stimulation strategy matters: A B-cell focused single-cell transcriptomic study on *in vitro* stimulated immune cells"

### Supplementary Materials and methods

#### PBMC isolation and thawing

Peripheral blood mononuclear cells (PBMCs) were isolated within three hours of blood collection. Peripheral blood was collected in tubes with K_2_EDTA as anticoagulant. The blood was diluted 1:2 with Hank's Balanced Salt Solution (HBSS; Fisher Scientific; 24020117) and layered on Ficoll-Paque density separation medium (GE Healthcare; 17-1440-02). After gradient centrifugation (18 minutes – 800g – room temperature (RT)), PBMCs were washed with cold HBSS and manually counted using a Neubauer chamber with trypan blue staining. Cells were cryopreserved in 90% complete Roswell Park Memorial Institute medium (RPMI) and 10% dimethyl sulfoxide (DMSO; Sigma Aldrich; D2650). Complete RPMI included 90% RPMI containing GlutaMAX (ThermoFisher Scientific; 72400047), 1% penicillin/streptomycin (P/S, Gibco, ThermoFisher Scientific, 15140-122), 10% fetal calf serum (FCS; Sigma-Aldrich; F7524) and 0.001% β-mercaptoethanol (β-ME, Gibco, ThermoFisher Scientific, 31350-010). Vials were cooled in a controlled rate freezing container at -80°C before final storage at -150°C.

Frozen samples were thawed in a 37°C water bath and transferred to 15 mL pre-warmed (37°C) complete RPMI. PBMCs were centrifuged (7 minutes – 400g – RT), resuspended in 8 mL complete RPMI, and allowed to recover from thawing in a 5% CO_2_ incubator for 30 minutes at 37°C. After another centrifugation step, cells were resuspended in 15 mL complete RPMI, counted, and diluted to 10⁷ PBMCs/mL.

#### Flow cytometry workflow

##### FACS data analysis

The preprocessed files were subsampled, and three million cells, selected over the different samples, were aggregated into a single FCS file to generate a FlowSOM^1^ model with 100 clusters and 5 metaclusters based on CD45 and CD19. All preprocessed FCS files were mapped onto this FlowSOM model, and CD45^+^ cells (~85% of events after preprocessing but before debris removal) were extracted and saved into new FCS files. These files were used to generate a second FlowSOM model with 100 clusters and 15 metaclusters using markers CD45, CD19, CD27, CD69, CD80, CD83, CD86 and IgD (**Supl.** **Table S2** and **Supl. Figure S1A**). The metaclusters were then combined based on CD19 expression, resulting in two CD45+ subpopulations: CD19^+^ and CD19^-^. Cells within the CD19^+^ clusters were extracted, saved into new FCS files, and used to generate a final FlowSOM model with 100 clusters and 15 metaclusters based on CD19, CD27, CD69, CD80, CD83, CD86 and IgD expression, used as input for cell phenotype annotation.

Cell counts and mean fluorescence intensity (MFI) values per sample and B cell subset were extracted using the GetFeatures() function of the FlowSOM^1^ package. Fractions of activated (CD69^+^) and inactivated
(CD69^-^) B cell subsets (CD27^-^IgD^+^, CD27^+^IgD^+^ and CD27^+^IgD^-^) were calculated per sample (characterized by healthy donor, stimulatory condition and incubation time point) by dividing the cell counts per subsets by the total cells observed for that specific sample. MFI values of the surface markers of interest were normalised with respect to the unstimulated samples by dividing the MFI value of the surface marker of interest per sample by the mean MFI value of the unstimulated samples.

#### Single-cell RNA-seq experimental workflow

Stimulated PBMCs were tagged with cell hashing antibodies for sample multiplexing, enriched for B cells using Fluorescence-Activated Cell Sorting (FACS), and pooled equimolarly into six batches (three pools per incubation time point, **Supl. Table S3**) loading on a microfluidics device for scRNA-seq profiling.

#### Single-cell RNA-seq data-analysis workflow

###### Data integration

Objects originating from distinct pools over the two incubation time points were merged (Seurat::merge()) and afterwards split (Seurat::split()) at sample level using a splitting argument involving a combination of the healthy control identifier and stimulation condition (e.g. *HD1_MegaCD40L*). Data normalization, variable gene identification, and scaling of the splits were performed using Seurat::NormalizeData()*,* Seurat::FindVariableFeatures()*,* and Seurat::ScaleData()*,* respectively. Dimensionality reduction was performed (Seurat::RunPCA()) prior to the Canonical Correlation Analysis (CCA) integration step to consequently remove technical variability using Seurat::IntegrateLayers(…, method=CCAIntegration)^2^. Post-integration, cells were clustered using the default Louvain approach (Seurat::FindNeighbors(), Seurat::FindClusters(resolution=1.5)), Uniform Manifold Approximation and Projection (UMAP) were generated based on the CCA integration output (Seurat::RunUMAP(reduction=“integrated.cca”)), and all splits were joined into a final Seurat object, used for downstream analysis (Seurat::JoinLayers()).

###### Doublet detection

Doublet detection in the integrated Seurat object was performed using the scDblFinder package. First, the Seurat object was transformed to a SingleCellExperiment^3^ object (SingleCellExperiment:: as.SingleCellExperiment(assay=“RNA”)) prior to performing doublet detection using scDblFinder::scDblFinder(samples=“samplename”, clusters=“cca_clusters_rna”). After removing 12.7% of the 92,958 total events annotated as multiplets, a final dataset of 81,160 singlets was obtained (**Supl. Figure S4A**)^4^.

#### In-depth analysis of B-cell subsets

##### B-cell subset selection, integration and filtering

For 8h of stimulation, a total of 24 initial clusters were obtained. Cluster 17 (640 cells) and 22 (112 cells) were excluded from downstream data analysis due to a high number of remaining cells annotated as CD4+ T cells based on the Azimuth annotation and exclusively containing plasmablasts, respectively. For 30h of stimulation (26 clusters), the same reasoning was followed for the removal of cluster 12 (1,019 cells) and 24 (109 cells). Next, cluster 19 (696 cells) was removed due to a high percentage of mitochondrial DNA in comparison to the other clusters.

##### Generation and annotation of neighbourhoods

Differential abundance analysis using miloR^5,6^ was performed on the filtered B-cell subsets (8h and 30h of stimulation separately) by transforming the original B-cell Seurat objects to a SingleCellExperiment object (Seurat::as.SingleCellExperiment()) and afterwards to a Milo object (miloR::Milo()). Next, a k-nearest neighbour graph was built using miloR::buildGraph(…, k=50, d=20), neighbourhoods (Nhoods) were generated using miloR::makeNhoods(…, k=50, d=20, prop=0.15), the distance between Nhoods was calculated using miloR::calcNhoodDistance() and the final visualization of the Nhoods was generated using buildNhoodGraph(…, overlap = 5) based on the PCA dimensionality reduction embedding obtained after CCA integration. These Nhoods contain a unique set of highly similar cells, but it should be noted that one cell can be linked to multiple Nhoods. Contrasts for DA testing with miloR::testNhoods(…, fdr.weighting=‘graph-overlap’) were generated for every stimulation separately, in which the unstimulated cells were used as reference: e.g. ‘StimulationMegaCD40L – StimulationUnstimulated’. Nhood graphs were plotted using miloR::plotNhoodGraphDA() based on the CCA integrated UMAP embeddings.

##### Selection of differentially abundant (un)stimulated neighbourhoods

The tables below indicate the selection of Louvain clusters showing enrichment for highly differentially abundant Nhoods based on the generated beeswarm plots (**Table Sup. Mat. 1**, **Figure 3G-H** and **Sup. Figure S5**) as a first step in the Nhood selection process. The number of finally retained Nhoods, per stimulatory condition and incubation time point, is shown in **Table Sup. Mat. 2**.

Table Sup. Mat. 1: Clusters selected for extraction of highly differentially abundant Nhoods. Differentially abundant Nhoods are selected only if they were annotated as one of the original Louvain clusters listed below. This selection if condition and incubation time point specific.

| Condition | Clusters for 8h of stimulation | Clusters for 30h of stimulation |
| --- | --- | --- |
| Unstimulated | 0 – 9 – 11 – 13 | 2 – 4 |
| ODN2006 | 1 – 3 – 6 – 12 | 0 – 1 – 3 – 5 – 6 – 7 – 8 – 10 – 12 – 15 |
| IFN-γ/R-848 | 1 – 3 – 6 – 12 | 0 – 1 – 3 – 7 – 8 – 9 – 12 – 14 |
| MegaCD40L | 2 – 5 – 7 | 6 – 11 |
| Anti-CD3/CD28 | 2 – 5 – 7 | 0 – 1 – 3 – 5 – 6 – 7 – 9 – 11 – 13 |

Table Sup. Mat. 2: Number of retained Nhoods per stimulatory condition and incubation time point. From the 1.811 and 2.153 Nhoods called within the 8h and 30h stimulated B-cell subset, only a fraction per condition is retained by taking into account the two selection criteria as mentioned in the text.

| Condition | Retained Nhoods for 8h of stimulation | Retained Nhoods for 30h of stimulation |
| --- | --- | --- |
| Unstimulated | 104 | 71 |
| ODN2006 | 290 | 435 |
| IFN-γ/R-848 | 246 | 457 |
| MegaCD40L | 163 | 81 |
| Anti-CD3/CD28 | 136 | 351 |

##### Refinement of cell annotation related to stimulation status

The finally retained number of cells used as input for differentially expression analysis after stimulation status refinement is shown in the **Table Sup. Mat. 3**.

Table Sup. Mat. 3: Retained number of cells per condition and incubation time point.

| Condition | Retained cells for 8h of stimulation | Retained cells for 30h of stimulation |
| --- | --- | --- |
| Unstimulated | 3,165 | 1,627 |
| ODN2006 | 2,522 | 3,813 |
| IFN-γ/R-848 | 3,422 | 7,368 |
| MegaCD40L | 1,802 | 1,033 |
| Anti-CD3/CD28 | 2,643 | 4,055 |
| Total | 13,554 | 17,896 |

### References

1.	Van Gassen, S., Callebaut, B., Van Helden, M.J., Lambrecht, B.N., Demeester, P., Dhaene, T., and Saeys, Y. (2015). FlowSOM: Using self-organizing maps for visualization and interpretation of cytometry data. Cytometry A *87*, 636–645. https://doi.org/10.1002/cyto.a.22625.

2.	Hao, Y., Stuart, T., Kowalski, M.H., Choudhary, S., Hoffman, P., Hartman, A., Srivastava, A., Molla, G., Madad, S., Fernandez-Granda, C., et al. (2024). Dictionary learning for integrative, multimodal and scalable single-cell analysis. Nat Biotechnol *42*, 293–304. https://doi.org/10.1038/s41587-023-01767-y.

3.	Amezquita, R.A., Lun, A.T.L., Becht, E., Carey, V.J., Carpp, L.N., Geistlinger, L., Marini, F., Rue-Albrecht, K., Risso, D., Soneson, C., et al. (2020). Orchestrating single-cell analysis with Bioconductor. Nat Methods *17*, 137–145. https://doi.org/10.1038/s41592-019-0654-x.

4.	Germain, P.-L., Lun, A., Garcia Meixide, C., Macnair, W., and Robinson, M.D. (2022). Doublet identification in single-cell sequencing data using scDblFinder. F1000Res *10*, 979. https://doi.org/10.12688/f1000research.73600.2.

5.	Dann, E., Henderson, N.C., Teichmann, S.A., Morgan, M.D., and Marioni, J.C. (2022). Differential abundance testing on single-cell data using k-nearest neighbor graphs. Nat Biotechnol *40*, 245–253. https://doi.org/10.1038/s41587-021-01033-z.

6.	Morgan, D.M., and Dann, E. (2025). miloR: Differential neighbourhood abundance testing on a graph.
